# Neural Mechanisms Supporting the Relationship between Working Memory Capacity and Proactive Control

**DOI:** 10.1101/2025.05.09.653198

**Authors:** Rebecca Feldman, Maya Quale, Joset Etzel, Todd S. Braver

## Abstract

Recent prior work suggests a preferential relationship between working memory capacity (WMC) and proactive control, yet the neural mechanisms that support this relationship are still not well understood. We directly addressed this question by leveraging the Dual Mechanisms of Cognitive Control (DMCC) project, as it employed a fMRI neuroimaging design optimized to test for individual differences (sample N > 100), with task variants that independently assessed proactive and reactive control relative to baseline conditions. Behavioral analyses replicated prior work with the AX-CPT paradigm, in which a measure of target preparation based on contextual cues (the A-cue Bias index) was both reliably increased under task conditions encouraging proactive control and positively associated with WMC. Analyses of fMRI activity indicated that A-cue Bias was selectively linked to increased cue-related neural activity in left motor cortex (lMOT). Additionally, WMC was associated with increased cue-related activation in right dorsolateral prefrontal cortex (rDLPFC), even when statistically controlling for baseline and reactive conditions. The relationship between these two effects was supported by a latent path analysis, which suggested that the rDLPFC-lMOT circuit preferentially mediates the WMC-A-cue Bias relationship present under proactive task conditions. The results suggest this neural circuit may translate strategic task goals into active response preparation as a mechanism of proactive control. Individuals high in WMC may be better able to implement proactive task strategies when instructed via contextual cues. The sensitivity of the rDLPFC-lMOT circuit to individual differences suggest it as a potential target for cognitive enhancement.

## Introduction

Decades of research have gone into understanding the core factors and primary mechanisms that give rise to cognitive control, which reflects the ability to regulate, coordinate, and sequence thoughts and actions in accordance with internally maintained behavioral goals (Chiew & Braver, 2017; Egner, 2017; Miller & Cohen, 2001; Posner & Snyder, 1975; Stokes et al., 2017). Critically, cognitive control function – which has also been referred to somewhat interchangeably as executive control, executive function, or attentional control – is strongly impacted by individual differences (Braver et al., 2010; Friedman & Miyake, 2017; Kane & Engle, 2002). To better understand the source of these individual differences, more granular constructs have been invoked. In particular, working memory capacity (WMC), the ability to temporarily store, manipulate, and retrieve a limited amount of goal-relevant information, is one of the most well-studied individual differences constructs associated with cognitive control (Cowan, 2016). For example, individuals with higher WMC tend to have better attentional control, response interference resolution, and dual-task management (Belletier et al., 2019; M. A. Boudewyn et al., 2015; Burgess et al., 2011; Redick, 2014; Redick & Engle, 2011; Richmond et al., 2015; Stawarczyk et al., 2014; Wiemers & Redick, 2019). Nevertheless, this literature is not without controversy, as there have been disputes over the degree to which cognitive control reflects a unitary construct, and the degree to which an individual differences perspective can gain sufficient traction within this domain (Cohen, 2017; Friedman & Robbins, 2022; Gratton et al., 2018). Some of this work has focused on questions of measurement reliability, and if behavioral measures of cognitive control are suitable for individual differences research (Schuch et al., 2022; Von Bastian et al., 2020; Whitehead et al., 2018; Zorowitz & Niv, 2023)

An important perspective on this issue comes from the Dual Mechanisms of Control (DMC) framework (Braver, 2012). The DMC framework offers an account of cognitive control that explains variation across multiple levels – intra-individual (i.e., state- and context-related), individual differences, and between-population (e.g., age, clinical disorders) – by specifying a core temporal dimension of cognitive control that distinguishes between proactive and reactive modes. The proactive mode of control reflects the sustained active maintenance of goal-relevant information in anticipation of upcoming control demands, whereas reactive control reflects transient re-activation or retrieval of goal-relevant information, triggered by the detection of processing conflict or stimulus features associated with high control demands. The DMC framework suggests that uncontrolled variation related to the utilization of proactive vs. reactive control modes might complicate attempts to reliably measure cognitive control capacity. Moreover, the DMC framework postulates that distinct, testable dimensions of individual differences should be associated with proactive and reactive control (Braver, 2012), for example, that WMC is preferentially linked to proactive control mechanisms through top-down biasing of actively maintained goal-relevant information.

Prior work examining the relationship between proactive and reactive control, as well as the role of individual differences in the DMC framework, has frequently utilized the AX-CPT as a key experimental paradigm. Briefly, the AX-CPT measures cognitive control in terms of the active maintenance of contextual cues used to guide response preparation for subsequent probe items. Participants are instructed to make a target response if and only if an X-probe follows an A-cue (i.e., an AX trial). For all other trial types (BX, BY, and AY, where B and Y represent any letter other than A and X), a non-target response should be made. Accuracy and response time markers of cue- vs. probe-biased responding index proactive and reactive control, respectively. For example, the propensity to make a target response following an A-cue (i.e., independent of probe identity), which is termed the A-cue Bias, has been used as an index of proactive control (Braver et al., 2008; Gonthier, Braver, et al., 2016; Richmond et al., 2015). Conversely, the degree of reaction time interference on BX trials (relative to the low-control demand BY) trials, has been utilized as an index of reactive control (Braver et al., 2021; Tang et al., 2023).

In the DMC framework, proactive control relies on the ability to actively maintain task- and goal-relevant information. Since WMC is theoretically dependent on utilization of goal-relevant information as a means to regulate short-term storage, proactive control should also depend on WMC. In line with this, studies investigating the AX-CPT task have frequently utilized WMC as an individual difference measure to explain variation in cognitive control strategies and success, particularly in connection with proactive control (Belletier et al., 2019; M. A. Boudewyn et al., 2015; Stawarczyk et al., 2014). Indeed, individuals with higher WMC have been consistently shown to more frequently or strongly engage proactive control in comparison to lower WMC participants, with evidence that such findings extend to children as young as 5 (Gonthier et al., 2019; Redick, 2014; Redick & Engle, 2011; Richmond et al., 2015; Troller-Renfree et al., 2020; Wiemers & Redick, 2018). For example, individuals with higher WMC tend to make more cue-biased (AY) errors but fewer probe-biased (BX) errors, suggesting they are utilizing and/or maintaining the cue-related information in a proactive manner. This does not mean individuals with lower WMC are incapable of implementing proactive control; strategy training, practice, incentives, and providing more time can lead to proactive shifts in task performance intra-individually (Belletier et al., 2019; M. A. Boudewyn et al., 2015; Redick, 2014; Redick & Engle, 2011; Richmond et al., 2015; Rosales et al., 2022; Stawarczyk et al., 2014; Wiemers & Redick, 2018).

The prior work relating cognitive control in the AX-CPT to individual difference constructs such as WMC leads to an important question regarding when and how such individual differences come into play to modulate task strategies and performance. In particular, a key question is whether high WMC generally increases the tendency to spontaneously utilize proactive control in tasks such as the AX-CPT, or whether it primarily reflects the ability to implement this mode of control when required by task demands. A useful analogy is that of a stress test, used to index cardiovascular functioning. The role of the stress test is to elevate the participant’s heart rate, to reveal cardiovascular problems that would not be easily detected when the heart was at rest. Similarly, the role of WMC in proactive control can be revealed through experimental correlational designs, in which the demands on proactive control are systematically manipulated, and the effects of individual differences in WMC examined across the manipulations.

Experimental correlational designs have recently been employed to test for a selective role for WMC in proactive control (Lin et al., 2022; Rosales et al., 2022). In Lin et al. (2022), Bayesian modeling demonstrated a preferential positive association between WMC and proactive control indices within the AX-CPT during a proactive experimental condition, in which participants were explicitly instructed that utilizing proactive control would increase performance; conversely, such an association was not present at baseline, nor in conditions that encouraged reactive control. This finding indicates that WMC specifically influences the ability to flexibly implement proactive control when it is deemed beneficial (e.g., via an instructed strategy). If higher WMC was related to a general (or more obligatory) tendency to implement proactive control, it should have equivalently influenced increased proactive tendencies (i.e., A-cue Bias) under baseline and reactive conditions. In the context of the DMC theory, the findings support the notion that stable individual differences directly influence cognitive control.

In parallel to these behavioral findings, neuroimaging studies have studied mechanisms of proactive and reactive control within the AX-CPT (Braver et al., 2009; Gómez-Ariza et al., 2017; Paxton et al., 2008). Both fMRI and EEG studies have linked proactive control to cue-related activity in fronto-parietal regions, particularly the dorsolateral prefrontal cortex (DLPFC), whereas reactive control has been linked to probe-related activity in both the fronto-parietal and cingulo-opercular networks. Additional work has utilized within-subject manipulations—such as incentives and strategy training—to induce shifts from reactive to proactive control, and vice versa (Braver et al., 2009; Lopez-Garcia et al., 2016; Mäki-Marttunen et al., 2019b; Ryman et al., 2019). Such manipulations produced the expected behavioral effects in proactive and reactive control indices, as well as changes in the timing of DLPFC activation, which together suggest a shift from cue-based to probe-based engagement.

Despite the rich neuroimaging literature examining the role of DLPFC in cognitive control, and in the AX-CPT more specifically, a key gap remains: Is DLPFC a critical locus of WMC effects related to cognitive control? Neuroimaging studies are essential for uncovering the mechanisms underlying individual differences in WMC and their relationship to proactive and reactive control modes. For example, prior work by Burgess et al. (2011) demonstrated that shared individual variation in WMC and general fluid intelligence could be explained by the engagement of interference control mechanisms. This was evident not only through behavioral metrics, but also through increased interference-related activity within DLPFC and other fronto-parietal regions.

The current study provides a further investigation into the relationship between WMC and proactive control. We first tested for replication of the selective behavioral relationship identified in Lin et al. (2022) between the well-established proactive control index, A-cue Bias, and a composite working memory capacity (WMC) measure derived from two span tasks the OSPAN and SYMSPAN. This replication was conducted with a new sample of participants who performed the AX-CPT while undergoing fMRI scanning. Such a replication is critical, given that the Lin et al (2022) results were discrepant from another recent behavioral study addressing a similar question with different statistical methods (Rosales et al., 2022).

Critically, we next tested for reliable brain-behavior relationships linking WMC to proactive control utilizing an experimental correlational design and neuroimaging dataset large enough (N>100) to enable well-powered individual differences analyses. We focused our analysis on the brain activation during the cue period, isolating activity related to preparation, but prior to when the probe appeared and responding occurred. Specifically, we analyzed brain activation in pre-defined regions of interest (ROI), by dividing the brain into parcels (Schaefer 17 Network 400 parcellation). Each parcel was tested to identify those whose activity varied with cue type in a specific way (i.e. increased activation to A-cue selective to the proactive condition as well as A>B-cue activation). Each identified parcel was then tested for brain-behavior relationships, specifically, between A-cue-related activity in the proactive condition, and either the A-cue Bias index or WMC, with integration of the two measures culminating in a final latent path analysis. To preview the results, these analyses revealed that WMC is selectively linked to the proactive control A-cue Bias index, via a neural circuit involving the right DLPFC engaging (presumably) response preparation processes within the left motor cortex.

## Methods

The dataset analyzed here is part of the larger Dual Mechanisms of Cognitive Control (DMCC) project first described in Braver et al. (2021) and Etzel et al., (2022). Specific components of the DMCC dataset have also been the focus of additional previous reports (Freund et al., 2021, 2025a; Singh et al., 2022; Tang et al., 2023). Here we briefly describe aspects of the DMCC dataset most relevant for the current work; additional details regarding the broader project can be found in these other references.

### Participants

The full DMCC dataset contains 129 healthy young adults (age: mean = 31 years, *SD* = 6.1 years; female = 78, male = 39, Prefer not to answer = 1). Of these, 30 participants were members of a twin pair, and 53 participants also took part in the HCP Young Adult Study (Van Essen et al., 2012); for this latter group, the mapping of DMCC to HCP IDs can be found in the HCP ConnectomeDB (db.humanconnectome.org), under the title “DMCC (Dual Mechanisms of Cognitive Control) subject key”. Data from an additional 11 participants who were enrolled in the study were excluded, due to incomplete datasets, technical problems during task fMRI or anatomical MRI acquisition and/or unacceptable levels of motion or task performance. Self-reported demographic breakdowns are: White/Caucasian = 85, Black/African American = 17, Asian = 10, Native Hawaiian or Pacific Islander = 2, and more than one race = 4.

Of the 30 twin pairs, 27 were monozygotic pairs. Analyses of twin pair effects are reported in Supplemental Materials. Briefly, the WMC measures were highly correlated amongst twins, while the A-cue Bias measures were not. Cue-related activation in key brain regions of interest (lMOT and rDLPFC) were also not correlated among twin pairs. Consequently, primary analyses were conducted using all participants. Supplemental analyses examined the effect of only including unrelated participants (by randomly removing one member of each twin pair). None of the primary conclusions reported below were changed (see Supplementary Material, Section S5).

Before enrollment, all participants were screened and excluded for MRI safety contraindications. Eligibility criteria included: participant age between 18-45 years, no severe mental illness or neurological trauma, and with restricted drug/medication usage (screening form located at https://osf.io/6efv8/). All participants provided written consent and the study was approved by the Institutional Review Board at Washington University in St. Louis.

### Design and Procedure

#### Out-of-Scanner: WMC

All participants underwent an initial 1-2 hours out-of-scanner behavioral session. In this session, after consent and MRI screening forms, participants completed a set of self-report individual difference and cognitive ability measures (Etzel et al., 2022). In the current work we consider only the Operation Span and Symmetry Span tasks, which are briefly described below.

#### Out-of-Scanner: Operation Span Task

The Operation Span Task (OSPAN; Turner & Engle, 1989; Unsworth et al., 2005) was one of the two tasks used to assess WMC. The task involved remembering a sequence of letters while completing a series of math problems. Each trial consisted of a sequence of math-letter pairs, with set sizes ranging from three to seven. The task consisted of two trials of each set size for a total of 10 trials (note that in the dataset analyzed for Lin et al., 2022, three trials of each set size were performed). After each math trial, participants were asked to select the letters in the order presented. Each participant’s response time deadline was based on their average performance during practice trials. To motivate valid performance, participants were informed that their data would only be included if their math accuracy was at least 85%, but this was not actually applied as a formal exclusion criterion. WMC was calculated by summing the total number of correctly recalled letters over the entire task (i.e., partial span score).

#### Out-of-Scanner: Symmetry Span Task

The Symmetry Span Task (SYMSPAN; Unsworth et al., 2009), was the second measure of WMC. Similar to the OSPAN, in this task, participants were tasked with remembering a sequence of spatial locations, while making a series of left-right vertical symmetry judgments about shapes. Each trial consisted of a sequence of location-symmetry judgment pairs, with set sizes ranging from two to five squares. After each trial, participants were asked to select the spatial sequence of the locations within a 4×4 grid. The task consisted of two trials of each set size, for a total of 10 trials (unlike Lin et al., 2022, which gave three trials of each set size). Response-time deadline for the symmetry portion was also calculated based on average performance during practice trials. Similar to the OSPAN, participants were told their data would be used if their symmetry judgment accuracy was at least 85%, but this was not actually applied as a formal exclusion criterion. WMC was calculated with the partial span score approach, as the sum of correctly recalled locations across the entire task.

#### In-Scanner: AX-CPT

In the AX-CPT participants are visually presented with sequential letter pairs (cue-probe pairs) in each trial (see **Figure 1**). For all cues, participants were instructed to press the button with their right index finger. A target response (right middle finger press) was required when an X-probe was presented following an A-cue (AX trials). A nontarget response (right index finger) was required for other probe types (AY, BX, and BY, where B and Y indicated any letter other than A or X, respectively). Additionally, the task also included no-go trials, which required withholding the response to the probe; no-go trials were indicated by a digit (1-9) probe rather than a letter (Ang or Bng trials). Task trials consisted of the following events and durations (**Figure 1a**): trial onset flicker (300 msec), cue (500 msec), delay (4000 msec), probe border + probe (300 + 500 msec), inter-trial interval (either 1600, 2800, 4000 msec with equal probability). Each AX-CPT task block consisted of 24 trials: 8 AX, 8 BY, 2 AY, 2 BX, and 4 no-go, 2 with A-cue and 2 with B-cue with 72 trials per scan run, 144 trials per condition. To separate task blocks, participants were presented with a 30 second fixation cross both prior to and following each task block, during which they were instructed to remain awake and as still as possible.

**Figure 1:**
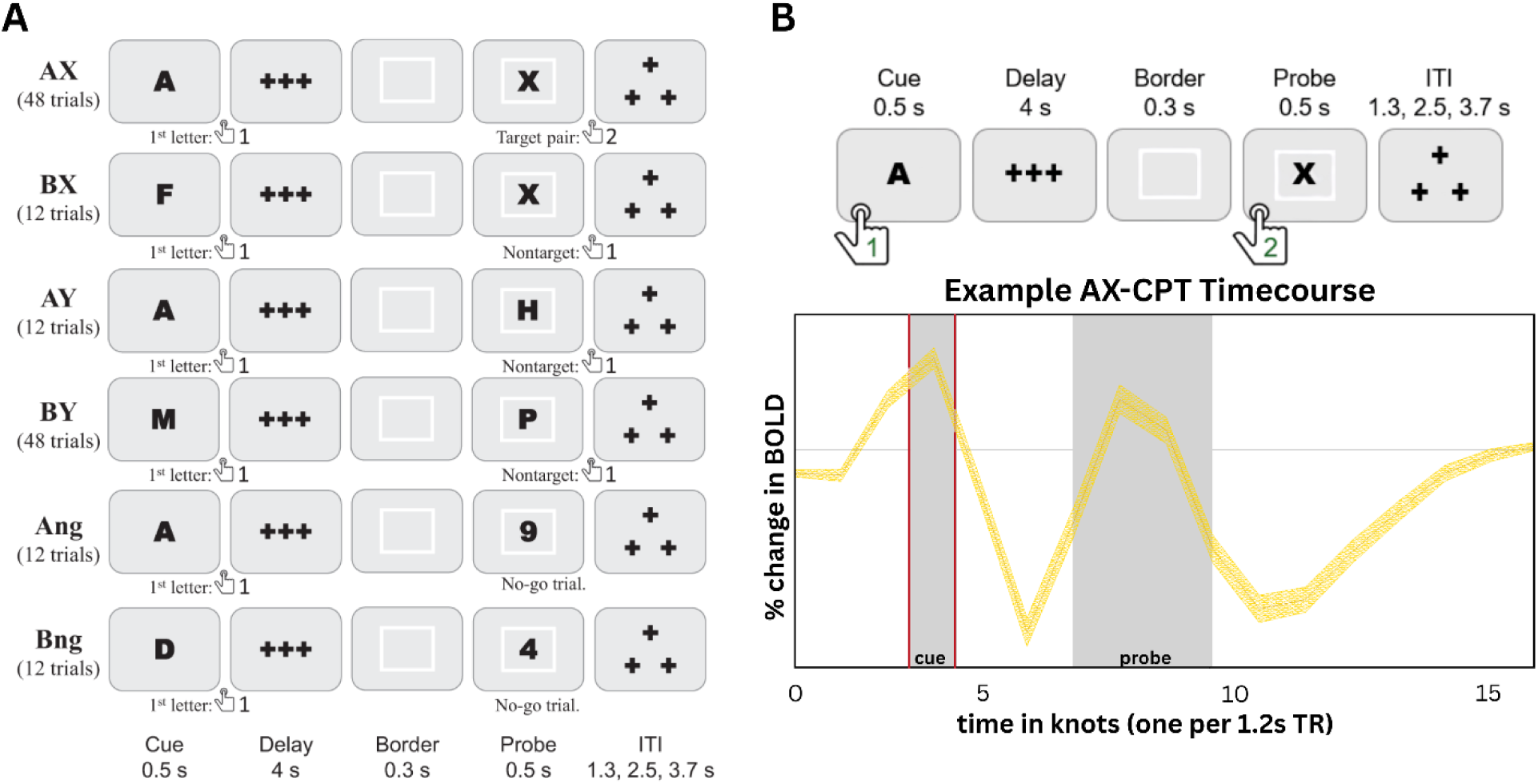
**a)** Shows the DMC AX-CPT task variant trial types, their frequency, and the required response. Note that the reactive condition (in which the border color and probe letter location varies) is not shown, b) Shows an example of how the cue and probe periods map onto observed GLM timccourscs. The particular timecourse shows A-cue activation, averaged across all conditions, of a representative parcel in the visual cortex. The peak activation periods associated with the cue and probe stimuli are highlighted in the shaded windows. The shaded cue period (TR 4) was the primary focus of all neuroimaging activation analyses

As indicated by the trial type frequencies, a key feature of the task is that X-probes are most likely to follow A-cues, thus leading to both cue-based and probe-based target expectancy biases. As a consequence, the tendency to make a target response following an A-cue can be used to index proactive control (as the A-cue Bias), whereas the interference caused by the X-probe on BX trials can be used to index reactive control. A second important aspect of the design is that A and B cues occurred with equal frequency, thus eliminating confounds present in prior AX-CPT designs (Gonthier, Macnamara, et al., 2016; Richmond et al., 2015; Rosales et al., 2022). Finally, the inclusion of no-go trials increased response uncertainty and decreased the overall predictive utility of context information. As such, the no-go AX-CPT variant decreases the overall proactive control bias typically seen in healthy young adults, which also provides a more sensitive baseline for detecting changes in control mode in the proactive and reactive conditions (Gonthier, Macnamara, et al., 2016).

All participants performed the AX-CPT task in three different conditions – Baseline, Proactive, and Reactive – each in a different scanning session with Baseline always being the first condition and Proactive and Reactive orders counterbalanced across participants (hereafter when explicitly referring to these experimental task conditions, we capitalize Baseline, Proactive and Reactive). The Proactive and Reactive conditions were identical to Baseline, with the following exceptions. In the Proactive condition of the task, participants received explicit strategy instructions regarding the cue-probe probability relationship. Specifically, they were instructed that following an A-cue, the X-probe would most likely occur and that they should utilize this information to begin preparing a target response following an A-cue during the delay. In the Reactive condition, probe stimuli were presented in different spatial locations on the screen, with the less frequent and high-conflict trials (AY, BX, no-go) appearing in the lower half, while more frequent low-conflict trials (AX, BY) appeared in the upper half. Additionally, the square border that appeared immediately (300 msec) before probe onset was a red color for high-conflict probes and a white color for low-conflict probes. This association between probe border and location and high vs. low conflict was not made explicit to participants, and so had to be learned through experience. However, once learned, this additional probe-related information could be utilized to reactively adjust to the presence of conflict-related interference. In the Baseline and Proactive conditions, all probe stimuli appeared in the center of the screen with a white border.

The AX-CPT was programmed and presented to participants using E-prime software (Version 2.0, Psychology Software Tools). Scripts can be requested at https://sites.wustl.edu/dualmechanisms/request-form/. Additional details regarding task stimuli and procedures can be found in the OSF repository (https://osf.io/xfe32/).

#### Imaging Session

Participants completed three separate imaging sessions in which they completed the DMCC task battery under different conditions: Baseline, Proactive, and Reactive, with the order of Proactive and Reactive counterbalanced across participants (for all participants the Baseline condition was performed first). Each imaging session lasted approximately 3 hours, began with task instruction and practice, and included two scanning runs of each task. Each 12-min AX-CPT task run was comprised of three task blocks, each preceded and followed by 30 seconds of fixation (a mixed block/event-related format Petersen & Dubis, 2012; Visscher et al., 2003).Trial duration and intertrial intervals were timed to synchronize with the scanner pulses to facilitate estimation of event-related responses. A description of the full DMCC protocol and all task variants is provided in Braver et al. (2021) and Etzel et al. (2022).

Briefly, neuroimaging data were acquired on a 3T Siemens Prisma with a 32-channel head coil in the East Building MR Facility of the Washington University Medical Center (protocol sheets can be found at https://osf.io/tbhfg). At the first session, high-resolution T1-weighted MPRAGE (0.8mm isotropic voxels; 2.4 s TR, 0.0222 s TE, 1 s TI, 8* flip angle) and T2-weighted (0.8mm isotropic voxels; 3.2 s TR, 0.563 s TE, 120* flip angle) anatomical scans were collected. Functional scans were acquired with CMRR multiband sequences (factor 4), without in-plane acceleration (iPat=none), for 2.4 mm isotropic voxels, 1.2 s TR (TE 33 msec, flip angle 53*). In each session two scanning runs of each task were collected, the first with an Anterior to Posterior (AP) encoding, immediately followed by a Posterior to Anterior (PA) run.

### Data Preprocessing and Analysis

#### Behavioral Analyses

For the AX-CPT task, the primary behavioral index of interest was the A-cue Bias, which prior work has suggested to be a reliable and selective metric of proactive control (Gonthier et al, 2016) and WMC (Richmond et al., 2015). The A-cue Bias metric indicates the overall bias to make a target (over non-target) response following an A-cue. This behavioral metric was chosen because it is only influenced by the cue, thus paralleling our neural metric. Following Lin et al (2022), the A-cue Bias was estimated using a single-trial logistic regression approach, combined with Hierarchical Bayesian Modeling (HBM), using the brms package in R (Bürkner, 2017). In particular, A-cue Bias was first modeled as the log-odds of making a target response to the probe within each session separately using the following regression model (see Supplementary Material, Section S1.1):

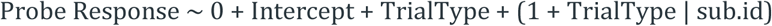

The resulting individual subject-level coefficients for A-cue Bias (the intercept term) were stored for each participant and used in subsequent analyses. Note that this trial-level formulation of the A-cue Bias is somewhat different than the traditional A-cue Bias measure, which is based on participant-wise accuracy scores and calculated as the log-linear correction of hits (AX) and false alarms (AY):

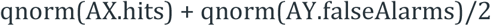

The trial-level approach was chosen for A-cue Bias calculation because it avoids the traditional limitations of summary or aggregate scores by taking into account both trial and subject-level variability (Lin et al., 2022; Rouder & Haaf, 2021). Nevertheless, the two A-cue Bias measures are comparable and highly correlated (Pro ρ=0.925, Bas ρ=0.929, Rea ρ=0.807; Supplementary Material, Section S1.2b).

Hierarchical Bayesian analyses were conducted to test for the effect of condition (e.g., Proactive vs. Baseline) and their interaction with WMC. For example, to replicate analyses conducted in Lin et al (2022), we used the following regression model:

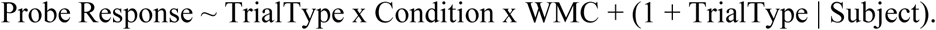

The key effect of interest was the Condition x WMC interaction, specifically the coefficient for the Condition:Proactive x WMC effect (the differential effect of WMC in the Proactive condition). Each estimated model was run with 4 Monte Carlo chains, each containing 2,000 sample iterations and 1,000 warm-up iterations, with warm-up iterations discarded. For each parameter estimate, we report the mean, standard deviation, and 95% credible interval (CI; quantile-based equal-tailed interval) of the posterior distribution, exponentiated log-odds to odds, as well as the R-hat and ESS (effective sample sizes) values (Supplementary Material, Section 1.3).

### fMRI Data

Preprocessing of fMRI data was performed with fMRIPrep 1.3.2 (RRID:SCR_016216; Esteban et al., 2019; Markiewicz et al., 2025), which is based on Nipype 1.1.9 (RRID:SCR_002502; Esteban et al., 2025; Gorgolewski et al., 2011). The full description can be found in the “Preprocessing” section of Etzel et. al (2022). Analyses were conducted in R version by reviewing temporal mean and standard deviation images (Etzel, 2023). Frames with more than 0.9 mm FD were censored, and runs dropped if more than 20% of its frames were censored.

Task fMRI data were analyzed using mixed block/event-related general linear model (GLM) estimation in AFNI (Cox, 1996). For each participant and condition (Baseline, Proactive, Reactive), AFNI GLM procedures (3dDeconvolve, 3dREMLfit) were applied whole brain. Briefly, separate task regressors modeled each trial type of interest (i.e., AX, AY, BX, BY, Ang, Bng) with piecewise linear spline deconvolution (TENTzero; also known as finite impulse response modeling FIR), resulting in event-related activation estimated as beta coefficient timecourses at each TR following trial onset (See Supplementary Material; Section 2.1). Regressors of no interest were included for the six realignment parameters, polynomial trends, and block onsets and offsets. Block regressors were included to model sustained activity but are not considered here. For analysis, the estimated activity at each knot and voxel was averaged within each of the cortical-spanning parcels from the independent Schaefer atlas (7 Network 400 Parcel version Schaefer et al., 2018), resulting in parcel-wise activity timecourses such as that shown in **Figure 1b**.

The key focus of our analysis was on A-cue-related activation (though A-cue > B-cue activation is considered in initial steps), so event-related activity was extracted at the 1-TR peak following cue onset, knot 4, allowing for the ∼5-second hemodynamic lag (see **Figure 1b** for a visualization of this cue period in a canonical event-related timecourse. These participant-level parcel estimates of A-cue activation for each condition became the primary dependent measure of interest included in subsequent analyses testing for effects of proactive control, brain-behavior associations with A-cue Bias, and individual difference relationships with WMC. Analyses were completed in R version 4.4.0 (2024-04-24) with WRS (Wilcox, 2012) and DescTools package functions for robust descriptive statistics. Trimmed (at .1) means and standard errors are reported unless otherwise specified.

## Results

The following overviews the key components of our analyses, reported below. We first conducted a behavioral analysis to demonstrate that trial-wise behavior in the form of A-cue Bias was linked to WMC specifically in the Proactive condition, replicating the approach used in Lin et. al (2022). In our initial fMRI analysis, we utilized a conjunction approach to identify candidate ROIs based on cue-related fMRI activity, focusing on activation that was selectively increased during A-cue presentation. Then, in these identified candidate ROIs, we tested for a positive relationship between the A-cue Bias and cue activity. Specifically, via correlation and within-subject mediation analyses, we identified regions in which higher A-cue Bias during the Proactive condition (relative to Baseline) is associated with higher cue-related activity in this condition. Further, we tested for a correlational relationship WMC and A-cue related activity in these candidate parcels. Finally, we conducted a latent path analysis to formally link the A-cue Bias and WMC findings. Supplementary materials, as well as of the data and R code to conduct and replicate the analyses reported below, are available on OSF, except for twin-related information (https://osf.io/d4k8n/). Twin-related data can be requested through the HCP.

### WMC effects on behavioral metrics of proactive control

Following Lin and colleagues (2022), we observed a high correlation between OSPAN and SYMSPAN scores (ρ=0.55, p<0.001; see Table 1 and Supplementary1, S1.1 for calculations). Consequently, they were combined into a composite WMC index in the same way as Lin et al., (2022), by z-score normalizing (relative to the group mean and standard deviation) and summing the two span scores.

**Table 1.**
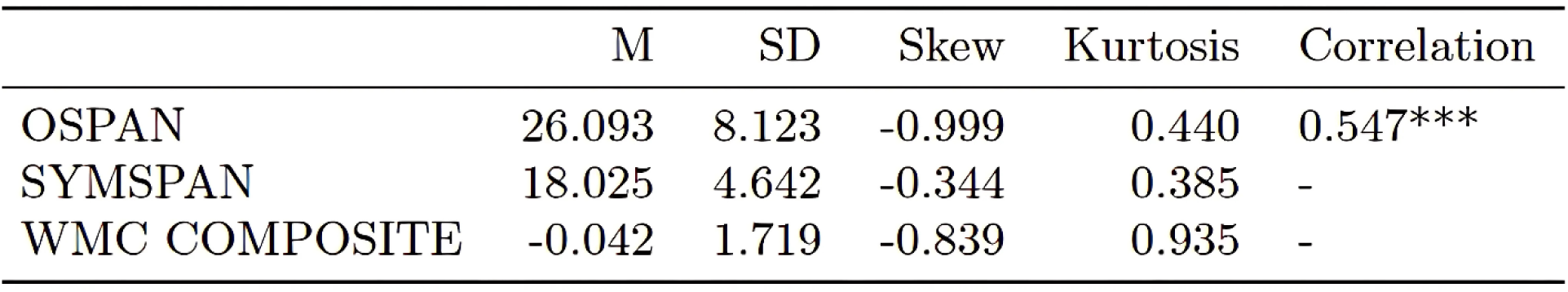
WMC composite scores were calculated by z-score normalizing (relative to the group mean and standard deviation) and summing the OSPAN and SYMSPAN scores, which were highly correlated with each other r=0.55, p<0.001.

In terms of zero-order correlations, WMC was significantly associated with A-cue Bias in both the Baseline (r=0.19, p = 0.04) and Proactive conditions (r=0.23, p=0.01; see **Figure 2a**), but not the Reactive condition (p=0.81). Further, the A-cue Bias index was significantly increased in the Proactive condition relative to both Baseline (t=19.28, df=113, p<0.001) and Reactive (t=20.23, df=115, p<0.001), indicating its sensitivity as a behavioral metric of proactive control (Lin et al., 2022). More expanded analyses of the behavioral data, including full descriptive statistics and details regarding the estimation of the trial-level A-cue Bias metric, are presented in Supplementary Material, Section S1.2a and S1.2b.

**Figure 2:**
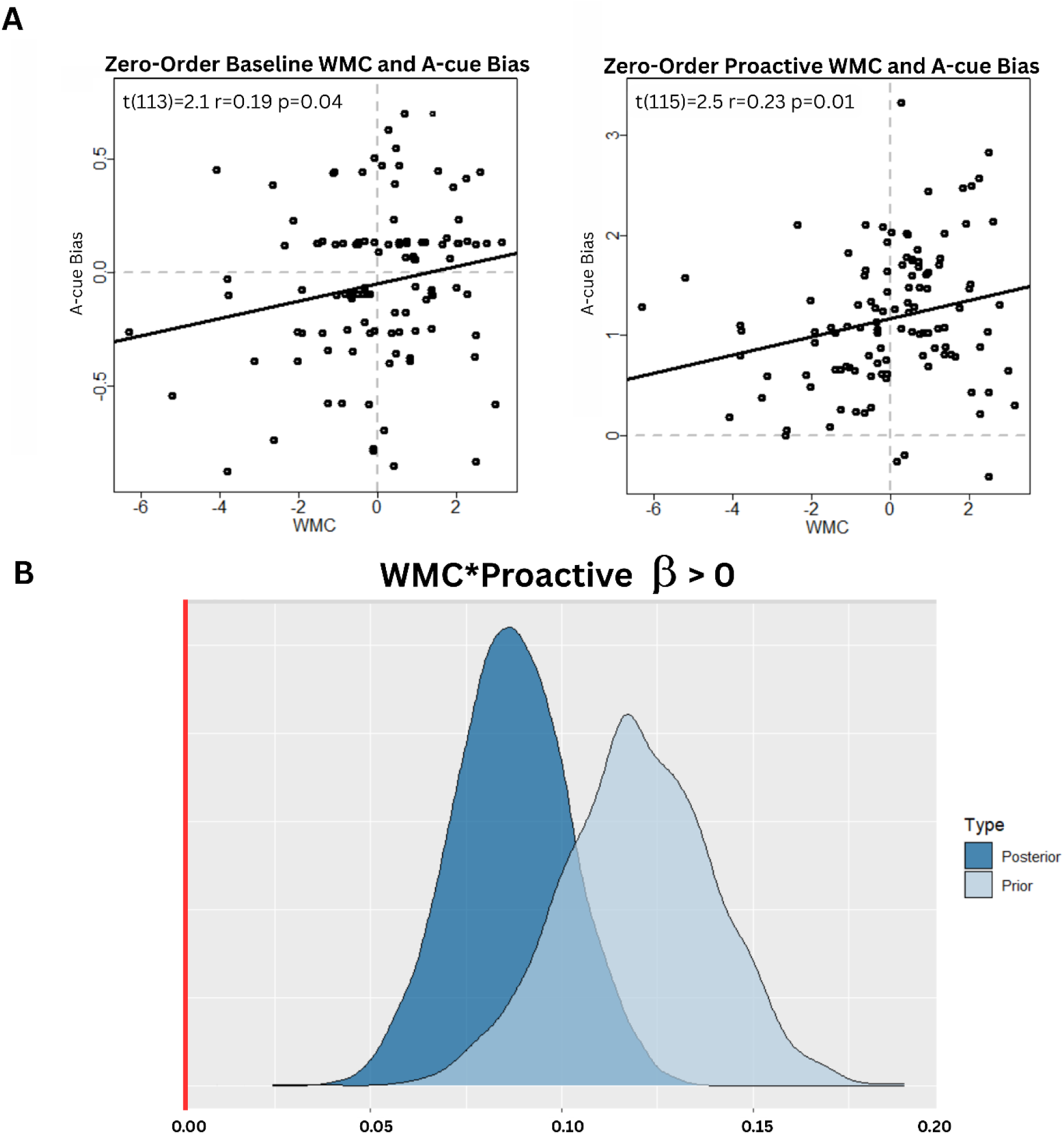
**a)** zero-order correlation between WMC and A-cue Bias in both Baseline and Proactive sessions. This correlation was not significant in Reactive, **b)** Prior and posterior distributions for the WMC x Proactive effect estimated from hierarchical Bayesian modeling. The prior distribution (light blue) reflects parameter estimates from the Lin et al., 2022 analyses. These estimates were entered as informed priors to derive posterior distributions to see the degree to which the results replicated. The overlap between the two distributions indicates that the WMC x Proactive effect is quite stable across datasets, and clearly shifted away from zero (red vertical bar), indicating a highly reliable effect.

We next used hierarchical Bayesian linear regression to conduct a more rigorous test of the hypothesis that the WMC – A-cue Bias relationship could be differentiated among the Proactive, Baseline, and Reactive conditions (Table 2; see Supplementary Material, Section S1.3 for more details). Although higher WMC was associated with a generally higher likelihood of making a target response to A-cues, there was an additional positive relationship that was unique to the Proactive condition (β= 0.11, SD= 0.01, 95% CI[0.01, 0.09]*, Eb=Inf^2,^ PD=100%), as no such effect was found in the Baseline condition (β=-0.02, SD=0.02, 95% CI[-0.06,0.01]).

**Table 2.**
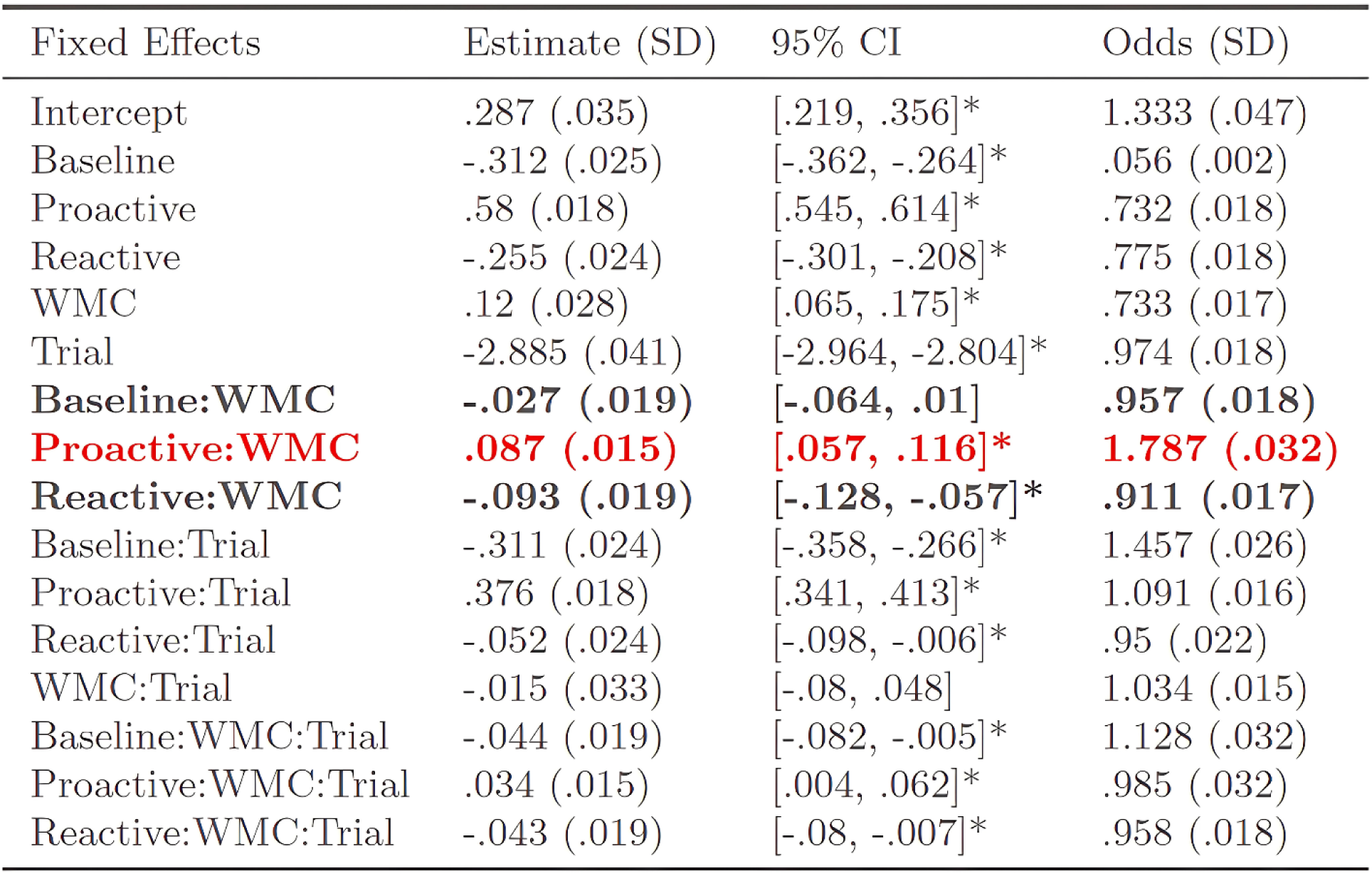
Results from the A-cue Bias replication analysis. Full formula: Probe Response ∼ TrialType X WMC X Session + (1 + trial type | subject). Posteriors from the online 2020 online mTURK DMC dataset were used as informed priors, which in turn were informed by posterior estimates as informed priors from the 2019 online mTURK DMC dataset. Fixed effects with * indicate a CI that does not include 0. The critical Proactive x WMC effect is highlighted in red font, while the contrasting Baseline x WMC and Reactive X WMC effects are shown in bold font.

| Fixed Effects | Estimate (SD) | 95% CI | Odds (SD) |
| --- | --- | --- | --- |
| Intercept | .287 (.035) | [.219, .356]* | 1.333 (.047) |
| Baseline | -.312 (.025) | [-.362, -.264]* | .056 (.002) |
| Proactive | .58 (.018) | [.545, .614]* | .732 (.018) |
| Reactive | -.255 (.024) | [-.301, -.208]* | .775 (.018) |
| WMC | .12 (.028) | [.065, .175]* | .733 (.017) |
| Trial | -2.885 (.041) | [-2.964, -2.804]* | .974 (.018) |
| **Baseline:WMC** | **-.027 (.019)** | **[-.064, .01]** | **.957 (.018)** |
| **Proactive:WMC** | **.087 (.015)** | **[.057, .116]\*** | **1.787 (.032)** |
| **Reactive:WMC** | **-.093 (.019)** | **[-.128, -.057]\*** | **.911 (.017)** |
| Baseline:Trial | -.311 (.024) | [-.358, -.266]* | 1.457 (.026) |
| Proactive:Trial | .376 (.018) | [.341, .413]* | 1.091 (.016) |
| Reactive:Trial | -.052 (.024) | [-.098, -.006]* | .95 (.022) |
| WMC:Trial | -.015 (.033) | [-.08, .048] | 1.034 (.015) |
| Baseline:WMC:Trial | -.044 (.019) | [-.082, -.005]* | 1.128 (.032) |
| Proactive:WMC:Trial | .034 (.015) | [.004, .062]* | .985 (.032) |
| Reactive:WMC:Trial | -.043 (.019) | [-.08, -.007]* | .958 (.018) |

Conversely, WMC was associated with *lower* log-odds of making a target response following an A-cue in the Reactive condition (β=-0.11, SD=0.01, 95% CI[-0.14,-0.07]*). In terms of statistical inference within the Bayesian framework, the PD (probability of direction) and ER (evidence ratio) can be used. Here, the interaction effect of Proactive condition and WMC has an ER approaching infinity, and PD of 100%, which together indicate extremely strong evidence in favor of this effect.

A key advantage of the Bayesian framework is that it allows for a formal test of replication, or conversely, a shift of the parameter estimate, by comparing the posterior parameter estimate to a prior taken from prior work, using the Savage-Dickey ratio (SDR). For interpretation, with SDR > 1, it means that the prior provides a good estimate of the posterior, whereas SDR < 1 means that the posterior has shifted away from the prior (Dickey, 1971; Kass & Raftery, 1995). Here, we take as a prior the estimate of the WMC x Proactive interaction estimate for A-cue Bias computed across two datasets of comparable size to the current DMCC neuroimaging sample, but taken from on-line behavioral studies (Lin et al., 2022). When comparing the posterior to the prior estimate taken from Lin et al (2022), we observe an SDR = 1.33, which indicates that the WMC x Proactive interaction effect is quite consistent with that estimated from the previous datasets (see **Figure 2b**). Moreover, the posterior estimate clearly has the full distribution mass shifted away from zero, indicating a decisive effect, in which WMC is differentially associated with the A-cue Bias index of proactive control, and preferentially in the Proactive condition.

### Cue-related activation in the AX-CPT Proactive condition

So far, we have demonstrated that stable individual differences in WMC uniquely interacted with the Proactive condition, such that increased WMC increased the log-likelihood of making a target response following an A-cue (i.e., the A-cue Bias index, modeled as a trial-by-trial effect), replicating Lin et al., (2022). Next, we examined the fMRI data, first looking for brain areas whose task-related activity suggests they change with AX-CPT cue type and control mode, and then for relationships between the brain and behavior.

To investigate the neural mechanisms supporting the relationship between A-cue Bias and WMC, we first identified parcels showing cue-related activity particularly on A-cue trials (i.e. mean activation across AX, AY and Ang trials), and that was selectively increased in the Proactive condition, given that this neural metric should be most closely linked with A-cue Bias (i.e., this index assesses behavioral tendencies specifically on A-cue trials). We calculated the fMRI activity in each of the 400 Schaefer parcels by estimating activation coefficients at distinct knots by GLMs, as described in the previous sections and illustrated in **Figures 1 and 3b**. We also utilized A-cue > B-cue activity as a contrast of interest, given the hypothesis that the explicit strategy instructions and training provided in the Proactive condition would increase the relative salience of A-cues for engaging advance preparation. A whole-brain conjunction approach was employed, selecting candidate parcels that satisfied each of the following conditions: 1) significant A-cue activity in Proactive (i.e., activation > 0); 2) A-cue > B-cue activity increased in Proactive relative to Baseline (i.e., [A-cue - B-cue]Proactive – [A-cue – B-cue]Baseline > 0); 3) A-cue > B-cue activity increased in Proactive relative to Reactive (i.e., [A-cue - B-cue]Proactive – [A-cue – B-cue]Reactive > 0; see Supplementary Materials, Section S2.2 for a step by step visualization of these tests). Because of our interest in utilizing this as a first-stage analysis to identify potential candidate ROIs, we were more concerned with false negatives than false positives, and so implemented a liberal approach to the conjunction, thresholding parcels for inclusion with uncorrected p-values (p < .05) via 1-tailed t-tests. This analysis yielded 40 candidate parcels, in a broadly distributed bilateral and primarily fronto-parietal network (**Figure 3a**; see Supplementary Material Section S2.3). As can be seen, for these ROIs, the estimated activation timecourse (shown for A- and B-cue trial types) exhibits prominent activation during both cue and probe periods, along with differentiation between A-cue and B-cue activity, and among the three task conditions (**Figure 3b**).

**Figure 3:**
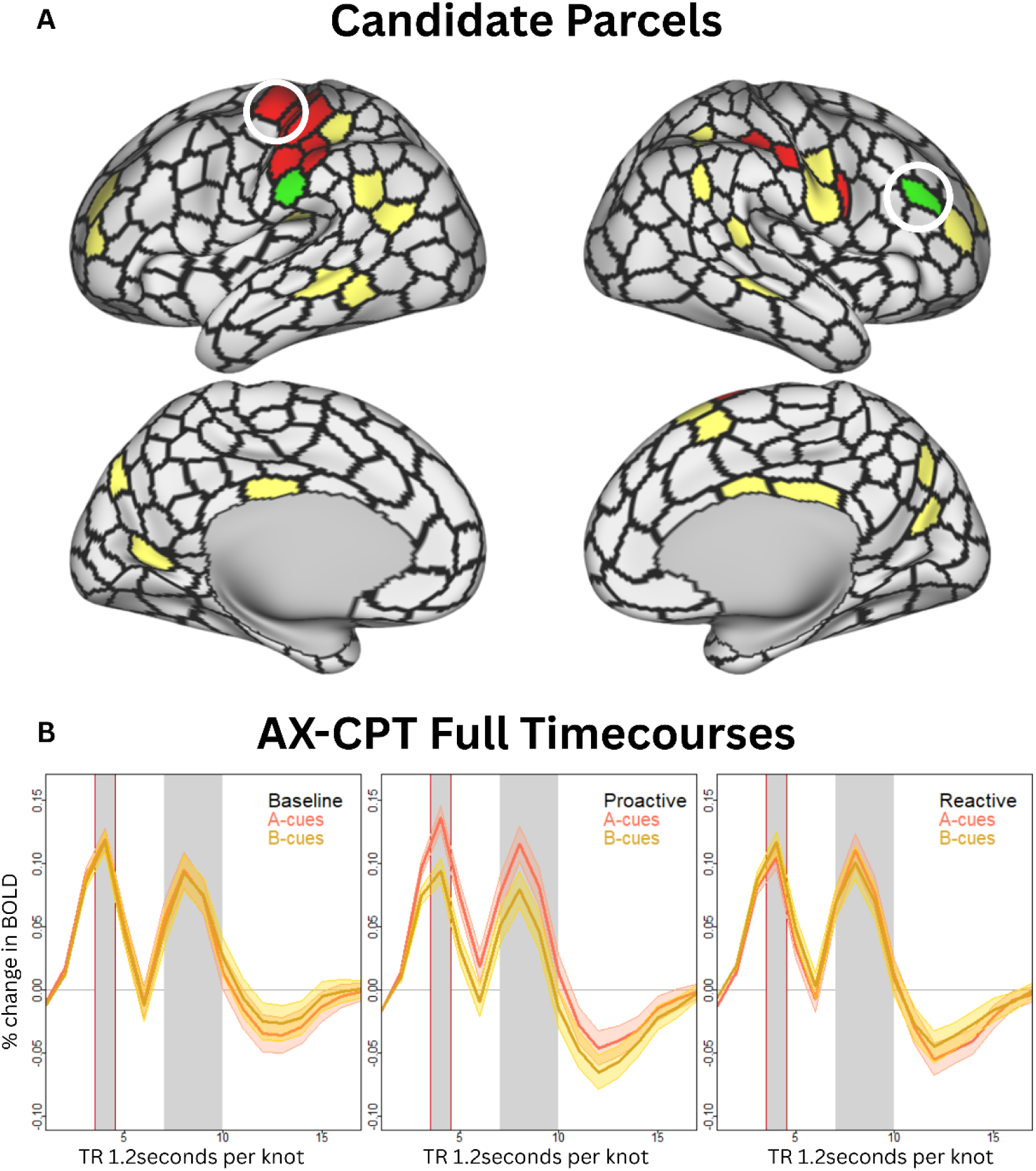
**a)** Conjunction analysis parcels in yellow that had A-cue activation > 0, more A>B-cue activation in Proactive relative to Baseline and Reactive sessions. Parcels in red were also significantly correlated with. A-cue Bias and those in green were correlated with WMC. The circled red parcel contains 2 1MOT parcels combined and used for subsequent analyses, while the circled green parcel shows the rDLPFC selectively associated with WMC in the Proactive condition, **b)** The full timecourse illustrates event-related activation (averaged across the 40 candidate ROIs) for A and B-cues respectively. Timecourses were created with robust means and standard errors (SEM) trimmed at 0.1.

### Brain-behavior relationships in Proactive control

Having identified the 40 candidate parcels that passed our screening criteria for activity likely to be related to increased attentional control allocated to the cue period of the AX-CPT, we then tested each for a brain-behavior relationship. Specifically, we tested whether cue-related activation in the Proactive condition within each ROI was correlated with the A-cue Bias behavioral metric of proactive control. Eleven of the candidate parcels passed this brain-behavior correlation analysis, highlighted in red within **Figure 3** (see also Supplementary Material, Section S2.3). We then examined these eleven parcels in greater detail.

Each of the eleven parcels were tested for a more selective brain-behavior relationship, via a within-person mediation analysis. The mediation analysis tests whether the behavioral metric of proactive control, i.e., the change in the A-cue Bias in Proactive relative to Baseline, can be linked to the increase in A-cue activation in this condition (Proactive relative to Baseline). Following procedures outlined in Judd et al., (2001), each of the 11 candidate ROIs was subjected to a formal test of within-person mediation. Specifically in addition to the behavioral and neural criteria mentioned above (A-cue Bias Proactive > A-cue Bias Baseline and Reactive; A-cue Bias Proactive correlated with A-cue activation Proactive) we also tested whether: 1) A-cue neural activation correlated with A-cue Bias within Baseline and Proactive respectively; 2) the Proactive – Baseline change in A-cue Bias correlated with the Proactive – Baseline change in A-cue activation; and 3) a significant regression is obtained with the following equation:

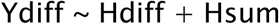

Where Ydiff indicates the difference in A-cue Bias in Proactive vs. Baseline, Hdiff indicates the difference in A-cue activation in Proactive vs. Baseline, and Hsum indicates the overall A-cue activation across both conditions. Within this regression equation, Hdiff tests for mediation; conversely, Hsum tests for the presence of moderation rather than mediation (e.g., whether A-cue activity is differentially related to A-cue Bias in Proactive rather than Baseline).

Two adjacent parcels in the left somatomotor cortex (lMOT; circled parcels in **Figure 3a**) passed all criteria for mediation; moreover, there was no evidence for significant moderation in either parcel (for additional visualization of the difference in A-cue activation and A-cue Bias between Baseline and Proactive, see Supplementary Material, Section S3.1). Because these two parcels are contiguous, for further analyses we aggregated them at the voxel level into a single combined parcel by creating a new mask. As expected, in this lMOT combined parcel A-cue activation correlated significantly with A-cue Bias (ρ=0.272, p=0.003), and then passed subsequent mediation testing (**Table 3**; difference correlation plotted in **Figure 4a**; ρ=0.23, p=0.01; see Supplementary Material, Section S3.3 for model output and visualization of the two parcels separately). To better visualize the relationship between A-cue activation and A-cue Bias, we sorted the participants by A-cue Bias score, forming extreme subgroups (N=30 each) of the participants. The subgroup exhibiting high levels of behavioral A-cue Bias had increased cue-related activation in the Proactive condition within the lMOT, both for A-cues and A > B-cue-related activity, compared to the Baseline and Reactive conditions; this pattern was not observed for individuals exhibiting low-levels of behavioral A-cue Bias (**Figure 4b**).

**Figure 4:**
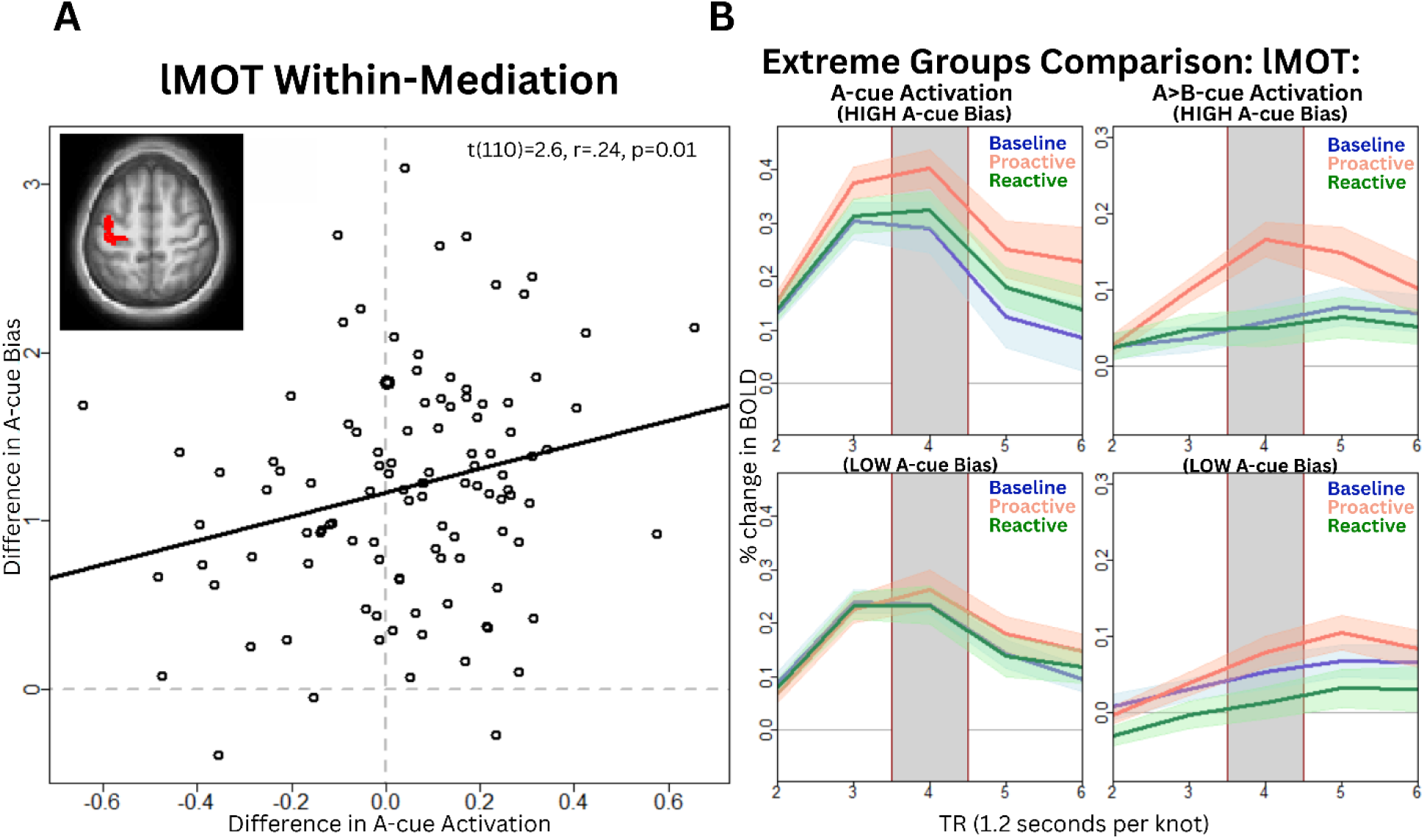
**a)** Plot visualizing the relationship between A-cue activation and A-cue Bias in the parcels whose voxels were combined in the main analysis, LH SomMot 22 and LH SomMot 26. Differences in A-cue knot activity between the Proactive and Baseline conditions can be said to mediate the differences in A-cue behavioral bias if: 1) the behavioral bias in Proactive is significantly greater than in Baseline, and 2) differences in A-cue brain activity is predictive of A-cue Bias (Judd et.al, 2001). **h)** Extreme groups visualization of the relationship between A-cue Bias and cue-related activity. Upper plots show activation in the N=30 participants with highest A-cue Bias scores, while the lower plots show activation in the N=30 participants with lowest A-cue Bias. Left plots show A-cue activation across the Proactive, Baseline, and Reactive conditions in the two groups. Right plots show A > B cue activation across conditions in these groups. Timccourses were created with robust means and standard errors (SEM) trimmed at 0.1.

**Table 3.**
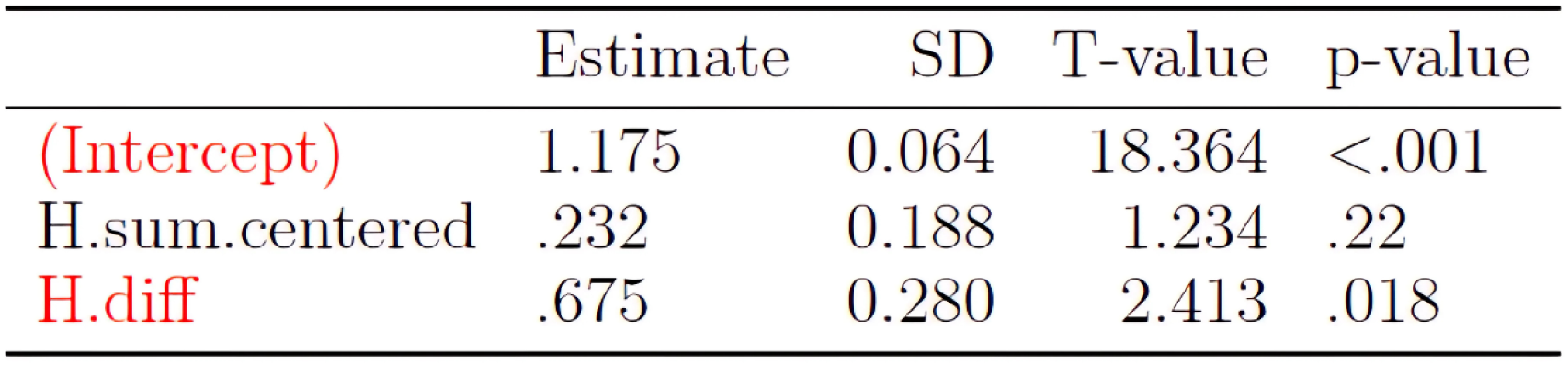
Combined 1MOT combined parcel A-cue activity can be said to mediate (H.diff) or moderate (H.sum.centered) differences in A-cue Bias if the neural activity difference between Proactive and Baseline is predictive of behavioral measure difference (i.e. Y.diff ∼ H.sum + H.diff); Y=behavioral measure i.e. A-cue Bias; Y.diff = Proactive-Baseline behavioral A-cue Bias; H=neural measure i.e. knot 4 A-cue activity; H.diff = Proactive-Baseline neural A-cue activity.

### WMC effects on neural indices of proactive control

Focusing on the same 40 candidate parcels, we next examined whether individual differences in WMC modulated A-cue activation, now treated as a neural index of proactive control. Interestingly, no correlations were observed within the left somatomotor ROIs. Instead, this relationship was found in two other ROIs (see **Figure 3a** parcels highlighted in green: right DLPFC (rDLPFC; ρ=0.22, p=0.015) and left parieto-occipital cortex (ρ=0.22, p=0.016). Based on the behavioral analyses and the findings from Lin et al (2022), we then tested whether this association between WMC and A-cue activation was selective to the Proactive condition. To test for selectivity, we computed the correlation of the Proactive – Baseline A-cue contrast. Only the rDLPFC was significant in this test (ρ=0.21, p=0.027; circled green parcel in **Figure 3a**). As an alternative, and potentially statistically more powerful approach, we utilized partial correlation to remove shared variation with the Baseline condition. Again, only the rDLPFC showed a significant partial correlation (ρ=0.24, p=0.009; visualization in **Figure 5a**; see Supplementary Material, Section S4.1 for partial and zero-order correlations table). To visualize this effect, we again split the data into two extreme subgroups (N=30), this time based on the highest and lowest WMC scores. The high WMC subgroup had increased cue-related activation in the Proactive condition within the rDLPFC, both for A-cues and A > B-cue-related activity, compared to the Baseline and Reactive conditions; this pattern was not observed in low WMC individuals (**Figure 5b**).

**Figure 5:**
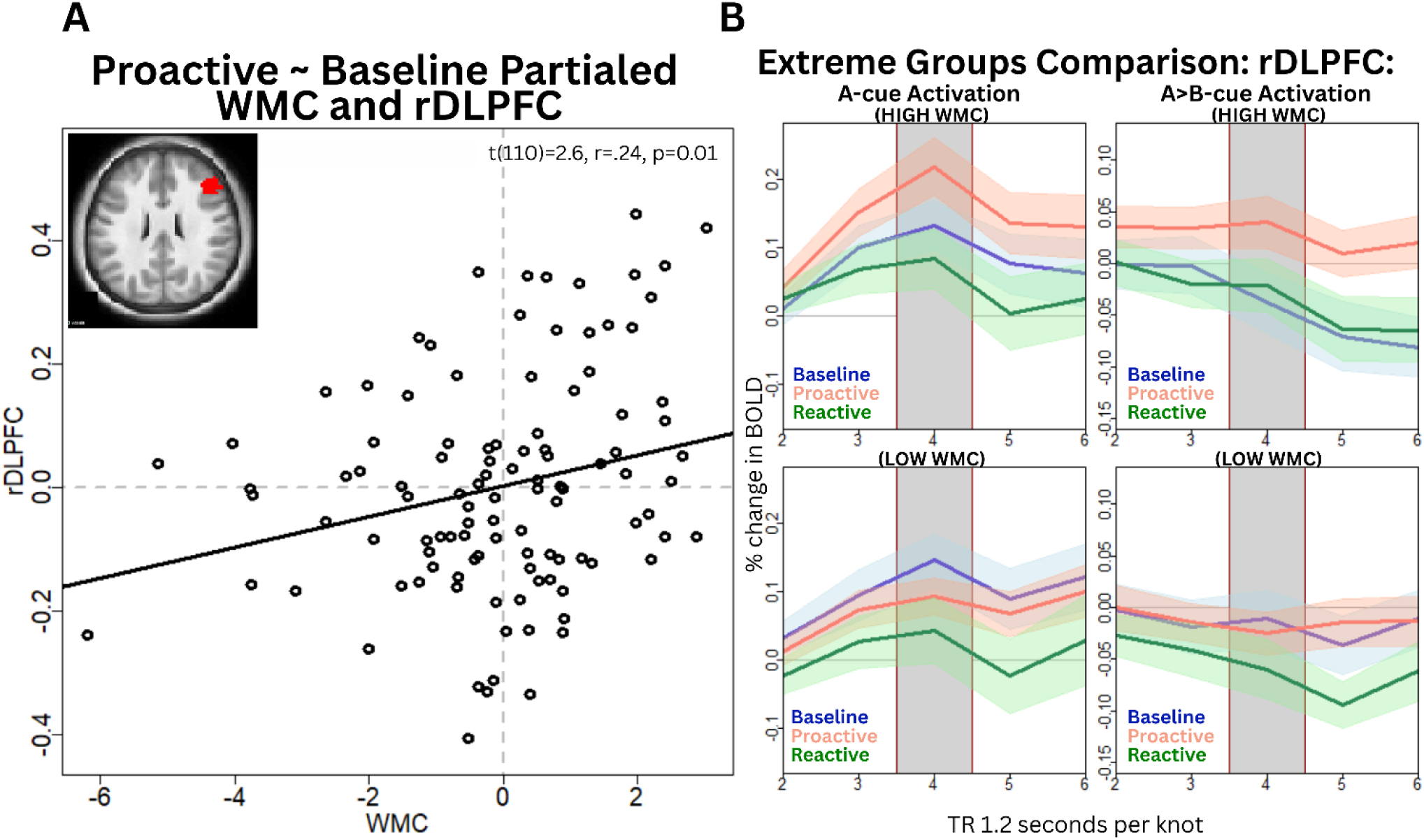
**a)** Plot showing the partial correlation between WMC and Proactive rDLPFC activation. Partialling was used to remove shared variance between the Baseline and Proactive session, **b)** Extreme groups visualization of the individual differences effect in cue-related activity. Upper plots show activity in the N=30 participants with the highest WMC scores, while the lower plots show activity in the N=30 participants with lowest WMC. Left plots show A-cue activation across the Proactive, Baseline, and Reactive conditions in the two groups; Right plots show A > B cue activation across conditions in these groups. Timecourses were created with robust means and standard errors (SEM) trimmed at 0.1.

### A WMC – proactive control neural circuit

Up until this point, the results indicated that A-cue activation in left somatomotor cortex predicted the A-cue Bias behavioral index of proactive control, while WMC predicted A-cue activation in right DLPFC. We next examined whether these two effects were linked, with the two parcels forming a neural circuit. To do this, we first tested whether the A-cue related activation in the two regions was associated across individuals, and then explored the relationships using latent path analysis.

In fact, this was the case, as a highly significant between-subjects correlation was present among the ROIs (rDLPFC-lMOT: r = 0.40, p<0.001). Thus, based on the set of significant zero-order correlations within the Proactive condition (1:WMC – A-cue Bias; 2: WMC – rDLPFC A-cue activation; 3: rDLPFC – LMOT A-cue activation; 4: LMOT A-cue activation – A-cue Bias; see Supplementary Material, Section S4.1 for correlations within the Proactive and Baseline conditions) we conducted a formal latent path analysis. This analysis tests whether in the indirect pathway: (a) individuals with higher WMC have higher rDLPFC activity, (b) higher rDLPFC activity results in increased LMOT activation during A-cue presentation, and (c) this results in an increased likelihood to make a target response; and further, whether this indirect pathway explains significant variance in (d) the direct path of WMC predicting A-cue Bias. Importantly, this latent path analysis was significant, while the rDLPFC and A-cue Bias as well as lMOT and WMC were not significantly related to each other on their own, suggesting indirect path mediation (see Supplementary Material, Section S4.3 for the full path analysis results using zero-order Proactive condition values).

To further probe the selectivity of this effect to the Proactive condition, we re-ran the latent path analysis while controlling for the Baseline condition. The first analysis subtracted out the Baseline condition values from Proactive for each variable. Again, similar results were obtained (indirect a*b*c: β= 0.007, p=0.084, CI95%[0.000, 0.017]; total path d+(a*b*c): β=0.07, p=0.11, 95%CI[-0.006, 0.161]; see Supplementary Material, Section S4.4 for full results). The second analysis more rigorously controlled for shared variance with Baseline through a partial correlation approach with each variable (see Supplementary Material, Section S4.1 for a comparison of zero-order and partial correlations). With this partial correlation approach, all of the significant zero-order relationships were still observed ((a) WMC-rDLPFC ρ=0.24, p=0.009; (b) rDLPFC-LMOT ρ=0.36, p=0.009; (c) lMOT-Proactive bias ρ=0.27 p=0.004), and more importantly the full indirect path remained significant (indirect a*b*c: β=0.010, p=0.015, CI 95%[0.003,0.019]; total path total d+(a*b*c): β=0.081, p=0.038, CI 95%[0.010,0.163]). In contrast, the direct path was no longer significant (WMC-Proactive bias ρ=0.16, p=0.80), suggesting further support for mediation of the behavioral relationships by rDLPFC-lMOT neural circuit (see **Table 4**; **Figure 6**). The overall model was not an excellent fit (SRMR=0.069, RMSEA =0.182, CFI=0.809), partially due to small correlations not allowing a full causal pathway to emerge using the current univariate and composite WMC scores (see **Table 4** for full model results, **Figure 6** for a visualization, and Supplementary Material, Section S4.5).

**Figure 6:**
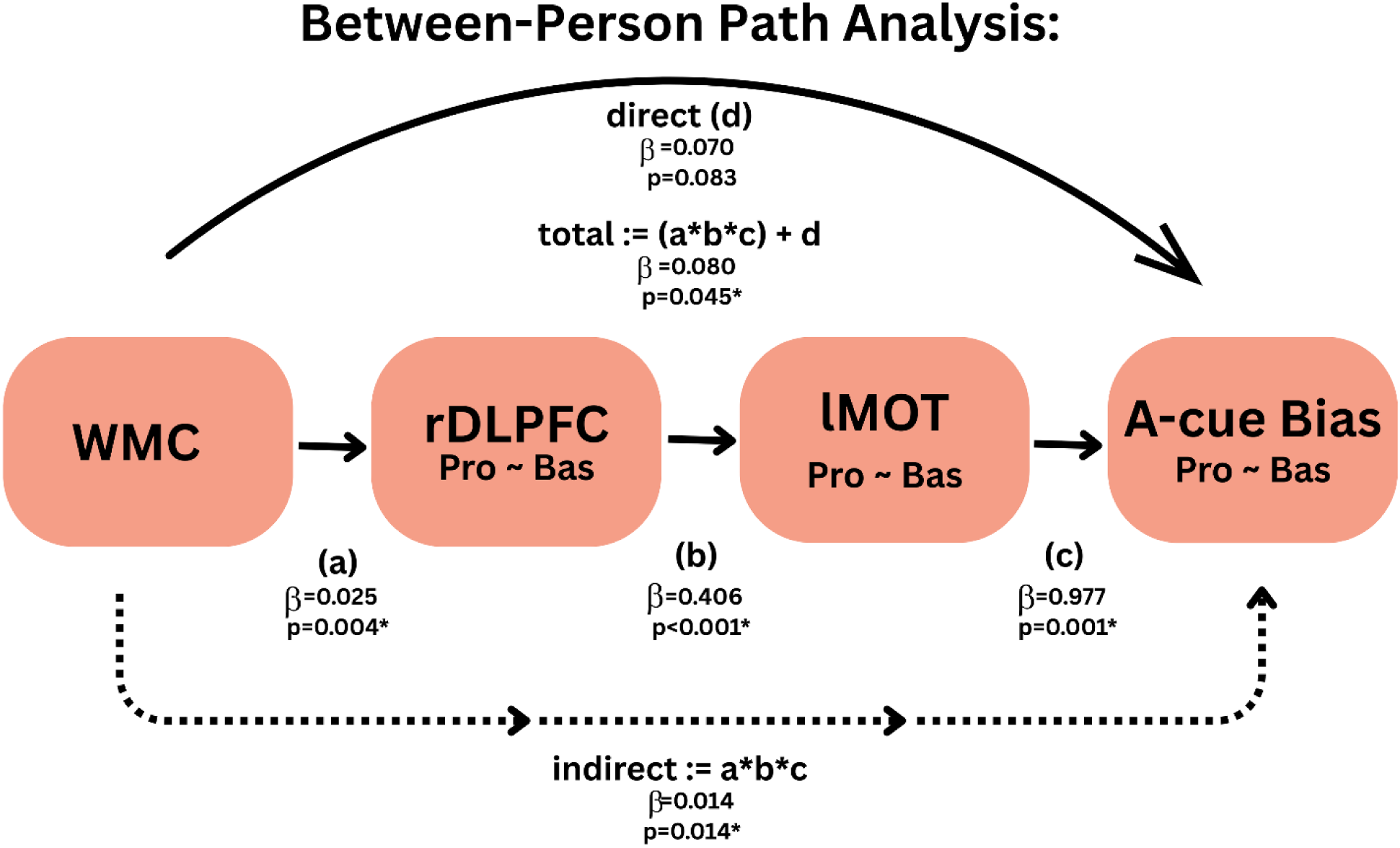
Visualization of between path analysis using residualized proactive session values with significant beta estimates listed. In this model, the direct path was no longer significant, but the indirect (a*b*c) and total d + (a*b*c) was significant. For full model results see **Table 4**

**Table 4.**
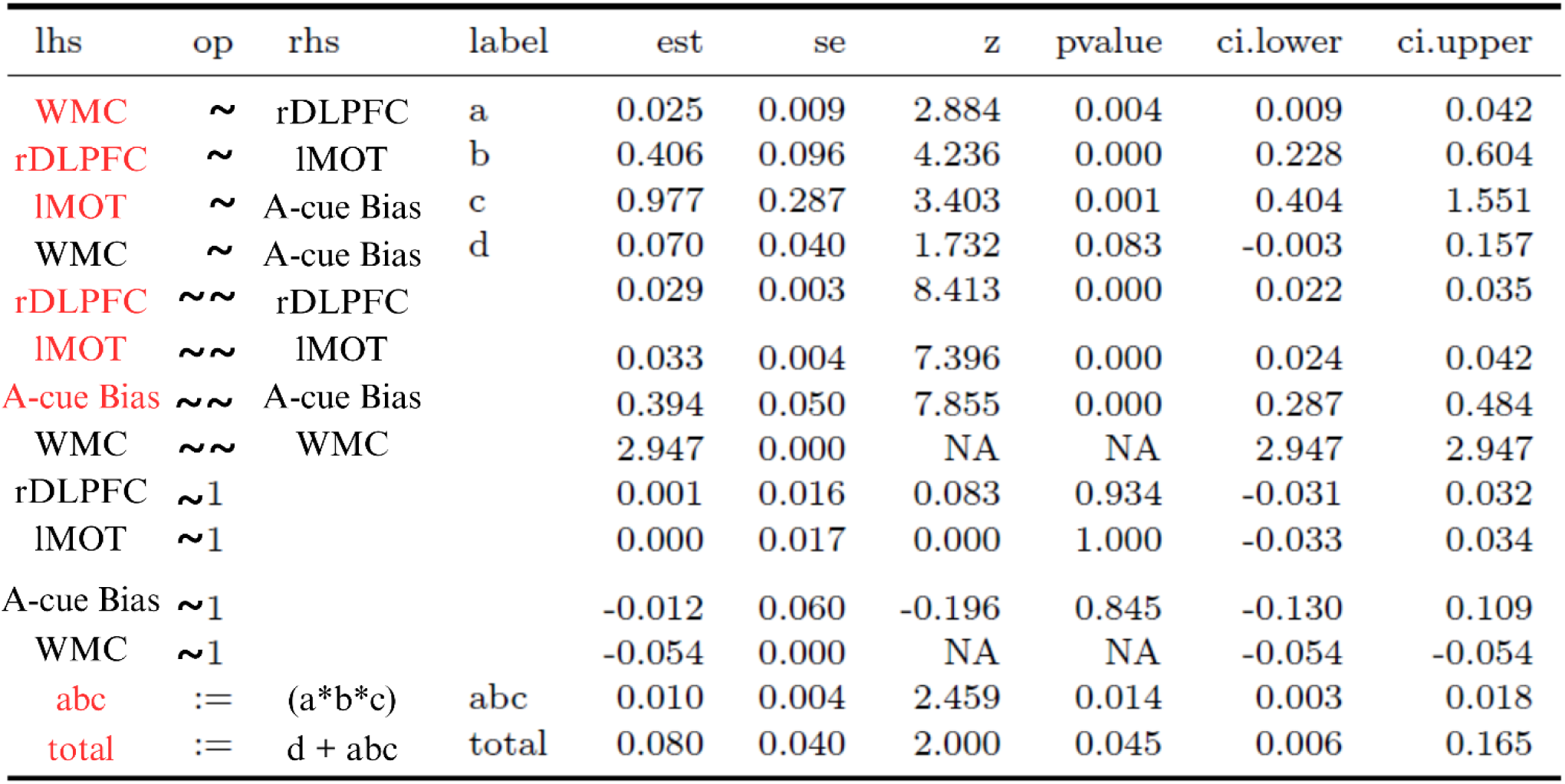
Indirect results using Proactive session values after partialling out shared variance from the baseline session. This model tests for an additive relationship between WMC-rDLPFC-lMOT-A-cue Bias. Rows with RED text denote a confidence interval that is all positive or negative; LHS: left hand side; RHS: right hand side; OP:= operator; ∼ regression; — (residual)(co)variance; 1 intercept; NOTE WMC is a static measure mediation path testing for (a) individuals with higher WMC have higher rDLPFC activity, (b) higher rDLPFC activity results in increased 1MOT activation during A-cue presentation, and (c) which results in an increased likelihood to make a target response, and further, whether this pathway explains significant variance in (d) the direct path of WMC predicting A-cue Bias.

| lhs | op | rhs | label | est | se | z | pvalue | ci.lower | ci.upper |
| --- | --- | --- | --- | --- | --- | --- | --- | --- | --- |
| WMC | ~ | rDLPFC | a | 0.025 | 0.009 | 2.884 | 0.004 | 0.009 | 0.042 |
| rDLPFC | ~ | IMOT | b | 0.406 | 0.096 | 4.236 | 0.000 | 0.228 | 0.604 |
| IMOT | ~ | A-cue Bias | c | 0.977 | 0.287 | 3.403 | 0.001 | 0.404 | 1.551 |
| WMC | ~ | A-cue Bias | d | 0.070 | 0.040 | 1.732 | 0.083 | -0.003 | 0.157 |
| rDLPFC | ~~ | rDLPFC |  | 0.029 | 0.003 | 8.413 | 0.000 | 0.022 | 0.035 |
| IMOT | ~~ | IMOT |  | 0.033 | 0.004 | 7.396 | 0.000 | 0.024 | 0.042 |
| A-cue Bias | ~~ | A-cue Bias |  | 0.394 | 0.050 | 7.855 | 0.000 | 0.287 | 0.484 |
| WMC | ~~ | WMC |  | 2.947 | 0.000 | NA | NA | 2.947 | 2.947 |
| rDLPFC | ~1 |  |  | 0.001 | 0.016 | 0.083 | 0.934 | -0.031 | 0.032 |
| IMOT | ~1 |  |  | 0.000 | 0.017 | 0.000 | 1.000 | -0.033 | 0.034 |
| A-cue Bias | ~1 |  |  | -0.012 | 0.060 | -0.196 | 0.845 | -0.130 | 0.109 |
| WMC | ~1 |  |  | -0.054 | 0.000 | NA | NA | -0.054 | -0.054 |
| abc | := | (a*b*c) | abc | 0.010 | 0.004 | 2.459 | 0.014 | 0.003 | 0.018 |
| total | := | d + abc | total | 0.080 | 0.040 | 2.000 | 0.045 | 0.006 | 0.165 |

Finally, we conducted a series of control analyses to test the robustness of the observed path analysis results. First, we conducted latent path analysis using data from the Baseline condition (instead of Proactive). This resulted in a non-significant indirect and direct path (abc: β=-0.001, n.s.; Total: β=0.033, n.s.; see Supplementary Material, Section S4.6). Note that we did not try a parallel analysis with the Reactive condition, given that the Reactive conditio was not meant provide a direct contrast with Proactive, as it engages a different control mode of operation. Next, we re-ran the path analysis using the Proactive condition data after partialling out shared variance from Baseline, but with the dependencies in a different order (i.e. WMC-LMOT-rDLPFC-A-cue Bias instead of WMC-rDLPFC-lMOT-A-cue Bias) to more formally test whether the proposed causal directionality operated in a top-down manner via the PFC preparing for a motor response (i.e., directionally ordered), rather as opposed to a simple correlation. The indirect path was again non-significant (abc: β=0.000, n.s.; total: β=0.101, p=0.014, CI 95%[0.026, 0.187]; see Supplementary Material, Section S4.7 for full results). Together, these results suggest that the observed effects are quite reliable and selective to both the Proactive condition, and to the rDLPFC – LMOT circuit.

## Discussion

The current study is the first of its type, leveraging the behaviorally validated DMCC variants of the AX-CPT, to test for individual differences in the neural mechanisms of cognitive control with a sample size that is sufficiently well-powered (N>100) for an experimental-correlational design. The study was aimed at addressing two key questions: 1) does higher working memory capacity (WMC), as measured by two span tasks, uniquely enhance behavioral metrics of proactive control, replicating prior results? 2) what are the neural mechanisms that mediate this relationship? We discuss how the findings provide clear answers to each of these questions, in turn.

### WMC effects on behavioral metrics of proactive control

With regard to the behavioral findings, our results provide rigorous quantitative support for a preferential relationship between WMC and proactive control within the AX-CPT paradigm, by successfully replicating prior findings (Lin et al., 2022). Importantly, this replication occurred in a new sample with different demographic characteristics (i.e., narrower age range and with additional screening to meet neuroimaging inclusion criteria), and who performed the AX-CPT in a qualitatively distinct testing environment (i.e. neuroimaging vs. online). The use of Bayesian sequential updating approaches enabled strong statements regarding the consistency of WMC in exerting selective effects on proactive control. Moreover, this pattern of selectivity suggests that WMC influences the *ability* to implement, rather than a general *tendency* to use proactive control. Specifically, individuals with higher WMC seem to be better at actively maintaining and preparing to respond based on contextual cues, when having an explicit goal or instruction to do so, but do not inherently default to proactive processing.

Given contradictory findings with prior studies that have also utilized experimental-correlational designs with the AX-CPT, our findings are important in providing additional evidence and clarification regarding WMC’s preferential relationship with proactive control. Specifically, Rosales et al. (2022) found WMC was not selectively related to proactive control, but instead exerted a more general influence on performance via faster RTs across trial types. Given our complementary findings with Lin et al. (2022), we believe methodological factors in the different AX-CPT variants explain the mixed findings. Critically, the DMCC variants of the AX-CPT were designed to remove potential confounds of trial frequency and type across control conditions, and to better differentiate the selective enhancement of reactive control vs. the reduced engagement of proactive control, effects that had been confounded in prior designs (Rosales et al., 2022). In particular, we used item-specific cueing to induce reactive control; no-go trials were present and matched across all conditions. Matching trial-type frequencies across conditions served the dual purpose of both removing a potential confound and reducing the tendency to rely on proactive control under baseline conditions (see Braver et al., 2021 for more details). Furthermore, the aggregated trial-level Bayesian mixed modeling approach used in the current study was designed to assess condition-level interactions, thus allowing for more sensitivity. This type of analysis avoids issues associated with statistical non-independence of repeated subject-level measures, by using the entirety of the dataset and without relying on summary or subtraction level scores or multiple analyses. Our hope is that other researchers will be able to make use of these current results, taking advantage of the cumulative approach to psychological and cognitive neuroscience findings afforded by Bayesian modeling and sequential updating procedures (Brand et al., 2017; Ly et al., 2019). In particular, the posterior estimates reported here can be utilized by other researchers as priors in future studies examining proactive control in the AX-CPT and the moderating role of WMC, similar to how we did with Lin et al., (2022).

### WMC effects on neural indices of proactive control

A number of prior studies have used fMRI and EEG methods with the AX-CPT (or a closely-related variant known as Dot Pattern Expectancy task; DPX) to investigate the neural mechanisms of proactive control in terms of goal maintenance and context processing. Yet this prior work has primarily focused on group average activation patterns, rather than individual differences (Blackman et al., 2016; Braver et al., 2009; Gómez-Ariza et al., 2017; Janowich & Cavanagh, 2018; Lesh et al., 2013; Mäki-Marttunen et al., 2019a; Pulopulos et al., 2022; Ryman et al., 2019; Sharifian et al., 2021; Wiwatowska et al., 2022). Like the current study, prior studies have focused on DLPFC in WM and cognitive control, treating it as a hub for the representation of both stimulus content and top-down signals directing attention and action (Barch et al., 1997; Constantinidis & Klingberg, 2016; Holmes et al., 2005; Paxton et al., 2008; Perlstein et al., 2001; Vogel et al., 2005). More broadly, the DLPFC has been strongly linked to goal, context, or task-set maintenance aspects of working memory—processes that are dissociable from general task difficulty and conflict-related incongruency (Barch et., al 1997; Barch et al., 2001; Holmes et al., 2005; Lesh et al., 2013; MacDonald & Carter 2003; MacDonald, Carter, et al., MacDonald et al., 2000).

Several studies have found temporal shifts in activation of the DLPFC from the cue (i.e. proactive control) to the probe (i.e. reactive control) (or vice versa) as a function of instruction, training, or item-specific cueing (Braver et al., 2009; Gonthier et al., 2019; Gonthier, Macnamara, et al., 2016; Li et al., 2018). A central goal of the current study was to utilize both brain-behavior relationships and individual difference approaches to advance understanding of proactive control mechanisms. To accomplish this, in addition to using trial-level estimates for behavioral metrics, our current study focused exclusively on cue-period activation, to identify different patterns of cue-based preparation associated with experimental manipulations and whether these relate to individual differences in WMC.

Consistent with the DMC theory and prior work (M. Boudewyn et al., 2019; Braver, 2012; Braver et al., 2009; Lesh et al., 2013; Lopez-Garcia et al., 2016; Paxton et al., 2008; Pulopulos et al., 2022) we found that the Proactive condition was associated with increased activation in the DLPFC during A-cues, as well as differential activation in A- vs. B-cues, compared to both Reactive and Baseline conditions. The fMRI BOLD signals in DLPFC are thought to reflect a proxy measure of neural population activity that might be influenced by sustained neuromodulatory activity (such as from the midbrain dopamine system), with increased signal values indicating higher capacity or more stable representations (Constantinidis & Klingberg, 2016; D’Ardenne et al., 2008; Durstewitz et al., 2000; Edin et al., 2009). Consequently, we interpret this increased DLPFC activity as reflecting an increase in processing or representational capacity used for WM and attention control. That is, more attentional control was being allocated to A-cues over B-cues under conditions in which participants were more strongly oriented towards A-cue response preparation from the proactive strategy training manipulation. Strikingly, we found that right DLPFC was the only area with both significant A>B cue activation and a relationship with WMC. This pattern suggests that WM-based preparation in the Proactive condition was qualitatively distinct, but also that the degree of preparation was related to WMC, with high WMC individuals showing larger effects. Indeed, in visualizing the cue-related activation patterns for the individuals with highest and lowest WMC this point becomes clear, since individuals with high WMC showed the greatest differentiation of the Proactive condition, in terms of both A-cue and A>B-cue activation patterns (**Figure 5b**).

While individuals with greater WMC exhibited stronger right DLPFC activation and A-cue Bias (a behavioral index of proactive control), the A-cue Bias effect was somewhat surprisingly more tightly associated with activation in contralateral motor cortex, rather than PFC *per se*. However, this finding make sense, considering that heightened expectations for an upcoming target probe would involve priming of a right-handed button press, based presumably on preparatory response activation in the left motor cortex (lMOT). Since PFC is involved in gating, maintaining, and relaying motor information, potentially via modulation from midbrain dopamine inputs (Constantinidis & Klingberg, 2016; Dardenne et al., 2013; Ott & Nieder, 2019), we first tested if the increased lMOT activation mediated the increased A-cue Bias observed within-subjects as a function of contralateral motor-preparation, theoretically through top-down influence from the right DLPFC. The specificity of the WMC-rDLPFC relationship and rDLPFC-lMOT co-activation to the Proactive condition mirrors and extends the idea that WMC preferentially enhances the proactive bias towards target-related response preparation. Indeed, through latent path analyses, we confirmed the presence of a rDLPFC-lMOT neural circuit that translates task goals into active response preparation, with the strength of this circuit influenced by WMC. This suggests individuals with higher WMC were better able to maintain and utilize context cues that are ultimately relayed to behavioral responses (motor action) as a mechanism of proactive control.

### Limitations and Future Directions

Although many studies have used the AX-CPT (and DPX) to study both WMC and neural mechanisms of cognitive control, interpretations of prior findings can be complicated by the fact that the basic task paradigm has variations in instruction, stimuli, trial type frequency, timings, and manipulations. These might have a subtle, and previously under-appreciated impact on the nature of cognitive control strategies engaged, and the importance of WM function. The variants used in the current study were intentionally designed to be qualitatively distinct from those developed within the original AX-CPT, and were implemented with the goal of more cleanly double dissociating proactive and reactive control (Tang et al., 2023). To accomplish this, cue-types were made to be equal in frequency to remove biased conflict-driven confounds and no-go trials were added to reduce the overall reliance on proactive control under baseline conditions, which in turn increased the dynamic range of cognitive control effects across conditions. Finally, unlike prior versions of the AX-CPT, three different experimental variants were created which had the same exact trial structure, but relied on other manipulations, that of instructions and incidental stimulus features, to manipulate utilization of proactive and reactive control.

The approach used here for AX-CPT paradigm design allows for the isolation of proactive control, which can be done by comparing the same A-cue trials across conditions. Moreover, since the A- and B-cues occur with equal frequency within and between conditions, any cue-related effects cannot be due to frequency-related factors (e.g., surprise, novelty) that may be present in standard AX-CPT designs (for which the A-cue is much more frequent than B-cue). Nevertheless, despite the strengths of this approach, and the fact that we observed consistency with other studies that have used the same design (Lin et al, 2022), our results may differ from the prior literature simply due to subtle differences in design between them. As such, it is important to consider these constraints on generality with regard to our findings when placing them in the context of the broader literature, or when designing new AX-CPT studies.

From a methodological standpoint, the current study examined the neural mechanisms of cognitive control from an individual differences perspective, by utilizing conventional univariate statistical analyses, and ROI-based methods to specify neural activation patterns in cognitive control-related brain regions. Univariate statistical approaches have distinct advantages for these kinds of scientific questions, in that they are conceptually simple, with pre-defined ROIs allowing for ease of comparison across individuals and between studies. Nevertheless, other approaches, that rely on trial-by-trial activation patterns and functional connectivity methods (e.g., PPI; Cisler et al., 2014; O’Reilly et al., 2012; Strosche et al., 2021; Yu et al., 2015), or multivariate pattern analysis (MVPA), could yield distinct and complementary findings. It is important to acknowledge that fMRI signals are inherently noisy, and as such, identifying trial-by-trial and/or MVPA-based activation patterns may be challenging due to low signal-to-noise ratio, risks of overfitting, and potential loss of important information when working with high-dimensional fMRI data, particularly when it comes to mapping decoding performance back to neural mechanisms (Etzel et al., 2009; Hebart & Baker, 2018; Kriegeskorte & Bandettini, 2007). Yet now that our results have been established at a univariate level, there may be reduced concerns regarding the statistical challenges of using multivariate fMRI approaches, particularly now that a candidate neural circuit has been identified, which reduces the anatomical search space. Moreover, recent work has suggested that, perhaps counter-intuitively, enhanced reliability in fMRI studies might be obtained from single-trial and MVPA-based estimates of neural activation patterns (Freund et al., 2025b).

Thus, adoption of single-trial and MVPA approaches may represent a highly fruitful next step for research on these questions. Such approaches might be particularly useful for examining the somewhat surprising lack of twin effects observed in the current results. Although the DMCC dataset included a subset of monozygotic twins, these twin pairs were not found to show reliable similarity in the nature of the WMC – proactive control relationships either at the behavioral or neural level aside from their composite WMC score (see Supplementary Material, Section S5 for results regarding twins). In other work, we have shown that twin effects can be observed within the DMCC dataset (Tang et al., 2021). Yet these findings were observed primarily in terms of neural pattern similarity; that is, through multivariate rather than univariate approaches. Indeed, we also found that within a larger neuroimaging dataset associated with the Human Connectome Project, the degree of neural pattern similarity varied substantially across different twin pairs (Etzel et al., 2020). Thus, it may be the case that multivariate, rather than univariate, approaches are the most sensitive and appropriate for detecting and decomposing twin-related similarity and individual differences related to cognitive control.

Another important limitation of the current work, and with standard neuroimaging approaches more generally, is that the inferences are primarily correlational rather than causal in nature. As such, an important direction to extend this current work would be to combine it with neurostimulation approaches, as these allow for stronger inferences about the causal nature and functional mechanisms of top-down control (D’Ardenne et al., 2012). One study found transcranial direct current stimulation (tDCS) over the DLPFC enhanced WM training practice effects for a verbal-memory task and had near transfer effects for novel WM tasks (Richmond et al., 2014). However, another study utilizing the AX-CPT found tDCS over the right DLPFC disrupted proactive tendencies on conflict trials (AY and BX), and increased reliance on reactive control, with no modulatory effects present in the left DLPFC (M. A. Boudewyn et al., 2015; Gómez-Ariza et al., 2017; Pulopulos et al., 2022). It is possible stimulation over the right DLPFC, which has been associated with temporal shifts between proactive and reactive control (Braver et al., 2009), is interrupting the dopamine-gating system by reducing the ability to actively maintain cue-related contextual information, thus pushing participants to rely more heavily on reactive control. Conversely, it is also possible that left DLPFC can contribute to WMC influences on proactive control, but is not necessary when right DLPFC signals can also be utilized. Neurostimulation methods have the promising potential to help determine causality, dissociate areas involved in different processes, and provide potential interventions for populations with WM deficits, or low WMC. Our findings suggest that these might be most effective when targeting the right DLPFC and/or left motor cortex, particularly during the cue-probe delay period, but further work is needed to provide stronger support for this claim.

An additional avenue to explore would be WMC training interventions, to see how they affect performance and activation in the regions of interest, or to potentially combine training with neurostimulation. Indeed, human neuroimaging studies suggest that WM training changes activation patterns most consistently in fronto-parietal regions relative to pre-training findings (Constantinidis & Klingberg, 2016; Olesen et al., 2004; Takeuchi et al., 2010; Thompson et al., 2016). Our work may provide more anatomical specificity in terms of a starting point from which to look for such training effects Further support of WM training’s impact on activation comes from primate electrophysiology studies in which PFC neurons increase during the delay period in a WM task post-training, aligning with behavioral improvements (Meyer et al., 2011). If after WM training, the rDLPFC-lMOT neural circuit is enhanced, along with behavioral indexes of proactive bias, this would provide further support for this circuit in the context of goal maintenance and response preparation. Further, a comparison of WMC training, neurostimulation, and a combination of the two interventions, which examine the longevity of, and potential transfer effects in this circuit, would be quite useful in providing insight on the relative effectiveness of these two types of interventions separately and in combination with each other.

Finally, a key feature of the DMCC project, and associated task battery is that it provides a heterogenous set of manipulations and cognitive paradigms, but with a common goal of assaying proactive and reactive control in a parallel fashion. Consequently, the DMCC battery provides a natural means of investigating fronto-parietal cortical neural circuitry related to WMC and proactive control in relation to the findings observed here, but from within different task paradigms that tap into different facets of cognitive control (i.e. verbal working memory [Sternberg], selective attention [Stroop], and task-switching). The DMCC task battery, analysis code, and much of the neuroimaging data are open-source and publicly available. Furthermore, the use of Bayesian modeling this allows for future groups to make use of, or add to the data and findings described here, much as we did in the current study, that built on the findings of Lin et al., (2022). It is our hope that the current findings and DMCC project approach will be useful in stimulating the scientific community to help us discover how individual differences contribute to key dimensions of variation in the mechanisms of cognitive control.

## Acknowledgements

We thank Yanli Lin for developing the hierarchical Bayesian modeling approach to the analysis of DMCC behavioral AX-CPT datasets, and for technical assistance in applying this approach to the current fMRI dataset. We thank the members of the Cognitive Control and Psychopathology (CCP) lab for fruitful discussions regarding this project. Early portions of this work were presented at the Cognitive Neuroscience Society meeting in 2023. Funding support for the project was provided NIH R37 MH066078 and ONR MURI N00014-22-S-FO.

## Open Practices Statement

The data and materials for all analyses are available at https://osf.io/d4k8n except for twin-related information. Twin-related information can be requested via email after submitting a formal request and agreement to HCP https://www.humanconnectome.org/study/hcp-young-adult/document/restricted-data-usage. None of these analyses were pre-registered.

