## Supplementary Materials for "Neural Mechanisms Supporting the Relationship between Working Memory Capacity and Proactive Control"

### Supplemental Material

source and input files available at `https://osf.io/d4k8n`
compiled May 9, 2025
Results Supplement 1 for "Neural Mechanisms Supporting Proactive Control" by Rebecca Feldman, Maya Quayle, Joset Etzel, and Todd S. Braver.

This is a `knitr` file (`https://yihui.name/knitr/`); see the `.rnw` file with the same name as this `.pdf` for the R code to generate all figures and results. To compile, change the `in.path` variable to the location of the `osfDATA/` directory downloaded from `https://osf.io/d4k8n`.

### Supplemental 1: Behavioral Analyses

#### S1.1 Establishing a Working Memory Capacity (WMC) metric

First, following the procedure from Lin et. al. (2022) we created a WMC composite score using z-scored, and subsequently summed SYMSPAN and OSPAN results. Similar to prior results, raw scores are highly correlated ($\rho$=0.547, p <0.001)

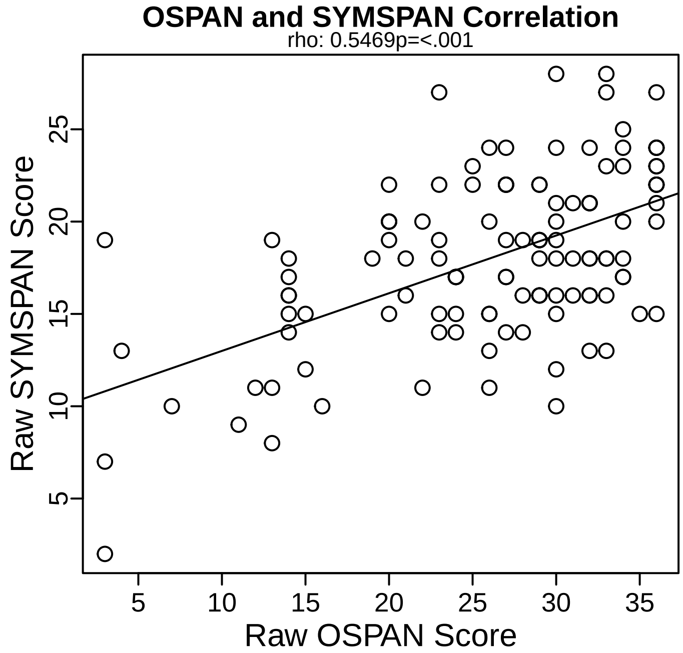


Table 1: OSPAN, SYMSPAN, and WMC summary statistics. WMC composite scores were calculated by z-score normalizing relative to the group mean and standard deviation) and summing the SYMSPAN and OSPAN scores.

| | M | SD | Skew | Kurtosis | Correlation |
|---|---|---|---|---|---|
| OSPAN | 26.093 | 8.123 | -0.999 | 0.440 | 0.547*** |
| SYMSPAN | 18.025 | 4.642 | -0.344 | 0.385 | - |
| WMC COMPOSITE | -0.042 | 1.719 | -0.839 | 0.935 | - |

[a] Note: This is Table 1 in the main text

#### S1.2 Establishing trial-level A-cue Bias metric

Next, we used the hierarchical mixed-level Bayesian model developed in Lin et. al, (2022) for a trial-level metric of A-cue Bias as opposed to the accuracy based summary score measure typically used. The summary score is calculated as the log-linear correction of hits (AX) and false alarms (AY) calculated as: (qnorm(AX.h) + qnorm(AY.fa))/2. The

trial-level metric is computed as the log-liklihood of making a target response following an A-cue regardless of if they are right (hit) or wrong (false alarm). In this case, each persons random effect beta value is stored as their A-cue Bias and was calculated for each session independently and then compared to the summary score to make sure the metrics are comparable.
One of the reasons we chose to use the trial-level metric as it parallels the behavioral replication of WMC's preferential interaction with the proactive session. Additionally, our neural metric that only takes into account the cue-period, and not reliant on any accuracy or probe-based activation. Finally, this method sidesteps analytic limitations of summary scores by avoiding aggregating across trials and takes trial-level variability into account (Rouder and Haaf, 2019; Lin et al., 2022).

##### S1.2a Descriptive Statistics:

**Summary Level Derived Measures, Trial Type RT, and accuracy from the AX-CPT across sessions**

| | Trial Type | Cue RT Mean (SEM) | Probe RT Mean (SEM) | Error Mean (SEM) |
|---|---|---|---|---|
| **Baseline** | | | | |
| | AX | 445.419 (2.281) | 434.2 (2.121) | .07 (.025) |
| | AY | 442.07 (4.461) | 588.364 (4.893) | .066 (.04) |
| | BX | 476.489 (4.703) | 609.684 (7.644) | .189 (.084) |
| | BY | 482.638 (2.52) | 477.413 (2.351) | .043 (.019) |
| | A-nogo | 435.657 (4.032) | - | .102 (.059) |
| | B-nogo | 480.482 (5.113) | - | .151 (.081) |
| **Proactive** | | | | |
| | AX | 361.055 (2.075) | 334.064 (1.933) | .076 (.03) |
| | AY | 355.977 (4.099) | 567.234 (5.588) | .321 (.105) |
| | BX | 390.967 (4.904) | 431.161 (6.633) | .131 (.069) |
| | BY | 387.341 (2.4) | 372.698 (2.113) | .046 (.019) |
| | A-nogo | 360.487 (4.36) | - | .23 (.095) |
| | B-nogo | 375.935 (4.581) | - | .334 (.112) |
| **Reactive** | | | | |
| | AX | 389.397 (2.277) | 395.192 (2.411) | .097 (.034) |
| | AY | 388.989 (4.545) | 592.135 (5.629) | .095 (.056) |
| | BX | 428.743 (5.195) | 584.961 (6.726) | .136 (.066) |
| | BY | 422.833 (2.513) | 430.781 (2.39) | .05 (.021) |
| | A-nogo | 391.499 (4.688) | - | .093 (.051) |
| | B-nogo | 420.698 (5.13) | - | .13 (.071) |

##### S1.2b Trial-Level A-cue Bias:

A-cue bias was calculated using the following equation within each session (run individually): **ProbeSlide.Response ∼ Trial.Type + (1 + Trial.Type | subject)**
Subject-level random effects are stored as each's participant's A-cue bias for later brain-behavior analyses.
**Proactive Trial-Level A-cue Bias output**

```
head(pro.data)
# create and run brms NO priors use uniformed
pro.model <- ProbeSlide.RESP ~ 0 + Intercept + TrialType + (1 + TrialType | sub.id)
# read in the model -- saves time compiling
pro.brms <- readRDS(paste0(in.path, "S1/proTrialBias.rds"))
# if the model hasn't been run, uncomment and run, save, and update path to read in
# pro.brms <- brm(pro.model,
#                       data = pro.data,
#                       family= bernoulli(link = "logit"),
#                       cores = 6,
#                       sample_prior = TRUE,
```

| Derived Measure | Error Mean (SD) |
|---|---|
| **Baseline** | |
| A-cue Bias | .121 (.286) |
| BX Interference | .925 (.627) |
| d'-context | 2.604 (1.124) |
| Proactive Behavioral Index | -.29 (.419) |
| **Proactive** | |
| A-cue Bias | .523 (.388) |
| BX Interference | .688 (.494) |
| d'-context | 2.722 (.995) |
| Proactive Behavioral Index | .312 (.465) |
| **Reactive** | |
| A-cue Bias | .082 (.311) |
| BX Interference | .652 (.554) |
| d'-context | 2.601 (1.105) |
| Proactive Behavioral Index | -.084 (.386) |

```
#                        seed = 35)
# saveRDS(pro.brms, paste0(in.path, "S1/proTrialBias.rds"))
s <- summary(pro.brms)
s
# store for later
pro.coef1 <- as.data.frame(coef(pro.brms))
pro.coef1 <- as.data.frame(cbind(rownames(pro.coef1),pro.coef1[,"sub.id.Estimate.Intercept"]))
colnames(pro.coef1) <- c("sub.id", "acue.coef")

##      sub.id TrialType ProbeSlide.RESP ProbeSlide.CRESP ProbeSlide.RTTime            session
## 121 f2499cq        AX          target                2            257801 proactiveBlockList
## 122 f2499cq        AY       nontarget                1            266233 proactiveBlockList
## 123 f2499cq        AX          target                2            280532 proactiveBlockList
## 124 f2499cq        AX          target                2            287781 proactiveBlockList
## 125 f2499cq        AX       nontarget                2            312934 proactiveBlockList
## 126 f2499cq        AX          target                2            358572 proactiveBlockList
##        wk.mem
## 121 -2.355979
## 122 -2.355979
## 123 -2.355979
## 124 -2.355979
## 125 -2.355979
## 126 -2.355979
##  Family: bernoulli
##   Links: mu = logit
## Formula: ProbeSlide.RESP ~ 0 + Intercept + TrialType + (1 + TrialType | sub.id)
##    Data: pro.data (Number of observations: 7056)
##   Draws: 4 chains, each with iter = 2000; warmup = 1000; thin = 1;
##          total post-warmup draws = 4000
##
## Multilevel Hyperparameters:
## ~sub.id (Number of levels: 119)
##                             Estimate Est.Error l-95% CI u-95% CI Rhat Bulk_ESS Tail_ESS
## sd(Intercept)                   0.85      0.09     0.69     1.05 1.00     1512     2439
## sd(TrialType1)                  0.84      0.09     0.67     1.03 1.00     1784     2809
## cor(Intercept,TrialType1)       0.43      0.13     0.15     0.67 1.00      875     1665
##
## Regression Coefficients:
##              Estimate Est.Error l-95% CI u-95% CI Rhat Bulk_ESS Tail_ESS
## Intercept        1.16      0.10     0.96     1.36 1.00     1243     2246
```

```
## TrialType1    -2.38      0.10    -2.58    -2.18 1.00     1855     2424
##
## Draws were sampled using sampling(NUTS). For each parameter, Bulk_ESS
## and Tail_ESS are effective sample size measures, and Rhat is the potential
## scale reduction factor on split chains (at convergence, Rhat = 1).
```

##### Baseline Trial-Level A-cue Bias output

```
bas.model <- ProbeSlide.RESP ~ 0 + Intercept + TrialType + (1 + TrialType | sub.id)
# read in the model -- saves time compiling
bas.brms <- readRDS(paste0(in.path,"S1/basTrialBias.rds"))
# if the model hasn't been run, uncomment and run, save, and update path to read in
# bas.brms <- brm(bas.model,
#                  data = bas.data,
#                  family= bernoulli(link = "logit"),
#                  cores = 6,
#                  sample_prior = TRUE,
#                  seed = 35)
# saveRDS(bas.brms, paste0(in.path,"S1/basTrialBias.rds"))
s <- summary(bas.brms)
s
# store for later
bas.coef1 <- as.data.frame(coef(bas.brms))
bas.coef1 <- as.data.frame(cbind(rownames(bas.coef1),bas.coef1[,"sub.id.Estimate.Intercept"]))
colnames(bas.coef1) <- c("sub.id", "acue.coef")

##  Family: bernoulli
##   Links: mu = logit
## Formula: ProbeSlide.RESP ~ 0 + Intercept + TrialType + (1 + TrialType | sub.id)
##    Data: bas.data (Number of observations: 7139)
##   Draws: 4 chains, each with iter = 2000; warmup = 1000; thin = 1;
##          total post-warmup draws = 4000
##
## Multilevel Hyperparameters:
## ~sub.id (Number of levels: 119)
##                           Estimate Est.Error l-95% CI u-95% CI Rhat Bulk_ESS Tail_ESS
## sd(Intercept)                 0.63      0.13     0.38     0.90 1.00     1425     2396
## sd(TrialType1)                1.06      0.14     0.80     1.38 1.00      728     1401
## cor(Intercept,TrialType1)    -0.06      0.26    -0.55     0.42 1.02      311      630
##
## Regression Coefficients:
##            Estimate Est.Error l-95% CI u-95% CI Rhat Bulk_ESS Tail_ESS
## Intercept     -0.07      0.15    -0.39     0.21 1.01      654     1438
## TrialType1    -3.70      0.17    -4.07    -3.39 1.00     1241     1959
##
## Draws were sampled using sampling(NUTS). For each parameter, Bulk_ESS
## and Tail_ESS are effective sample size measures, and Rhat is the potential
## scale reduction factor on split chains (at convergence, Rhat = 1).
```

##### Reactive Trial-Level A-cue Bias output

```
rea.model <- ProbeSlide.RESP ~ 0 + Intercept + TrialType + (1 + TrialType | sub.id)
# read in the model -- saves time compiling
rea.brms <- readRDS(paste0(in.path, "S1/reaTrialBias.rds"))
# if the model hasn't been run, uncomment and run, save, and update path to read in
# rea.brms <- brm(rea.model,
#                  data = rea.data,
#                  family= bernoulli(link = "logit"),
#                  cores = 6,
#                  sample_prior = TRUE,
#                  seed = 35)
```

```
# saveRDS(rea.brms, paste0(in.path,"S1/reaTrialBias.rds"))
s <- summary(rea.brms)
s
# store for later
rea.coef1 <- as.data.frame(coef(rea.brms))
rea.coef1 <- as.data.frame(cbind(rownames(rea.coef1),rea.coef1[,"sub.id.Estimate.Intercept"]))
colnames(rea.coef1) <- c("sub.id", "acue.coef")
```

```
##  Family: bernoulli
##   Links: mu = logit
## Formula: ProbeSlide.RESP ~ 0 + Intercept + TrialType + (1 + TrialType | sub.id)
##    Data: rea.data (Number of observations: 7098)
##   Draws: 4 chains, each with iter = 2000; warmup = 1000; thin = 1;
##          total post-warmup draws = 4000
##
## Multilevel Hyperparameters:
## ~sub.id (Number of levels: 120)
##                            Estimate Est.Error l-95% CI u-95% CI Rhat Bulk_ESS Tail_ESS
## sd(Intercept)                  0.74      0.12     0.52     0.98 1.00     1274     2284
## sd(TrialType1)                 0.78      0.12     0.56     1.01 1.01      690     1664
## cor(Intercept,TrialType1)      0.20      0.20    -0.24     0.56 1.01      543     1310
##
## Regression Coefficients:
##            Estimate Est.Error l-95% CI u-95% CI Rhat Bulk_ESS Tail_ESS
## Intercept     -0.18      0.13    -0.45     0.06 1.00      997     1724
## TrialType1    -3.21      0.13    -3.48    -2.97 1.00     1285     1717
##
## Draws were sampled using sampling(NUTS). For each parameter, Bulk_ESS
## and Tail_ESS are effective sample size measures, and Rhat is the potential
## scale reduction factor on split chains (at convergence, Rhat = 1).
```

##### A-cue Bias Trial-Level Measure vs. Summary Score

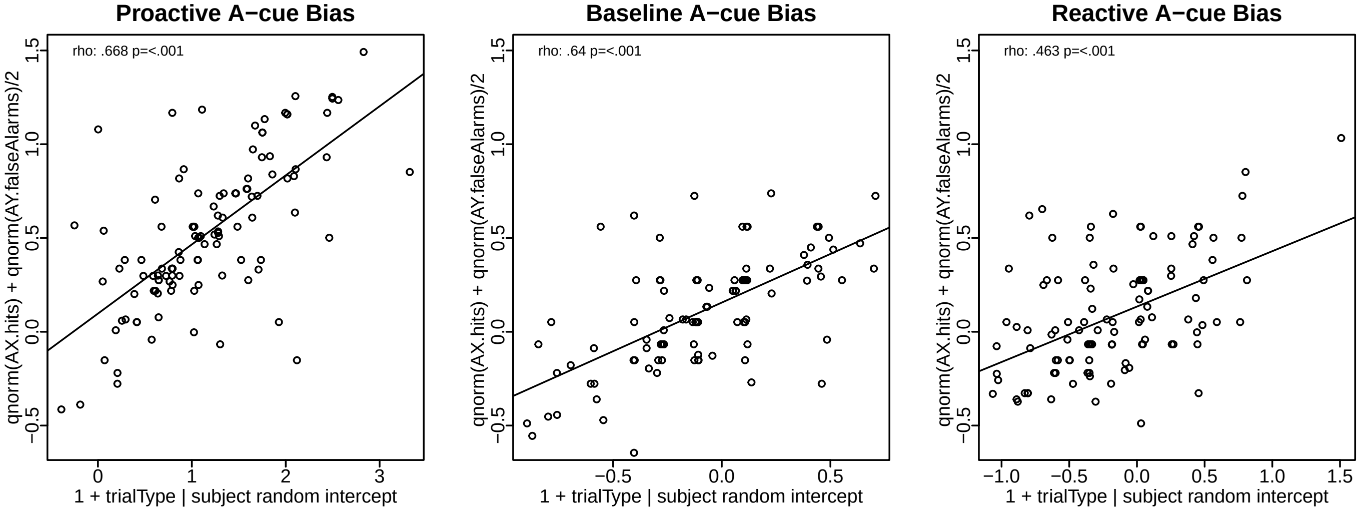


#### S1.3 Behavioral Replication of WMC preferential relationship with Proactive control

The following analyses replicates Lin et. al (2022) and utilizes posterior estimates from their analyses as priors. Note* This is Table 2 in the main paper

```
# create blank table to fill with output
rep.tbl <- data.frame(matrix(NA, nrow = 16, ncol=4))
colnames(rep.tbl) <- c("Fixed Effects", "Estimate (SD)", "95% CI", "Odds (SD)")
rep.tbl$`Fixed Effects` <- c("Intercept", sess.ids, "WMC", "Trial", paste0(sess.ids, ":WMC"), paste0(sess.ids, ":Trial"),

############################################################################################### set up the data
AXCPTD <- axcpt.tbl # get raw behavioral data
AXCPTD$session <- AXCPTD$Running

#subset relevant columns
AXCPTD <- AXCPTD[,c('sub.id', 'TrialType', 'ProbeSlide.RESP', 'ProbeSlide.CRESP', 'ProbeSlide.RTTime', 'session')]
#filter dataset to just have AX or AY trials and baseline or proactive mode.

AXCPTD <- AXCPTD %>%
  filter(TrialType == "AX" | TrialType == "AY") %>% #include only AX and AY trials
  mutate(TrialType = factor(TrialType, levels = c("AY", "AX"))) %>% #recode, make AY reference
  mutate(session = factor(session)) %>%
  mutate(sub.id = factor(sub.id)) %>%
  mutate(ProbeSlide.RESP = factor(ProbeSlide.RESP, levels = c(1, 2), labels = c("nontarget", "target"))) #recode into fac

contrasts(AXCPTD$TrialType) = contr.sum(2)
contrasts(AXCPTD$session) = contr.sum(3) # bas as ref; baseline=session1, pro=session2

AXCPTD_R <- AXCPTD %>% #change order of session to compare reactive to proactive
  mutate(session = factor(session, levels = c("reactiveBlockList", "proactiveBlockList", "baselineBlockList")))
contrasts(AXCPTD_R$session) = contr.sum(3) # set rea as ref so we can compare pro to rea, pro=session1; rea=session2

for (sid in 1:length(sub.ids)) {
  AXCPTD_R[which(AXCPTD_R$sub.id==sub.ids[sid]), "wk.mem"] <- wmc.tbl[which(wmc.tbl$sub.id==sub.ids[sid]), "composite.sco
}

for (sid in 1:length(sub.ids)) {
  AXCPTD[which(AXCPTD$sub.id==sub.ids[sid]), "wk.mem"] <- wmc.tbl[which(wmc.tbl$sub.id==sub.ids[sid]), "composite.score"]
}
################################################################################################### specifiy model and
m4D <- ProbeSlide.RESP ~ 0 + Intercept + session * wk.mem * TrialType + (1 + TrialType | sub.id)
# # set priors
prior <- c(set_prior("normal(-.36, .03)", class = "b", coef = "session1"),
           set_prior("normal(.51, .02)", class = "b", coef = "session2"),
           set_prior("normal(.16, .04)", class = "b", coef = "wk.mem"),
           set_prior("normal(-2.89, .05)", class = "b", coef = "TrialType1"),
           set_prior("normal(-.02, .03)", class = "b", coef = "session1:wk.mem"),
           set_prior("normal(.12, .02)", class = "b", coef = "session2:wk.mem"),
           set_prior("normal(-.28, .03)", class = "b", coef = "session1:TrialType1"),
           set_prior("normal(.34, .02)", class = "b", coef = "session2:TrialType1"),
           set_prior("normal(-.03, .05)", class = "b", coef = "wk.mem:TrialType1"),
           set_prior("normal(-.08, .03)", class = "b", coef = "session1:wk.mem:TrialType1"),
           set_prior("normal(0.00, .02)", class = "b", coef = "session2:wk.mem:TrialType1"),
           set_prior("normal(.24, .04)", class = "b", coef = "Intercept"))
#
# #run model (baseline as reference) if not already saved
# ACue.All.brm4D <- brm(m4D,
#                       data = AXCPTD,
#                       family= bernoulli(link = "logit"),
#                       cores = 6,
#                       prior = prior,
#                       sample_prior = TRUE,
#                       seed = 35)
# saveRDS(ACue.All.brm4D, paste0("S1/brms_basRef"))
# read in the saved model -- saves time compiling
ACue.All.brm4D <- readRDS(paste0(in.path,"S1/brms_basRef"))
```

```
############################################################################### Store results
s <- summary(ACue.All.brm4D)
s.basRef <- s
BasRef.est <- s$fixed
#compute probability of direction
BasRef.pd <- pd(ACue.All.brm4D)

#convert from logits to odds
odds <- posterior_samples(ACue.All.brm4D,"b") %>%
  mutate_all(exp) %>%
  gather() %>%
  group_by(key) %>%
    dplyr::summarize(mean = mean(value),
            sd   = sd(value),
            ll   = quantile(value, probs = .025),
            ul   = quantile(value, probs = .975))
# store for final table
BasRef.odds <- odds[c(1:12),]

###################################################################################################################
## fill table with results; session1=bas; session2=pro
want.rows <- c("Intercept", "Baseline", "Proactive", "WMC", "Trial", "Baseline:WMC", "Proactive:WMC",  "Baseline:Trial","
row.ids <- row.names(BasRef.est)
odds.ids <- BasRef.odds$key
for(rid in 1:length(want.rows)){
  rep.tbl[which(rep.tbl$`Fixed Effects`==want.rows[rid]),"Estimate (SD)"] <- paste0(doformat(BasRef.est[which(row.names(B
  rep.tbl[which(rep.tbl$`Fixed Effects`==want.rows[rid]),"95% CI"] <- paste0("[", doformat(BasRef.est[which(row.names(Bas
  rep.tbl[which(rep.tbl$`Fixed Effects`==want.rows[rid]),"Odds (SD)"] <- paste0( doformat(BasRef.odds[which(BasRef.odds$k
}
##################################################################################################################
#run model with comparing reactive (session1) to proactive (session2) as reference to get reactive results compared to pr
# need to set new priors from Lin et. al 2022 when Rea is the reference
m4DR <- ProbeSlide.RESP ~ 0 + Intercept + session * wk.mem * TrialType + (1 + TrialType | sub.id)

prior <- c(set_prior("normal(-.20, .03)", class = "b", coef = "session1"),
           set_prior("normal(.52, .03)", class = "b", coef = "session2"),
           set_prior("normal(.16, .04)", class = "b", coef = "wk.mem"),
           set_prior("normal(-2.89, .05)", class = "b", coef = "TrialType1"),
           set_prior("normal(-.11, .03)", class = "b", coef = "session1:wk.mem"),
           set_prior("normal(.11, .02)", class = "b", coef = "session2:wk.mem"),
           set_prior("normal(-.08, .03)", class = "b", coef = "session1:TrialType1"),
           set_prior("normal(.35, .02)", class = "b", coef = "session2:TrialType1"),
           set_prior("normal(-.03, .05)", class = "b", coef = "wk.mem:TrialType1"),
           set_prior("normal(-.07, .03)", class = "b", coef = "session1:wk.mem:TrialType1"),
           set_prior("normal(0.02, .02)", class = "b", coef = "session2:wk.mem:TrialType1"),
           set_prior("normal(.24, .04)", class = "b", coef = "Intercept"))
# # run model -- uncomment and run if not already saved
# ACue.All.brm4DR <- brm(m4DR,
#                       data = AXCPTD_R,
#                       family= bernoulli(link = "logit"),
#                       cores = 6,
#                       prior = prior,
#                       sample_prior = TRUE,
#                       seed = 35)
# saveRDS(ACue.All.brm4DR, paste0(in.path,"S1/brms_reaRef"))
# read in the data -- saves time compiling
ACue.All.brm4DR <- readRDS(paste0(in.path,"S1/brms_reaRef"))
############################################################################## store results
s <- summary(ACue.All.brm4DR)
s.reaRef <- s
ReaRef.est <- s$fixed
```

```r
#compute probability of direction
ReaRef.pd <- pd(ACue.All.brm4DR)

#convert from logits to odds
odds <- posterior_samples(ACue.All.brm4DR,"b") %>%
  mutate_all(exp) %>%
  gather() %>%
  group_by(key)  %>%
  dplyr::summarize(mean = mean(value),
            sd   = sd(value),
            ll   = quantile(value, probs = .025),
            ul   = quantile(value, probs = .975))


ReaRef.odds <- odds[c(1:12),]

# h2 <- hypothesis(ACue.All.brm4DR, "session1:wk.mem > 0") # check interaction effect > 0; pro is session 1 now
## fill the rest of final table
want.rows <- c("Reactive", "Reactive:WMC",  "Reactive:Trial","Reactive:WMC:Trial")
row.ids <- c("session1", "session1:wk.mem", "session1:TrialType1", "session1:wk.mem:TrialType1")
odds.ids <- c("b_session1", "b_session1:wk.mem", "b_session1:TrialType1", "b_session1:wk.mem:TrialType1")
for(rid in 1:length(want.rows)){
  rep.tbl[which(rep.tbl$`Fixed Effects`==want.rows[rid]),"Estimate (SD)"] <- paste0(doformat(ReaRef.est[which(row.names(R
  rep.tbl[which(rep.tbl$`Fixed Effects`==want.rows[rid]),"95% CI"] <- paste0("[", doformat(ReaRef.est[which(row.names(Rea
  rep.tbl[which(rep.tbl$`Fixed Effects`==want.rows[rid]),"Odds (SD)"] <- paste0( doformat(ReaRef.odds[which(ReaRef.odds$k
}
############################################################## cell specification for significance
#
#    Fixed Effects Estimate (SD)           95% CI    Odds (SD)   Odds (SD)
# 1      Intercept   .289 (.036)      [.22, .357] 1.336 (.048) 1.336 (.048) *
# 2            Bas  -.313 (.025)   [-.361, -.265]  .056 (.002)  .056 (.002) *
# 3            Pro   .581 (.018)     [.546, .617]  .732 (.018)  .732 (.018) *
# 4            Rea  -.256 (.025)   [-.303, -.208] 1.848 (.044)  .775 (.019) *
# 5            WMC    .12 (.028)     [.064, .175]  .732 (.018)  .732 (.018) *
# 6          Trial -2.886 (.043) [-2.971, -2.801]  .974 (.019)  .974 (.019) *
# 7        Bas:WMC   -.027 (.02)    [-.066, .012]  .957 (.018)  .957 (.018) NA
# 8        Pro:WMC   .086 (.015)     [.056, .116] 1.788 (.032) 1.788 (.032) *
# 9        Rea:WMC  -.093 (.019)    [-.13, -.056] 1.088 (.017)  .911 (.017) *
# 10     Bas:Trial  -.312 (.024)    [-.36, -.264] 1.457 (.026) 1.457 (.026) *
# 11     Pro:Trial   .376 (.018)     [.342, .411]  1.09 (.017)  1.09 (.017) *
# 12     Rea:Trial  -.053 (.025)     [-.1, -.004] 1.488 (.027)  .949 (.023) *
# 13     WMC:Trial  -.015 (.033)    [-.082, .049] 1.033 (.016) 1.033 (.016) NA
# 14 Bas:WMC:Trial  -.044 (.019)   [-.082, -.006] 1.128 (.032) 1.128 (.032) NA
# 15 Pro:WMC:Trial   .032 (.015)     [.003, .062]  .986 (.033)  .986 (.033) *
# 16 Rea:WMC:Trial  -.044 (.019)   [-.081, -.006] 1.088 (.017)  .958 (.018) *
inds <- c(1:6, 8:12,14:16) # significant CI's]

h1 <- hypothesis(ACue.All.brm4D, "session2:wk.mem > 0") # check interaction effect > 0  ## test prob of hyp and store

h1.plot <- plot(h1)[[1]] +
            theme(strip.text = element_blank(), xlim=c(0,.2))      # store hyp test as a plot
```

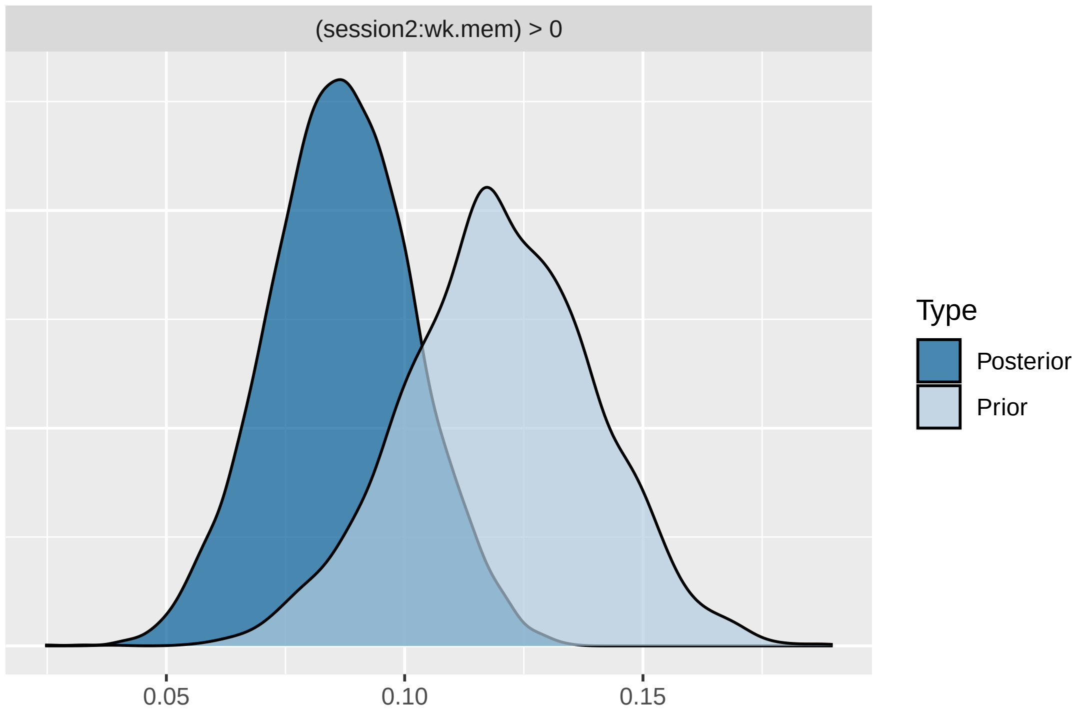


**Baseline Reference Model Output**

```
##  Family: bernoulli
##   Links: mu = logit
## Formula: ProbeSlide.RESP ~ 0 + Intercept + session * wk.mem * TrialType + (1 + TrialType | sub.id)
##    Data: AXCPTD (Number of observations: 20712)
##   Draws: 4 chains, each with iter = 2000; warmup = 1000; thin = 1;
##          total post-warmup draws = 4000
##
## Multilevel Hyperparameters:
## ~sub.id (Number of levels: 118)
##                              Estimate Est.Error l-95% CI u-95% CI Rhat Bulk_ESS Tail_ESS
## sd(Intercept)                    0.64      0.06     0.53     0.77 1.00     1304     2683
## sd(TrialType1)                   0.73      0.07     0.61     0.87 1.00      940     1506
## cor(Intercept,TrialType1)        0.24      0.12     0.00     0.46 1.00      643     1175
##
## Regression Coefficients:
##                               Estimate Est.Error l-95% CI u-95% CI Rhat Bulk_ESS Tail_ESS
## Intercept                         0.29      0.03     0.22     0.36 1.00     2333     2568
## session1                         -0.31      0.02    -0.36    -0.26 1.00     5626     3324
## session2                          0.58      0.02     0.55     0.61 1.00     6176     3186
## wk.mem                            0.12      0.03     0.07     0.17 1.00     1975     3156
## TrialType1                       -2.88      0.04    -2.96    -2.80 1.00     2344     2982
## session1:wk.mem                  -0.03      0.02    -0.06     0.01 1.00     4008     3208
## session2:wk.mem                   0.09      0.01     0.06     0.12 1.00     4929     3197
## session1:TrialType1              -0.31      0.02    -0.36    -0.27 1.00     5375     3163
## session2:TrialType1               0.38      0.02     0.34     0.41 1.00     5297     2832
## wk.mem:TrialType1                -0.02      0.03    -0.08     0.05 1.00     1493     2286
## session1:wk.mem:TrialType1       -0.04      0.02    -0.08    -0.01 1.00     4389     3291
## session2:wk.mem:TrialType1        0.03      0.01     0.00     0.06 1.00     4281     2967
##
## Draws were sampled using sampling(NUTS). For each parameter, Bulk_ESS
## and Tail_ESS are effective sample size measures, and Rhat is the potential
## scale reduction factor on split chains (at convergence, Rhat = 1).
```

##### Reactive Reference Model Output

```
##  Family: bernoulli
##   Links: mu = logit
## Formula: ProbeSlide.RESP ~ 0 + Intercept + session * wk.mem * TrialType + (1 + TrialType | sub.id)
##    Data: AXCPTD_R (Number of observations: 20712)
##   Draws: 4 chains, each with iter = 2000; warmup = 1000; thin = 1;
##          total post-warmup draws = 4000
##
## Multilevel Hyperparameters:
## ~sub.id (Number of levels: 118)
##                           Estimate Est.Error l-95% CI u-95% CI Rhat Bulk_ESS Tail_ESS
## sd(Intercept)                 0.67      0.06     0.56     0.81 1.00      978     1859
## sd(TrialType1)                0.73      0.06     0.61     0.86 1.00     1016     1731
## cor(Intercept,TrialType1)     0.21      0.12    -0.03     0.43 1.01      687     1285
##
## Regression Coefficients:
##                            Estimate Est.Error l-95% CI u-95% CI Rhat Bulk_ESS Tail_ESS
## Intercept                      0.28      0.04     0.21     0.35 1.00     2317     2670
## session1                      -0.25      0.02    -0.30    -0.21 1.00     5573     2861
## session2                       0.61      0.02     0.56     0.66 1.00     5383     2295
## wk.mem                         0.08      0.03     0.03     0.14 1.00     1793     2572
## TrialType1                    -2.89      0.04    -2.97    -2.80 1.00     2393     2682
## session1:wk.mem               -0.05      0.02    -0.09    -0.02 1.00     4722     2940
## session2:wk.mem                0.07      0.02     0.04     0.10 1.00     5380     3082
## session1:TrialType1           -0.06      0.02    -0.10    -0.01 1.00     5413     3146
## session2:TrialType1            0.40      0.02     0.36     0.44 1.00     4358     2684
## wk.mem:TrialType1              0.02      0.03    -0.05     0.08 1.00     1631     2416
## session1:wk.mem:TrialType1    -0.02      0.02    -0.06     0.02 1.00     4620     2963
## session2:wk.mem:TrialType1     0.01      0.02    -0.02     0.04 1.00     4514     3111
##
## Draws were sampled using sampling(NUTS). For each parameter, Bulk_ESS
## and Tail_ESS are effective sample size measures, and Rhat is the potential
## scale reduction factor on split chains (at convergence, Rhat = 1).
```

**Combined Model Output** Note* this is Table 2 in the main text.

| Fixed Effects | Estimate (SD) | 95% CI | Odds (SD) |
|---|---|---|---|
| Intercept | .287 (.035) | [.219, .356]* | 1.333 (.047) |
| Baseline | -.312 (.025) | [-.362, -.264]* | .056 (.002) |
| Proactive | .58 (.018) | [.545, .614]* | .732 (.018) |
| Reactive | -.254 (.024) | [-.3, -.205]* | .776 (.019) |
| WMC | .12 (.028) | [.065, .175]* | .733 (.017) |
| Trial | -2.885 (.041) | [-2.964, -2.804]* | .974 (.018) |
| **Baseline:WMC** | **-.027 (.019)** | **[-.064, .01]** | **.957 (.018)** |
| **Proactive:WMC** | **.087 (.015)** | **[.057, .116]*** | **1.787 (.032)** |
| **Reactive:WMC** | **-.054 (.019)** | **[-.092, -.018]*** | **.947 (.018)** |
| Baseline:Trial | -.311 (.024) | [-.358, -.266]* | 1.457 (.026) |
| Proactive:Trial | .376 (.018) | [.341, .413]* | 1.091 (.016) |
| Reactive:Trial | -.056 (.024) | [-.102, -.011]* | .946 (.023) |
| WMC:Trial | -.015 (.033) | [-.08, .048] | 1.034 (.015) |
| Baseline:WMC:Trial | -.044 (.019) | [-.082, -.005]* | 1.128 (.032) |
| Proactive:WMC:Trial | .034 (.015) | [.004, .062]* | .985 (.032) |
| Reactive:WMC:Trial | -.018 (.02) | [-.057, .02]* | .982 (.019) |

### Supplemental 2: Brain analyses

Descriptive statistics were analyzed using WRS2 and DescTools packages for robust statistics. Mean and standard errors (SEM) trimmed at 0.1.

##### S2.1 Example GLM call:

Example of full AFNI GLM call. Note hashtags are used to help word wrap the command within the page width. Remove before running any command.

```
# 3dDeconvolve \
#  -local_times \
#  -x1D_stop \
#  -GOFORIT 5 \
#  -input 'lpi_scale_blur4_tfMRI_AxcptBas1_AP.nii.gz
#   lpi_scale_blur4_tfMRI_AxcptBas2_PA.nii.gz' \
#  -polort A \
#  -float \
#  -censor movregs_FD_mask.txt \
#  -num_stimts 3 \
#  -stim_times_AM1 1 f1031ax_Axcpt_baseline_block.txt 'dmBLOCK(1)'
#  -stim_label 1 ON_BLOCKS \
#  -stim_times 2 f1031ax_Axcpt_baseline_blockONandOFF.txt 'TENTzero(0,16.8,8)'
#  -stim_label 2 ON_blockONandOFF \
#  -stim_times 3 f1031ax_Axcpt_baseline_allTrials.txt 'TENTzero(0,21.6,10)'
#  -stim_label 3 ON_TRIALS \
#  -ortvec motion_demean_baseline.1D movregs \
#  -x1D X.xmat.1D \
#  -xjpeg X.jpg \
#  -nobucket
#
# 3dREMLfit \
#  -matrix X.xmat.1D \
#  -GOFORIT 5 \
#  -input 'lpi_scale_blur4_tfMRI_AxcptBas1_AP.nii.gz
#   lpi_scale_blur4_tfMRI_AxcptBas2_PA.nii.gz' \
#  -Rvar stats_var_f1031ax_REML.nii.gz \
#  -Rbuck STATS_f1031ax_REML.nii.gz \
#  -fout \
#  -tout \
#  -nobout \
#  -verb
```

##### S2.2 Visualization of Different Conjunction Tests

Surface plots rendering significant uncorrected robust (Yuen, trimmed means) t-stat values >2

Each set of brain plots visualizes parcels which pass the conjunction test, first separately, and then as a final conjunction parcel set. Parcels needed to pass these 3 tests in the Proactive condition in order to be included, but Baseline and Reactive plots are included to show how these effects are most prominant in the Proactive condition. Color Legend Significant T Values

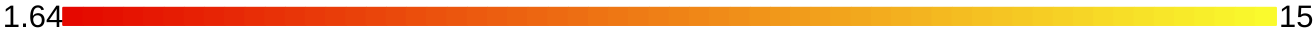


**Test 1: A-cue activation >0**

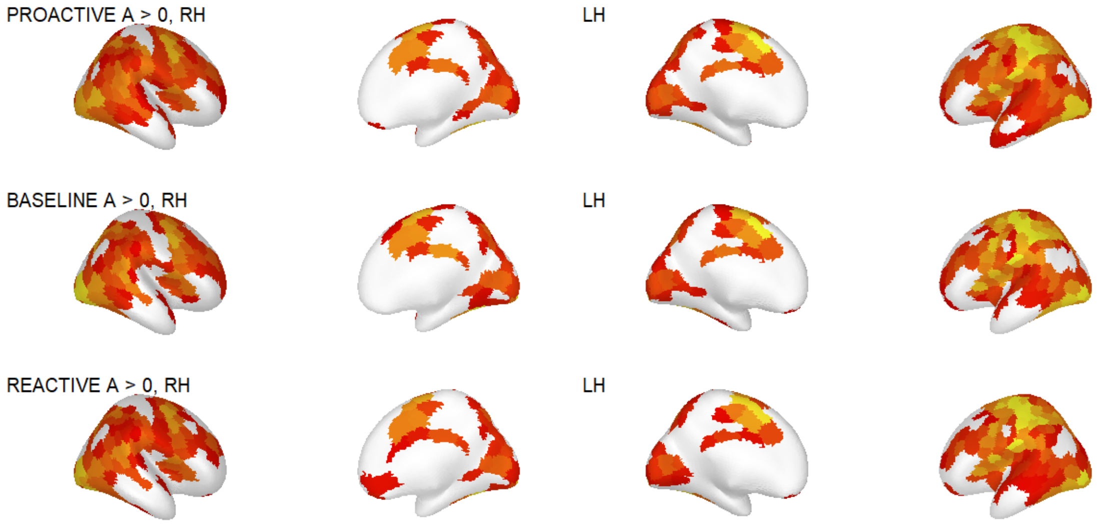


**Test 2: A>B cue activation >0**

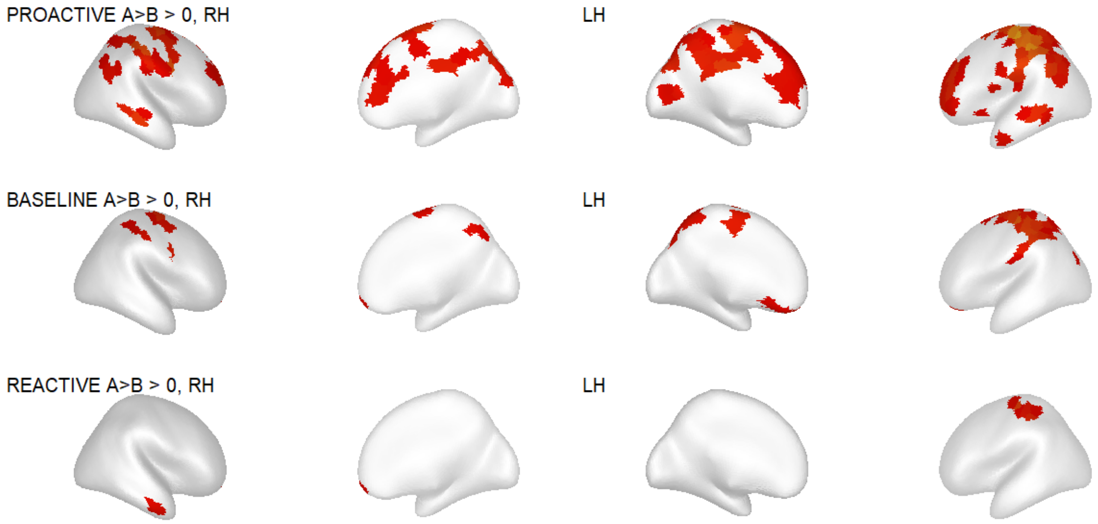


**Test 3: A >B-cue activation greater in Proactive session compared to Baseline and Reactive i.e. [A-cue - B-cue]Proactive - [A-cue - B-cue]Baseline >0 or [A-cue -B-cue]Reactive >0**

Note* There are parcels with B >A-cue activation since this is a comparison of contrasts where the Proactive session has *less* B >A-cue activation than Reactive or Baseline. The inclusion of Test 2 (A >B-cue activation is >0) ensures that only parcels selective to A-cue activation are included.

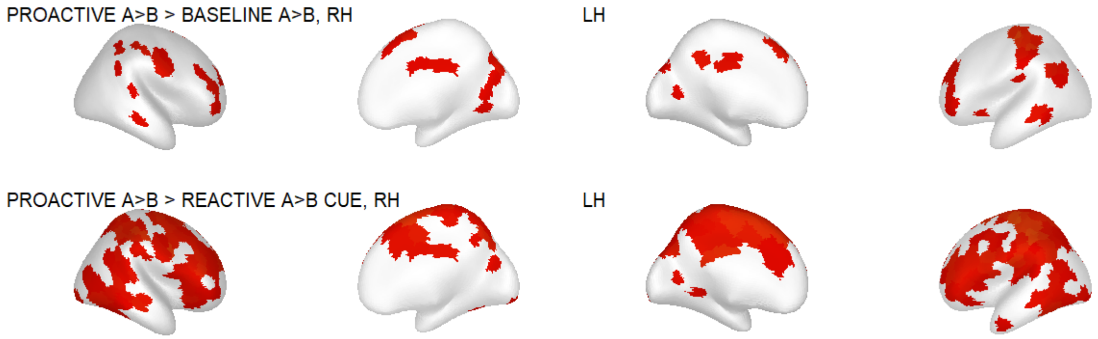

**Final Parcellation:**
Parcels that passed all 3 tests are now included in yellow.

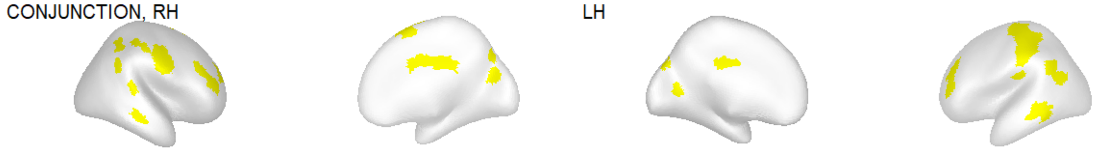


##### S2.3 Brain-Behavior Correlations

Conjunction analysis parcels were subsequently tested for correlations between A-cue activation and A-cue Bias as well as working memory capacity.

**A-cue Bias and A-cue Activation correlations:**

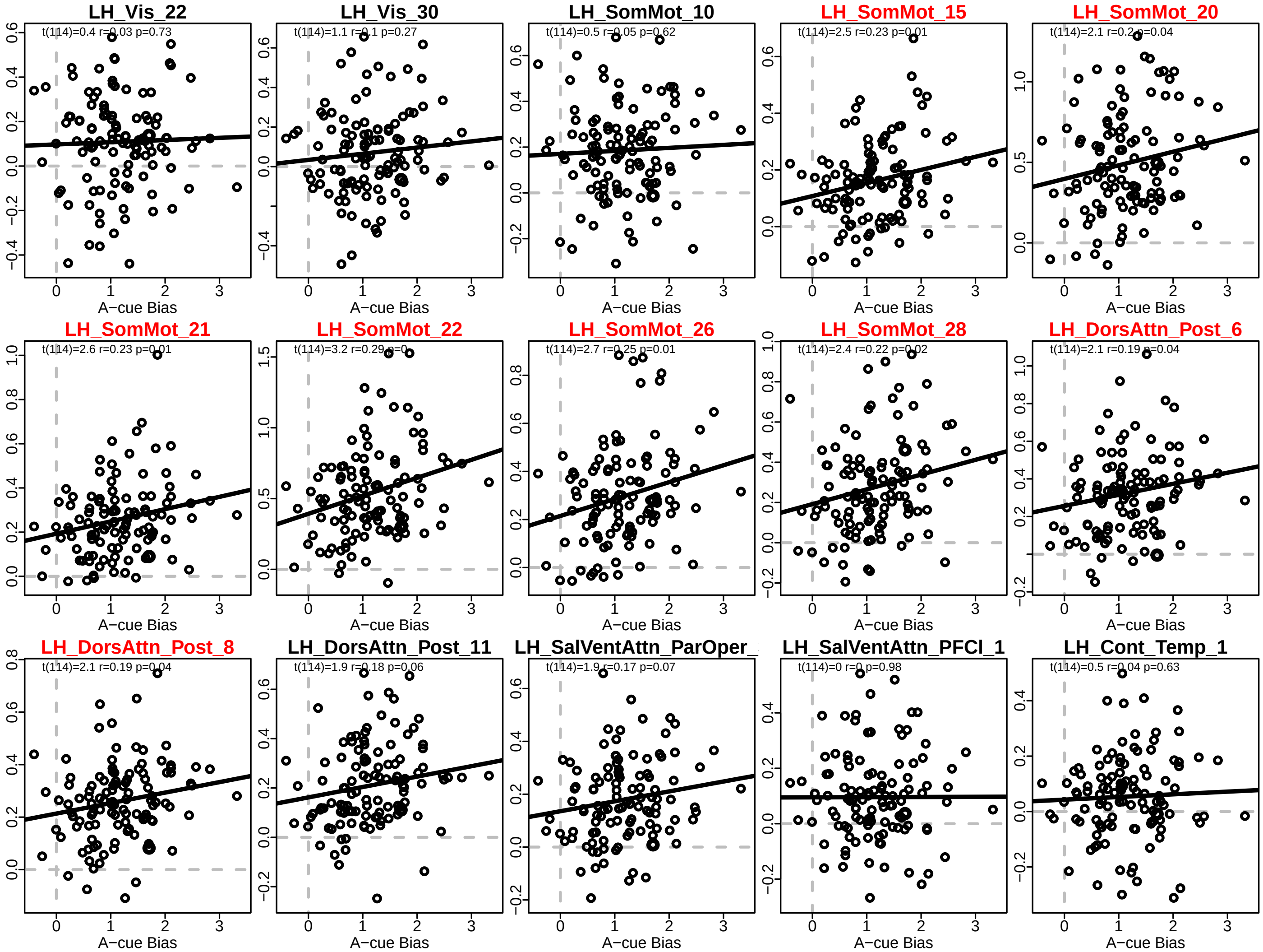

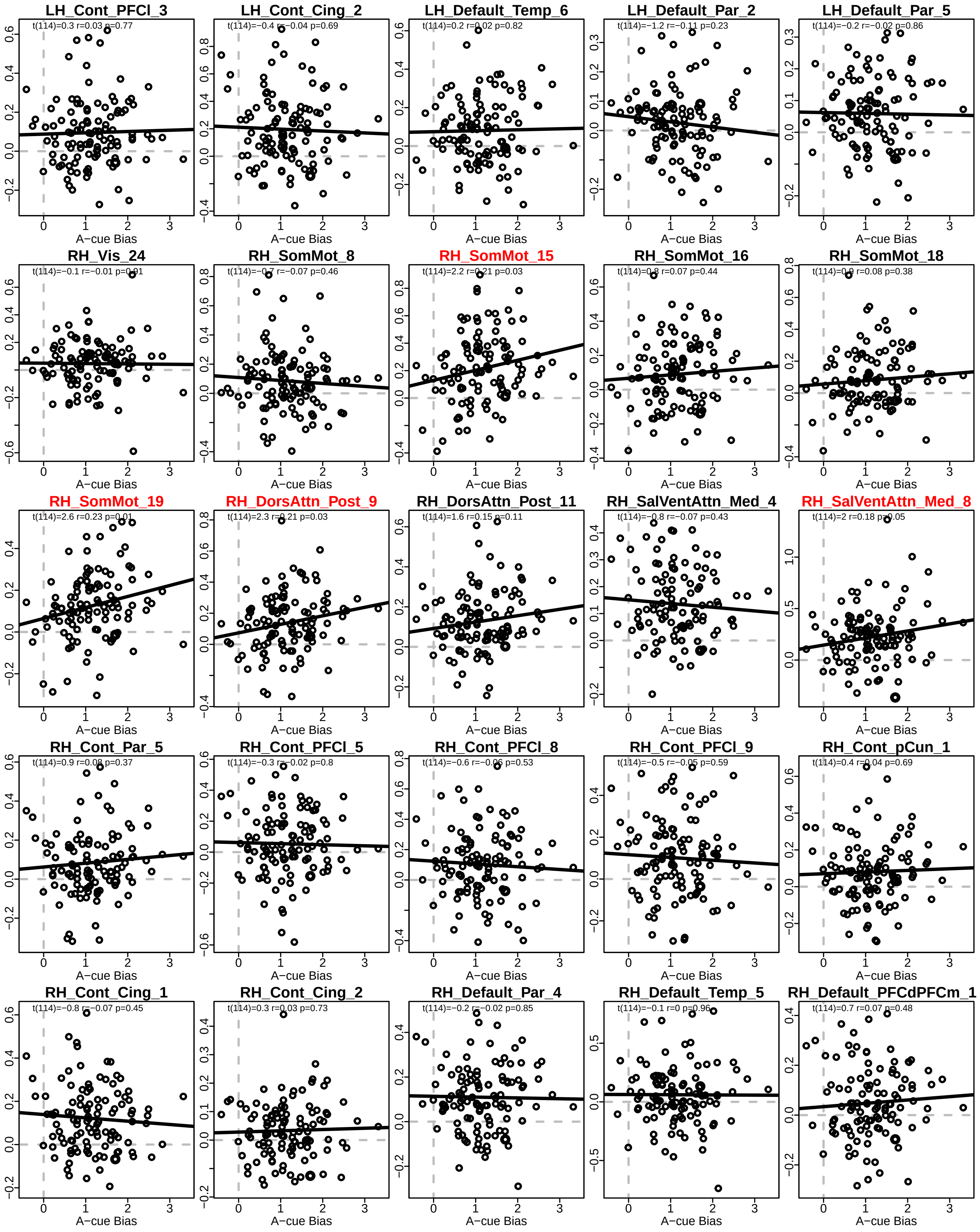
LH_Cont_PFCl_3
t(114)=0.3 r=0.03 p=0.77
LH_Cont_Cing_2
t(114)=−0.4 r=−0.04 p=0.69
LH_Default_Temp_6
t(114)=0.2 r=0.02 p=0.82
LH_Default_Par_2
t(114)=−1.2 r=−0.11 p=0.23
LH_Default_Par_5
t(114)=−0.2 r=−0.02 p=0.86
RH_Vis_24
t(114)=−0.1 r=−0.01 p=0.91
RH_SomMot_8
t(114)=−0.7 r=−0.07 p=0.46
RH_SomMot_15
t(114)=2.2 r=0.21 p=0.03
RH_SomMot_16
t(114)=0.8 r=0.07 p=0.44
RH_SomMot_18
t(114)=0.9 r=0.08 p=0.38
RH_SomMot_19
t(114)=2.6 r=0.23 p=0.01
RH_DorsAttn_Post_9
t(114)=2.3 r=0.21 p=0.03
RH_DorsAttn_Post_11
t(114)=1.6 r=0.15 p=0.11
RH_SalVentAttn_Med_4
t(114)=−0.8 r=−0.07 p=0.43
RH_SalVentAttn_Med_8
t(114)=2 r=0.18 p=0.05
RH_Cont_Par_5
t(114)=0.9 r=0.08 p=0.37
RH_Cont_PFCl_5
t(114)=−0.3 r=−0.02 p=0.8
RH_Cont_PFCl_8
t(114)=−0.6 r=−0.06 p=0.53
RH_Cont_PFCl_9
t(114)=−0.5 r=−0.05 p=0.59
RH_Cont_pCun_1
t(114)=0.4 r=0.04 p=0.69
RH_Cont_Cing_1
t(114)=−0.8 r=−0.07 p=0.45
RH_Cont_Cing_2
t(114)=0.3 r=0.03 p=0.73
RH_Default_Par_4
t(114)=−0.2 r=−0.02 p=0.85
RH_Default_Temp_5
t(114)=−0.1 r=0 p=0.96
RH_Default_PFCdPFCm_1
t(114)=0.7 r=0.07 p=0.48
A−cue Bias

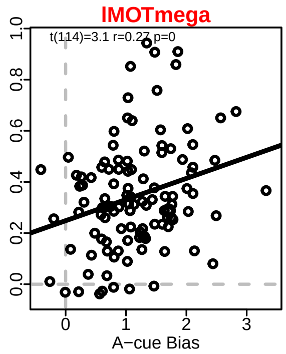


**WMC and A-cue Activation correlations:**

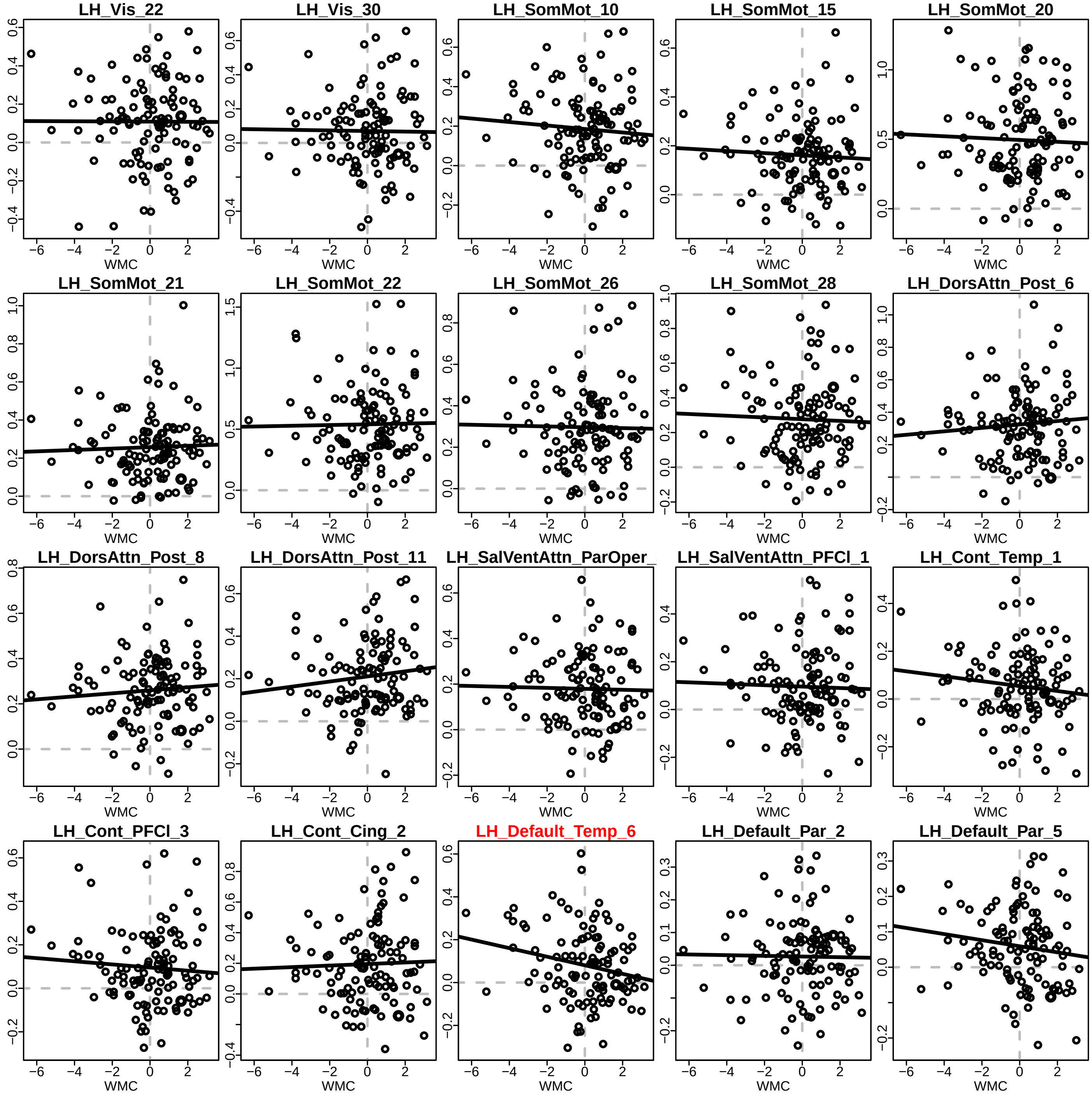

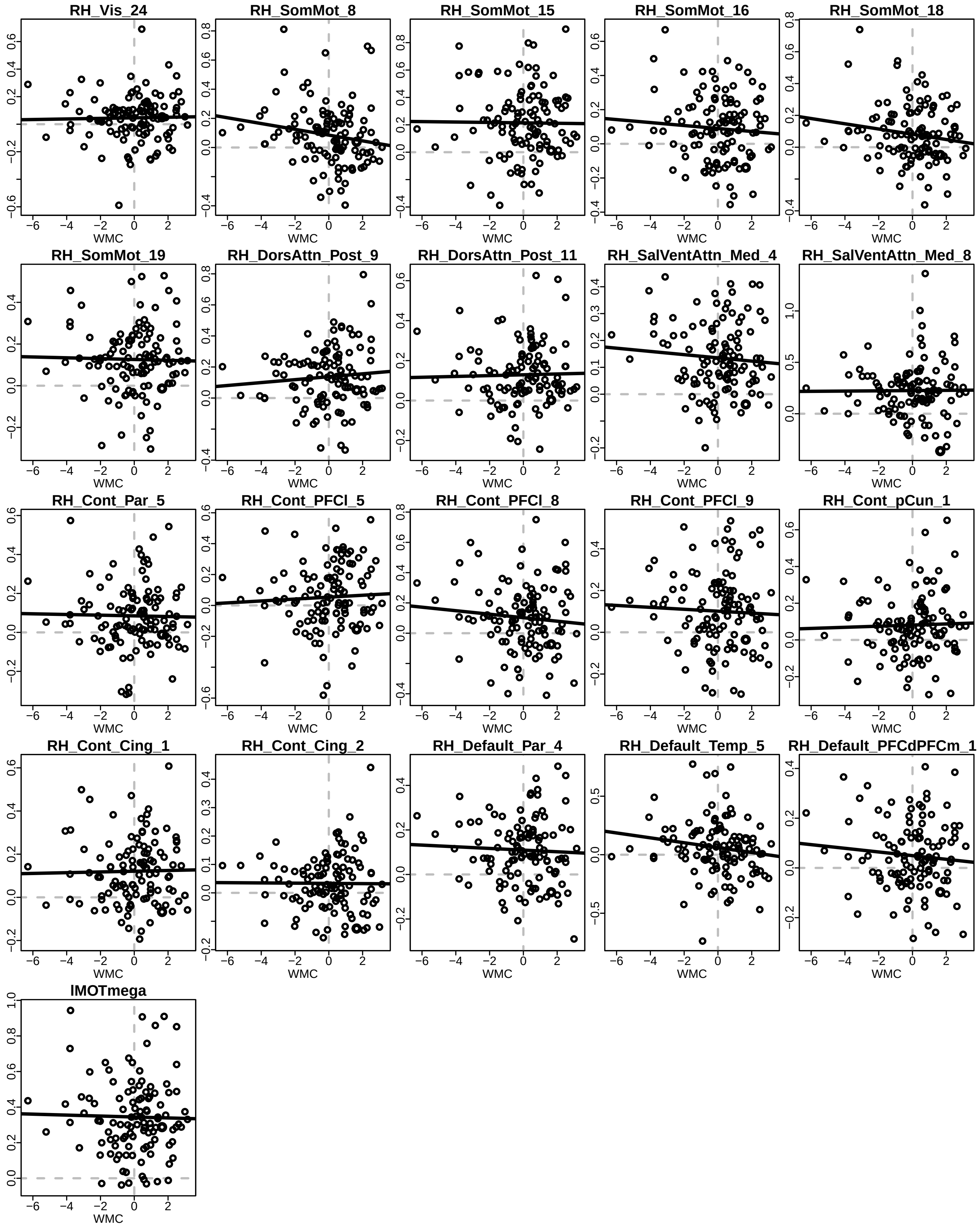
RH_Vis_24
RH_SomMot_8
RH_SomMot_15
RH_SomMot_16
RH_SomMot_18
RH_SomMot_19
RH_DorsAttn_Post_9
RH_DorsAttn_Post_11
RH_SalVentAttn_Med_4
RH_SalVentAttn_Med_8
RH_Cont_Par_5
RH_Cont_PFCl_5
RH_Cont_PFCl_8
RH_Cont_PFCl_9
RH_Cont_pCun_1
RH_Cont_Cing_1
RH_Cont_Cing_2
RH_Default_Par_4
RH_Default_Temp_5
RH_Default_PFCdPFCm_1
IMOTmega
WMC

##### S2.4 Conjunction Analysis Results

**Conjunction Table:** Parcels that passed the conjunction had A-cue activation >0, more A >B-cue activation in Proactive relative to Baseline and Reactive sessions. Parcels in red were also significantly correlated with A-cue behavioral index and the rDLPFC parcel in green is correlated with WMC.

| ROI | Proactive Acue > 0 T | Proactuve Acue > 0 P | A>B Proactive>Baseline T | A>B Proactive>Baseline P | A>B Proactive>Reactive T | A>B, Proactive>Reactive P |
|---|---|---|---|---|---|---|
| LHVis22 | 5.611 | <.001 | 1.832 | .035 | 1.980 | .025 |
| LHVis30 | 3.404 | <.001 | 1.664 | .05 | 2.736 | .004 |
| LHSomMot10 | 10.337 | <.001 | 2.059 | .021 | 3.017 | .002 |
| LHSomMot15 | 12.143 | <.001 | 2.509 | .007 | 2.714 | .004 |
| LHSomMot20 | 16.483 | <.001 | 3.834 | <.001 | 4.308 | <.001 |
| LHSomMot21 | 16.784 | <.001 | 3.232 | .001 | 4.496 | <.001 |
| LHSomMot22 | 18.256 | <.001 | 5.209 | <.001 | 5.672 | <.001 |
| LHSomMot26 | 16.523 | <.001 | 3.612 | <.001 | 5.213 | <.001 |
| LHSomMot28 | 11.861 | <.001 | 3.041 | .002 | 4.063 | <.001 |
| LHDorsAttnPost6 | 15.587 | <.001 | 1.913 | .029 | 4.992 | <.001 |
| LHDorsAttnPost8 | 19.889 | <.001 | 3.931 | <.001 | 5.305 | <.001 |
| LHDorsAttnPost11 | 13.537 | <.001 | 2.762 | .003 | 4.445 | <.001 |
| LHSalVentAttnParOper3 | 12.508 | <.001 | 1.792 | .038 | 3.301 | .001 |
| LHSalVentAttnPFCl1 | 6.060 | <.001 | 2.304 | .012 | 3.029 | .002 |
| LHContTemp1 | 4.188 | <.001 | 1.702 | .046 | 2.375 | .01 |
| LHContPFCl3 | 6.004 | <.001 | 1.889 | .031 | 2.995 | .002 |
| LHContCing2 | 7.569 | <.001 | 2.267 | .013 | 3.703 | <.001 |
| LHDefaultTemp6 | 4.648 | <.001 | 2.323 | .011 | 2.242 | .014 |
| LHDefaultPar2 | 2.408 | .009 | 1.855 | .033 | 2.067 | .021 |
| LHDefaultPar5 | 5.557 | <.001 | 2.708 | .004 | 3.334 | .001 |
| RHVis24 | 3.413 | <.001 | 1.795 | .038 | 1.925 | .029 |
| RHSomMot8 | 4.584 | <.001 | 1.741 | .043 | 2.459 | .008 |
| RHSomMot15 | 8.645 | <.001 | 2.088 | .02 | 4.514 | <.001 |
| RHSomMot16 | 4.533 | <.001 | 2.414 | .009 | 2.387 | .01 |
| RHSomMot18 | 4.651 | <.001 | 1.851 | .034 | 1.946 | .027 |
| RHSomMot19 | 9.383 | <.001 | 2.637 | .005 | 4.171 | <.001 |
| RHDorsAttnPost9 | 8.224 | <.001 | 2.669 | .005 | 4.343 | <.001 |
| RHDorsAttnPost11 | 9.383 | <.001 | 1.856 | .033 | 3.459 | <.001 |
| RHSalVentAttnMed4 | 10.103 | <.001 | 2.581 | .006 | 4.209 | <.001 |
| RHSalVentAttnMed8 | 9.537 | <.001 | 2.296 | .012 | 4.884 | <.001 |
| RHContPar5 | 5.696 | <.001 | 2.485 | .007 | 3.344 | .001 |
| RHContPFCl5 | 2.912 | .002 | 2.502 | .007 | 2.163 | .017 |
| RHContPFCl8 | 5.020 | <.001 | 2.088 | .02 | 3.138 | .001 |
| RHContPFCl9 | 5.429 | <.001 | 1.671 | .049 | 3.056 | .001 |
| RHContpCun1 | 4.731 | <.001 | 2.135 | .018 | 2.872 | .003 |
| RHContCing1 | 7.924 | <.001 | 2.072 | .021 | 2.464 | .008 |
| RHContCing2 | 3.411 | <.001 | 1.833 | .035 | 2.095 | .019 |
| RHDefaultPar4 | 7.798 | <.001 | 1.750 | .042 | 2.491 | .007 |
| RHDefaultTemp5 | 2.759 | .003 | 1.902 | .03 | 2.919 | .002 |
| RHDefaultPFCdPFCm13 | 3.595 | <.001 | 2.494 | .007 | 4.336 | <.001 |
| lMOTmega | 17.852 | <.001 | 4.036 | <.001 | 5.460 | <.001 |

Conjunction analysis parcels in yellow that had A-cue activation >0, more A >B-cue activation in Proactive relative to Baseline and Reactive sessions. Parcels in red were also significantly correlated with A-cue behavioral index and the rDLPFC parcel in green is correlated with WMC.

Note* This is highlighting the same parcels as panel B in Figure 3 of the main text, but the outlined version was made in WorkBench. The version in the supplemental is made using code frome (`https://osf.io/u9e82`).

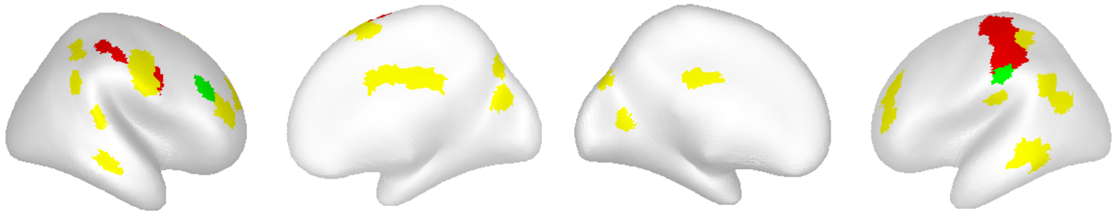

##### S2.5 A- and B-cue Timecourses Using Candidate Parcels

Example of full cue and probe timecourses for AX and BY trial types using the 40 significant parcels from the conjunction analysis. As can be seen, for these ROIs, the estimated activation timecourse (on AX and BY trial types) shows prominent activation during both cue and probe periods, along with differentiation between A-cue and B-cue activity, and among the three task conditions. Note* This is panel A in Figure 3 of the main text.

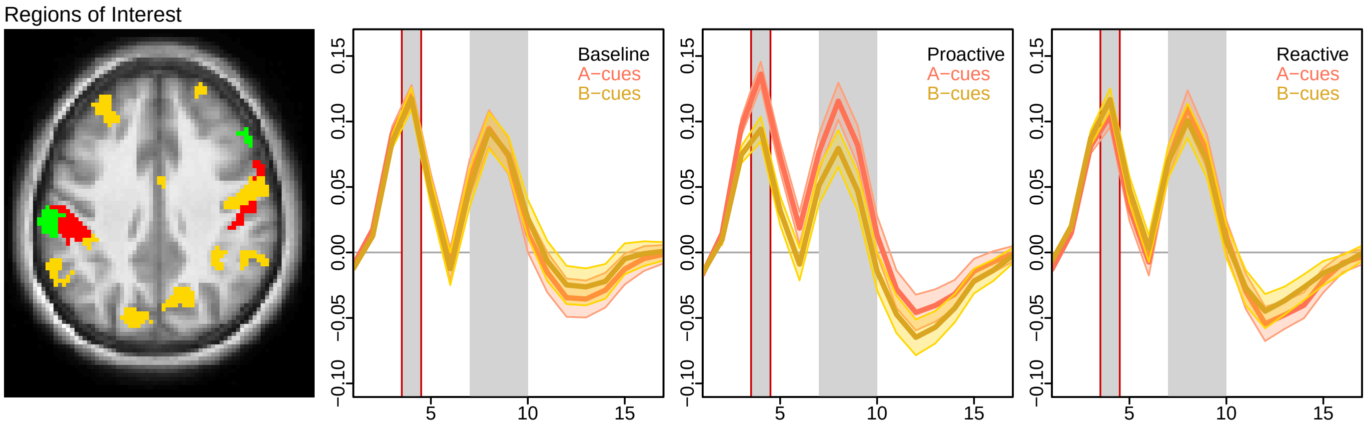

### Supplemental 3: Within-Person Mediation

#### S3.1: A-cue activation and A-cue Bias change from Baseline to Proactive

First we wanted to show how both in activation and in bias, their is more in Proactive compared to Baseline. The following values are from the combined lMOT parcel (LH SomMot 22 and LH SomMOt 26). The rest of this supplemental showcases how we ended up with these 2 contiguous regions.

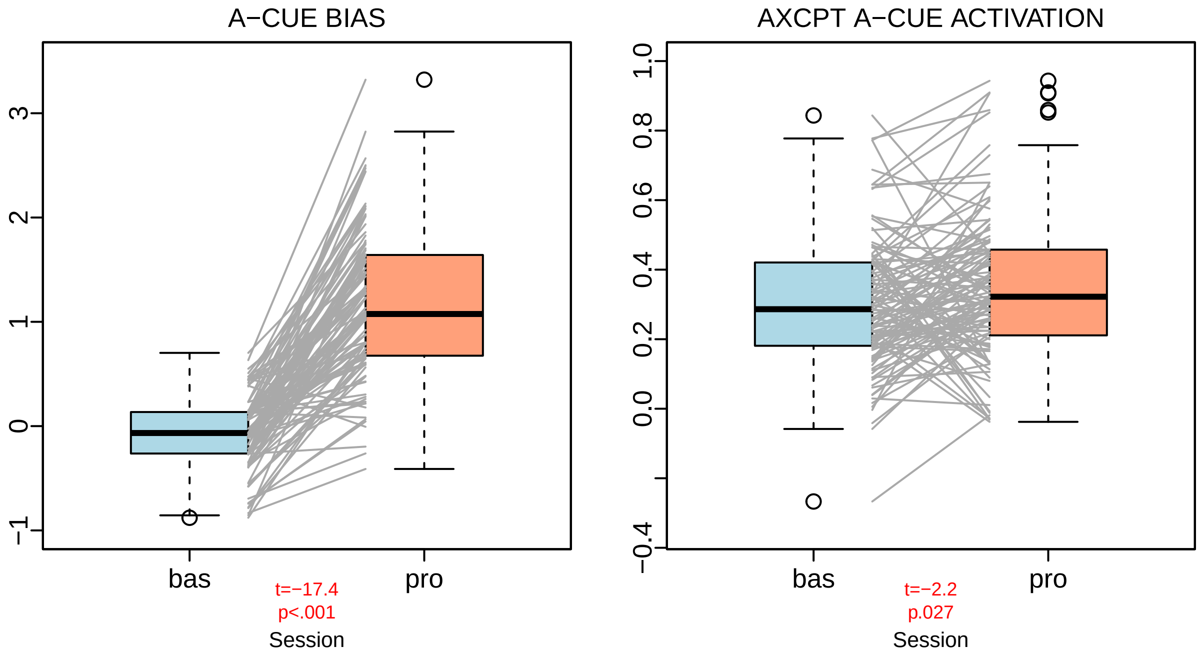


#### S3.2 Within Mediation

Of the parcels that correlated with A-cue bias, LH_SomMot_22 and LH_SomMot_26 also met the requirements for a within mediation analysis protocol from Judd (2011). Both parcels correlated with A-cue behavioral bias in Proactive and Baseline sessions independently and Proactive-Baseline difference in A-cue behavioral bias correlated with Proactive-Baseline A-cue activity. Finally, A-cue activity can be said to mediate (H.diff) or moderate (H.sum.centered) differences in A-cue bias if the neural activity difference between Proactive and Baseline predictive of behavioral measure difference (i.e. Y.diff $\sim$ H.sum + H.diff); Where Y=behavioral measure i.e. A-cue bias; Y.diff = Proactive-Baseline behavioral A-cue bias; H=neural measure i.e. knot 4 A-cue activity; H.diff = Proactive-Baseline neural A-cue activity

##### S3.2a: Is A-cue neural activity related to A-cue bias?

First, we analyzed each of the parcels with significant correlation between A-cue bias and Proactive A-cue activation for the same relationship in Baseline and Proactive-Baseline individually and if the neural activation is a predictor of behavior within the respective sessions:

**Correlations between A-cue Bias and Activation for both Baseline, Proactive, and Proactive-Baseline for parcels of interest:**
Only LH SomMot 22, LH SomMot 26, and their combined parcellation had significant correlations in all 3 conditions.

##### S3.2b: Is A-cue neural activity predictive of A-cue bias?

**Linear Regression Output:**
The parcels that had significant relationships were subsequently tested for predictive relationships in accordance with Judd et al., 2011.

Table 2: Correlations

| ROI | Baseline r | Baseline p-value | Proactive r | Proactive p-value | Proactive-Baseline r | Proactive-Baseline p-value |
|---|---|---|---|---|---|---|
| LHSomMot15 | 0.238 | .011 | 0.230 | .013 | 0.144 | .13 |
| LHSomMot20 | 0.181 | .055 | 0.195 | .036 | 0.153 | .108 |
| LHSomMot21 | 0.213 | .023 | 0.234 | .012 | 0.136 | .154 |
| LHSomMot22 | 0.238 | .011 | 0.286 | .002 | 0.215 | .023 |
| LHSomMot26 | 0.196 | .037 | 0.249 | .007 | 0.232 | .014 |
| LHSomMot28 | 0.157 | .095 | 0.218 | .018 | 0.152 | .11 |
| LHDorsAttnPost6 | 0.178 | .058 | 0.189 | .043 | 0.102 | .287 |
| LHDorsAttnPost8 | 0.230 | .014 | 0.190 | .041 | 0.100 | .296 |
| RHSomMot15 | 0.113 | .232 | 0.205 | .027 | 0.195 | .039 |
| RHSomMot19 | 0.248 | .008 | 0.233 | .012 | 0.091 | .338 |
| RHDorsAttnPost9 | 0.329 | <.001 | 0.208 | .025 | 0.072 | .453 |
| RHSalVentAttnMed8 | 0.233 | .012 | 0.184 | .048 | 0.109 | .255 |
| lMOTmega | 0.205 | .028 | 0.275 | .003 | 0.237 | .012 |

```
## [1] "LH_SomMot_22 Output: "
##
## Call:
## lm(formula = Y.diff ~ H.sum.centered + H.diff)
##
## Residuals:
##      Min       1Q   Median       3Q      Max
## -1.59435 -0.39217 -0.04088  0.39967  1.85005
##
## Coefficients:
##                Estimate Std. Error t value Pr(>|t|)
## (Intercept)     1.17207    0.06536  17.934   <2e-16 ***
## H.sum.centered  0.13434    0.13232   1.015   0.3122
## H.diff          0.42505    0.20420   2.082   0.0397 *
## ---
## Signif. codes:  0 '***' 0.001 '**' 0.01 '*' 0.05 '.' 0.1 ' ' 1
##
## Residual standard error: 0.6658 on 109 degrees of freedom
##   (6 observations deleted due to missingness)
## Multiple R-squared:  0.05528, Adjusted R-squared:  0.03795
## F-statistic: 3.189 on 2 and 109 DF,  p-value: 0.04508
##
## [1] "LH_SomMot_26 Output: "
##
## Call:
## lm(formula = Y.diff ~ H.sum.centered + H.diff)
##
## Residuals:
##      Min       1Q   Median       3Q      Max
## -1.59792 -0.38244 -0.03275  0.40642  1.74065
##
## Coefficients:
##                Estimate Std. Error t value Pr(>|t|)
## (Intercept)     1.17540    0.06415  18.324   <2e-16 ***
## H.sum.centered  0.21644    0.19569   1.106   0.2712
## H.diff          0.73037    0.30392   2.403   0.0179 *
## ---
## Signif. codes:  0 '***' 0.001 '**' 0.01 '*' 0.05 '.' 0.1 ' ' 1
##
## Residual standard error: 0.6626 on 109 degrees of freedom
##   (6 observations deleted due to missingness)
## Multiple R-squared:  0.0643, Adjusted R-squared:  0.04713
```

```
## F-statistic: 3.745 on 2 and 109 DF,  p-value: 0.02672
```

**S3.2c: Is the difference in A-cue neural activity a mediator (or moderator) of the difference A-cue bias between Baseline and Proactive sessions?**

**Final Parcels:** Only LH SomMot 22 and LH SomMot 26 had significant relationships between A-cue Bias and A-cue activation in both the Baseline and Proactive session as well difference scores (Proactive - Baseline) analyzed separately A-cue activity can be said to mediate (H.diff) or moderate (H.sum.centered) differences in A-cue bias if the neural activity difference between Proactive and Baseline predictive of behavioral measure difference (i.e. Y.diff ∼H.sum + H.diff); Y=behavioral measure i.e. A-cue bias; Y.diff = Proactive-Baseline behavioral A-cue bias; H=neural measure i.e. knot 4 A-cue activity; H.diff = Proacitve-Baseline neural A-cue activity

Table 3: Parcels that Pass

| ROI | Estimate | std.err | p-value |
|---|---|---|---|
| LHSomMot22H.sum.centered | 0.134 | 0.132 | .312 |
| LHSomMot22H.diff | 0.425 | 0.204 | .04 |
| LHSomMot26H.sum.centered | 0.216 | 0.196 | .271 |
| LHSomMot26H.diff | 0.730 | 0.304 | 0.0179444134964579 |

**S3.2d: Visualization of the within person mediation for each parcel:**

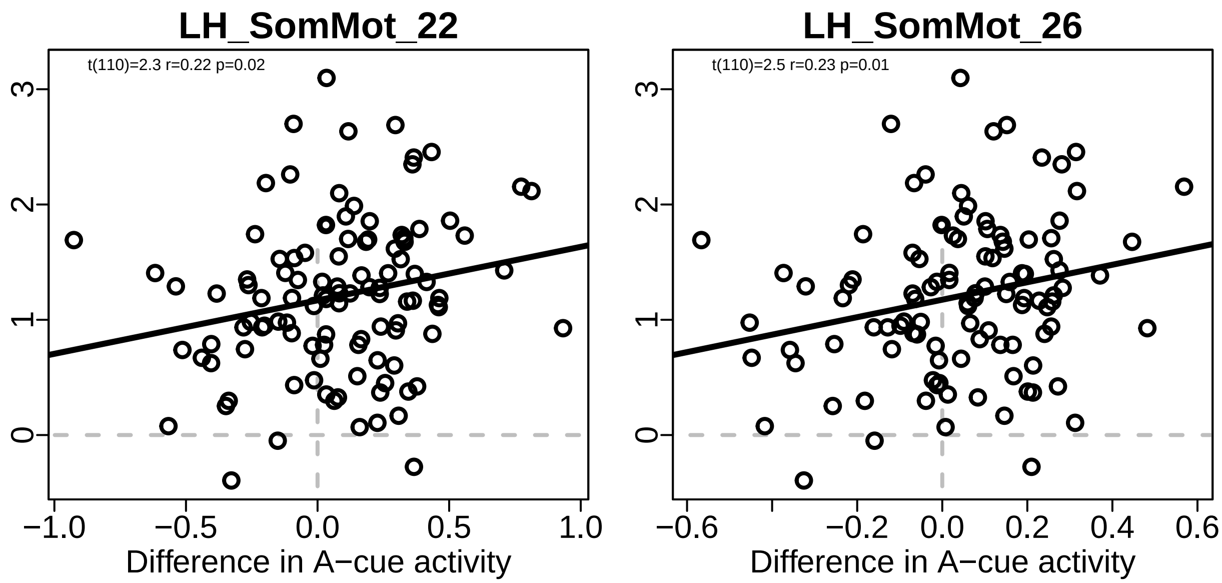


#### S3.3 Combining Contiguous lMOT regions:

Because the two parcels were contiguous, for further analyses we combined them into a single "mega-parcel". Subsequent analyses tables and figures seen in the main text as Table 3 and Figure 4a.
To create the parcellation, we made a new mask containing only voxels from LH SomMot 22 and LH SomMot 26 before using AFNI to create GLMs for all the regressors and contrasts. Information about our GLMs used, see Etzel et. al 2022. For the megaparcel mask, see folder S3.within on OSF.

```
# anatomy for underlay
# p.img <- readNifti(paste0(in.path,"mainTableFigures/nifti/Schaefer2018_400Parcels_7Networks_order_DMCC2p4.nii"));
```

```
# afni.path <- paste0(in.path, "S3.within/afni.exe") # path to afni
# temp.img <- array(0, dim(p.img));   # blank "brain" of zeros
# temp.img[which(p.img == 53)] <- 1;
# temp.img[which(p.img == 57)] <- 1;

# writeNifti(temp.img, paste0("in.path/S3.within/megaparcel.nii.gz"),  # store the new mask to run afni on
#            paste0(in.path, "mainTableFigures/nifti/Schaefer2018_400x7_81x96x81.nii.gz")
# megaparcel_mask.fname <- paste0(in.path, "S3.within/megaparcel.nii.gz");
# for (rid in 1:length(run.ids)) {     # sid <- 1; rid <- 1;
#   stats.fname <- paste0(in.path, "S3.within/sample_sub/sub-example_task-AXCPT_run-", rid, ".nii.gz");
#   txt.fname <- paste0(out.path, "prep_vol/sub-example_ses-wave1bas_task-AXCPT_run", rid, "_vol.txt");
#
#   if (file.exists(np.fname) & !file.exists(txt.fname)) {
#     system2(paste0(afni.path, "3dROIstats"),
#             args=paste0("-mask ", mask.fname, " ", stats.fname, " > ", txt.fname), stdout=TRUE);
#   }
# }
```

##### S3.3a lMOT "mega-parcel" within mediation visualization:

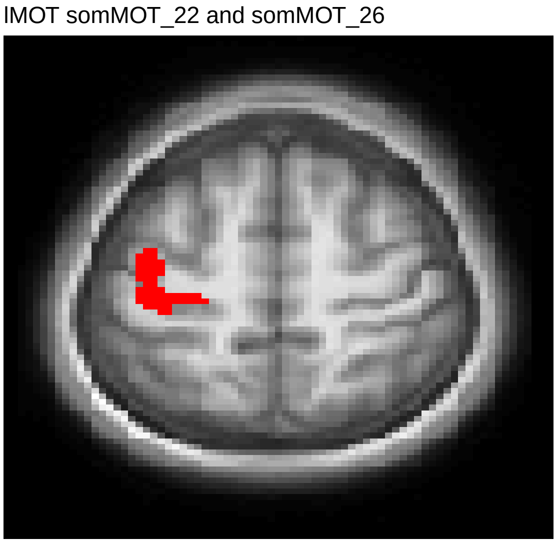


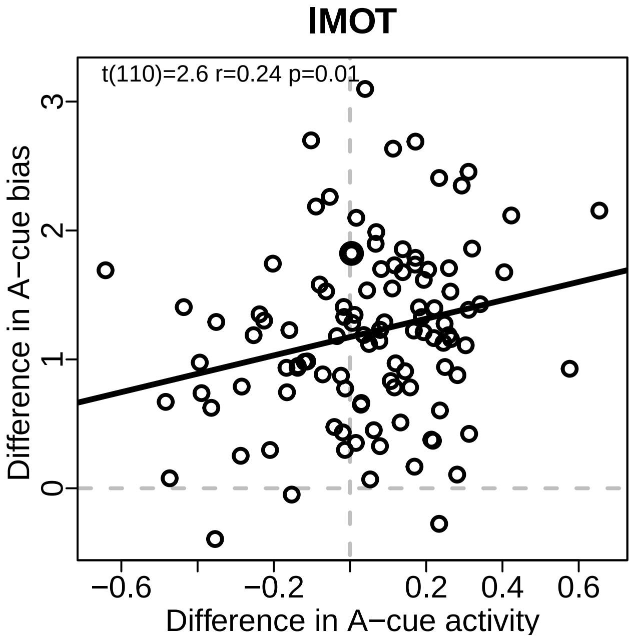


##### S3.3b Mediation output:

```
##
## Call:
## lm(formula = Y.diff ~ H.sum.centered + H.diff)
##
## Residuals:
##      Min       1Q   Median       3Q      Max
## -1.60203 -0.39293 -0.04934  0.40864  1.74207
##
## Coefficients:
##                Estimate Std. Error t value Pr(>|t|)
## (Intercept)      1.1752     0.0640  18.364   <2e-16 ***
## H.sum.centered   0.2321     0.1881   1.234   0.2200
## H.diff           0.6752     0.2799   2.413   0.0175 *
```

```
## ---
## Signif. codes:  0 '***' 0.001 '**' 0.01 '*' 0.05 '.' 0.1 ' ' 1
##
## Residual standard error: 0.661 on 109 degrees of freedom
##   (6 observations deleted due to missingness)
## Multiple R-squared:  0.06904, Adjusted R-squared:  0.05195
## F-statistic: 4.041 on 2 and 109 DF,  p-value: 0.02027
```

| | Estimate | SD | T-value | p-value |
|---|---|---|---|---|
| (Intercept) | 1.175 | 0.064 | 18.364 | <.001 |
| H.sum.centered | .232 | 0.188 | 1.234 | .22 |
| H.diff | .675 | 0.280 | 2.413 | .018 |

### Supplemental 4: Between-Person Path Analyses

##### S4.1 Zero-order and Partial Correlations with Variables of Interest: WMC, rDLPFC, lMOT, A-cue Bias

Proactive session is the only one with significant Zero-order values. Once partialed, the relationships all hold except for the direct relationship with A-cue Bias and WMC.

Table 4: Supplemental Table 4. Correlations between variables of interest for the Baseline and Proactive sessions; Zero-Order and partial correlations.

| Variables | Zero-Order (p-value) | Partial (p-value) |
|---|---|---|
| **Proactive** | | |
| WMC & A-cue Bias | .031 (.737) | .032 (.735) |
| WMC & rDLPFC Activation | -.045 (.631) | -.029 (.765) |
| rDLPFC & lMOT Activation | .401 (.001)* | .363 (.001)* |
| lMOT & A-cue Bias | .275 (.003)* | .271 (.004)* |
| **Baseline** | | |
| WMC & A-cue Bias | -.031 (.741) | -.04 (.67) |
| WMC & rDLPFC Activation | -.028 (.769) | -.026 (.786) |
| rDLPFC & lMOT Activation | .388 (.001)* | .372 (.001)* |
| lMOT & A-cue Bias | .205 (.028)* | .188 (.047)* |

Visualization of Zero-Order and Partial Order Correlations

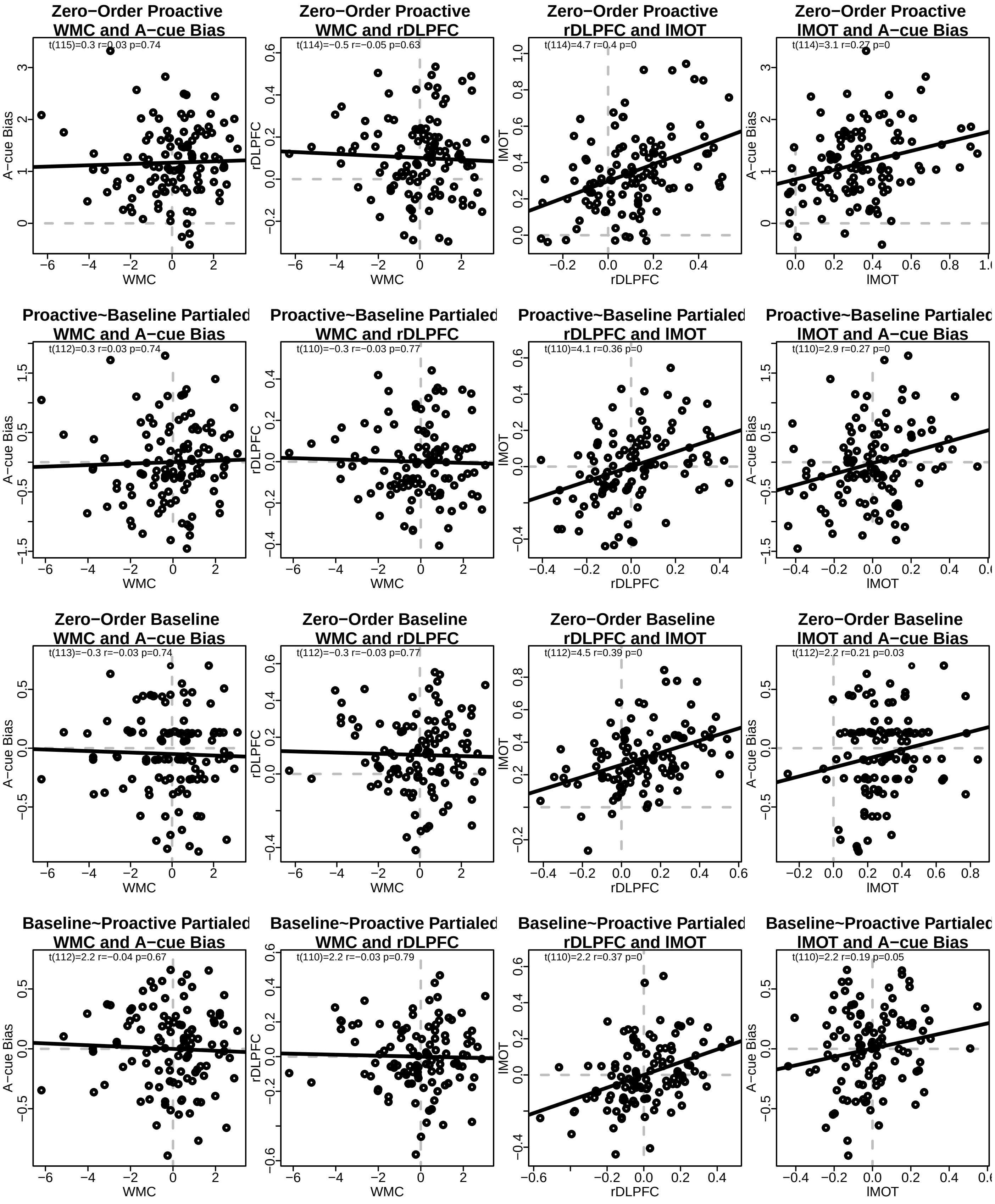

**S4.2 Brain-Behavior Correlations Extreme Groups:**

Timecourses using the 30 participants with the highest and lowest A-cue bias alongside the lMOT parcel used for later within-person mediation and path analyses to show how the effect is driven by this individual difference.
Note, these timecourses are figure 4 and 5 in the main manuscript.

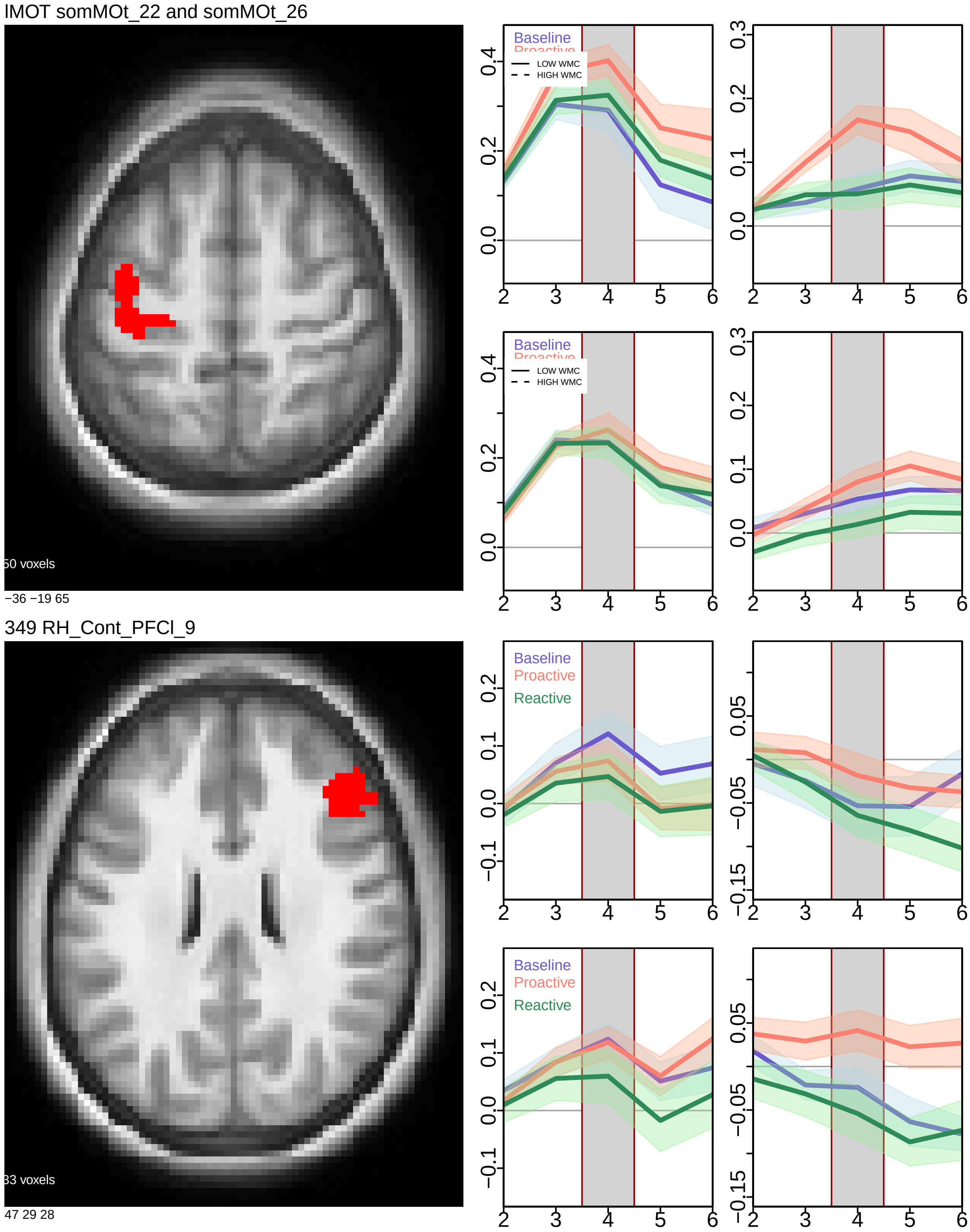

##### S4.3 Path Analysis Between Mediation: Zero-Order Values

Indirect results using proactive session values after partialling out shared variance from the baseline session. This model tests for an additive relationship between WMC-rDLPFC-lMOT-Acue bias. Rows with RED text denote a confidence interval that is all positive or negative; LHS: left hand side; RHS: right hand side; OP: operator; regression; (residual)(co)variance; 1 intercept; NOTE WMC is a static measure mediation path testing for (a) individuals with higher WMC have higher rDLPFC activity, (b) higher rDLPFC activity results in increased LMOT activation during A-cue presentation, and (c) which results in an increased likelihood to make a target response, and further, whether this pathway explains significant variance in (d) the direct path of WMC predicting A-cue bias

| lhs | op | rhs | label | est | se | z | pvalue | ci.lower | ci.upper |
|---|---|---|---|---|---|---|---|---|---|
| pfc9 | | wmc | a | -0.005 | 0.008 | -0.552 | 0.581 | -0.021 | 0.013 |
| lMOT | | pfc9 | b | 0.467 | 0.100 | 4.660 | 0.000 | 0.268 | 0.666 |
| bias | | lMOT | c | 0.917 | 0.252 | 3.646 | 0.000 | 0.427 | 1.421 |
| bias | | wmc | d | 0.016 | 0.039 | 0.423 | 0.672 | -0.056 | 0.097 |
| pfc9 | | pfc9 | | 0.033 | 0.004 | 7.820 | 0.000 | 0.024 | 0.040 |
| lMOT | | lMOT | | 0.037 | 0.005 | 7.171 | 0.000 | 0.027 | 0.047 |
| bias | | bias | | 0.450 | 0.061 | 7.376 | 0.000 | 0.326 | 0.563 |
| wmc | | wmc | | 3.117 | 0.000 | NA | NA | 3.117 | 3.117 |
| pfc9 | 1 | | | 0.101 | 0.017 | 5.963 | 0.000 | 0.068 | 0.136 |
| lMOT | 1 | | | 0.297 | 0.019 | 15.830 | 0.000 | 0.260 | 0.334 |
| bias | 1 | | | 0.845 | 0.103 | 8.212 | 0.000 | 0.648 | 1.050 |
| wmc | 1 | | | 0.006 | 0.000 | NA | NA | 0.006 | 0.006 |
| abc | := | (a*b*c) | abc | -0.002 | 0.004 | -0.525 | 0.600 | -0.009 | 0.006 |
| total | := | d+(a*b*c) | total | 0.014 | 0.038 | 0.380 | 0.704 | -0.056 | 0.094 |

```
##      lhs op       rhs label    est    se      z pvalue ci.lower ci.upper
## 1   pfc9  ~       wmc     a -0.005 0.008 -0.552  0.581   -0.021    0.013
## 2   lMOT  ~      pfc9     b  0.467 0.100  4.660  0.000    0.268    0.666
## 3   bias  ~      lMOT     c  0.917 0.252  3.646  0.000    0.427    1.421
## 4   bias  ~       wmc     d  0.016 0.039  0.423  0.672   -0.056    0.097
## 5   pfc9 ~~      pfc9        0.033 0.004  7.820  0.000    0.024    0.040
## 6   lMOT ~~      lMOT        0.037 0.005  7.171  0.000    0.027    0.047
## 7   bias ~~      bias        0.450 0.061  7.376  0.000    0.326    0.563
## 8    wmc ~~       wmc        3.117 0.000     NA     NA    3.117    3.117
## 9   pfc9 ~1                  0.101 0.017  5.963  0.000    0.068    0.136
## 10  lMOT ~1                  0.297 0.019 15.830  0.000    0.260    0.334
## 11  bias ~1                  0.845 0.103  8.212  0.000    0.648    1.050
## 12   wmc ~1                  0.006 0.000     NA     NA    0.006    0.006
## 13   abc :=   (a*b*c)   abc -0.002 0.004 -0.525  0.600   -0.009    0.006
## 14 total := d+(a*b*c) total  0.014 0.038  0.380  0.704   -0.056    0.094
## lavaan 0.6-18 ended normally after 1 iteration
##
##   Estimator                                         ML
##   Optimization method                           NLMINB
##   Number of model parameters                        10
##
##                                                   Used       Total
##   Number of observations                           116         118
##
## Model Test User Model:
##
##   Test statistic                                 3.907
##   Degrees of freedom                                 2
##   P-value (Chi-square)                           0.142
##
## Model Test Baseline Model:
##
##   Test statistic                                33.842
```

```
##   Degrees of freedom                                 6
##   P-value                                        0.000
##
## User Model versus Baseline Model:
##
##   Comparative Fit Index (CFI)                    0.932
##   Tucker-Lewis Index (TLI)                       0.795
##
## Loglikelihood and Information Criteria:
##
##   Loglikelihood user model (H0)                -57.469
##   Loglikelihood unrestricted model (H1)        -55.516
##
##   Akaike (AIC)                                 134.938
##   Bayesian (BIC)                               162.474
##   Sample-size adjusted Bayesian (SABIC)        130.864
##
## Root Mean Square Error of Approximation:
##
##   RMSEA                                          0.091
##   90 Percent confidence interval - lower         0.000
##   90 Percent confidence interval - upper         0.225
##   P-value H_0: RMSEA <= 0.050                    0.222
##   P-value H_0: RMSEA >= 0.080                    0.659
##
## Standardized Root Mean Square Residual:
##
##   SRMR                                           0.043
##
## Parameter Estimates:
##
##   Standard errors                            Bootstrap
##   Number of requested bootstrap draws             5000
##   Number of successful bootstrap draws            5000
##
## Regressions:
##                    Estimate  Std.Err  z-value  P(>|z|) ci.lower ci.upper   Std.lv  Std.all
##   pfc9 ~
##     wmc        (a)   -0.005    0.008   -0.552    0.581   -0.021    0.013   -0.005   -0.045
##   lMOT ~
##     pfc9       (b)    0.467    0.100    4.660    0.000    0.268    0.666    0.467    0.401
##   bias ~
##     lMOT       (c)    0.917    0.252    3.646    0.000    0.427    1.421    0.917    0.276
##     wmc        (d)    0.016    0.039    0.423    0.672   -0.056    0.097    0.016    0.042
##
## Intercepts:
##                    Estimate  Std.Err  z-value  P(>|z|) ci.lower ci.upper   Std.lv  Std.all
##    .pfc9              0.101    0.017    5.963    0.000    0.068    0.136    0.101    0.560
##    .lMOT              0.297    0.019   15.830    0.000    0.260    0.334    0.297    1.416
##    .bias              0.845    0.103    8.212    0.000    0.648    1.050    0.845    1.210
##
## Variances:
##                    Estimate  Std.Err  z-value  P(>|z|) ci.lower ci.upper   Std.lv  Std.all
##    .pfc9              0.033    0.004    7.820    0.000    0.024    0.040    0.033    0.998
##    .lMOT              0.037    0.005    7.171    0.000    0.027    0.047    0.037    0.839
##    .bias              0.450    0.061    7.376    0.000    0.326    0.563    0.450    0.923
##
## R-Square:
##                    Estimate
##     pfc9              0.002
##     lMOT              0.161
```

```
##     bias              0.077
##
## Defined Parameters:
##                    Estimate  Std.Err  z-value  P(>|z|) ci.lower ci.upper   Std.lv  Std.all
##     abc              -0.002    0.004   -0.525    0.600   -0.009    0.006   -0.002   -0.005
##     total             0.014    0.038    0.380    0.704   -0.056    0.094    0.014    0.037
```

##### S4.4 Path Analysis Between Mediation: Proactive-Baseline Subtraction Values

Next, we tested using Proactive-Baseline subtraction method for further sensitivity:

| lhs | op | rhs | label | est | se | z | pvalue | ci.lower | ci.upper |
|---|---|---|---|---|---|---|---|---|---|
| pfc9 | | wmc | a | 0.000 | 0.011 | 0.005 | 0.996 | -0.021 | 0.022 |
| lMOT | | pfc9 | b | 0.356 | 0.087 | 4.086 | 0.000 | 0.188 | 0.529 |
| bias | | lMOT | c | 0.720 | 0.273 | 2.635 | 0.008 | 0.174 | 1.246 |
| bias | | wmc | d | 0.020 | 0.037 | 0.541 | 0.588 | -0.047 | 0.098 |
| pfc9 | | pfc9 | | 0.050 | 0.007 | 6.821 | 0.000 | 0.036 | 0.065 |
| lMOT | | lMOT | | 0.044 | 0.007 | 6.236 | 0.000 | 0.031 | 0.059 |
| bias | | bias | | 0.430 | 0.058 | 7.374 | 0.000 | 0.312 | 0.542 |
| wmc | | wmc | | 3.022 | 0.000 | NA | NA | 3.022 | 3.022 |
| pfc9 | 1 | | | -0.005 | 0.021 | -0.223 | 0.824 | -0.046 | 0.038 |
| lMOT | 1 | | | 0.052 | 0.020 | 2.581 | 0.010 | 0.011 | 0.091 |
| bias | 1 | | | 1.174 | 0.062 | 18.825 | 0.000 | 1.054 | 1.296 |
| wmc | 1 | | | -0.065 | 0.000 | NA | NA | -0.065 | -0.065 |
| abc | := | (a*b*c) | abc | 0.000 | 0.003 | 0.005 | 0.996 | -0.007 | 0.007 |
| total | := | d+(a*b*c) | total | 0.020 | 0.037 | 0.538 | 0.590 | -0.047 | 0.098 |
| prop | := | a*b*c/total | prop | 0.001 | 25.508 | 0.000 | 1.000 | -0.836 | 0.810 |

```
## lavaan 0.6-18 ended normally after 7 iterations
##
##   Estimator                                         ML
##   Optimization method                           NLMINB
##   Number of model parameters                        10
##
##                                                   Used       Total
##   Number of observations                           112         118
##
## Model Test User Model:
##
##   Test statistic                                 0.900
##   Degrees of freedom                                 2
##   P-value (Chi-square)                           0.638
##
## Model Test Baseline Model:
##
##   Test statistic                                22.792
##   Degrees of freedom                                 6
##   P-value                                        0.001
##
## User Model versus Baseline Model:
##
##   Comparative Fit Index (CFI)                    1.000
##   Tucker-Lewis Index (TLI)                       1.197
##
## Loglikelihood and Information Criteria:
##
##   Loglikelihood user model (H0)                -87.233
##   Loglikelihood unrestricted model (H1)        -86.783
```

```
##
##   Akaike (AIC)                                 194.466
##   Bayesian (BIC)                               221.651
##   Sample-size adjusted Bayesian (SABIC)        190.047
##
## Root Mean Square Error of Approximation:
##
##   RMSEA                                          0.000
##   90 Percent confidence interval - lower         0.000
##   90 Percent confidence interval - upper         0.148
##   P-value H_0: RMSEA <= 0.050                    0.710
##   P-value H_0: RMSEA >= 0.080                    0.205
##
## Standardized Root Mean Square Residual:
##
##   SRMR                                           0.022
##
## Parameter Estimates:
##
##   Standard errors                            Bootstrap
##   Number of requested bootstrap draws             5000
##   Number of successful bootstrap draws            5000
##
## Regressions:
##                    Estimate  Std.Err  z-value  P(>|z|) ci.lower ci.upper   Std.lv  Std.all
##   pfc9 ~
##     wmc        (a)    0.000    0.011    0.005    0.996   -0.021    0.022    0.000    0.000
##   lMOT ~
##     pfc9       (b)    0.356    0.087    4.086    0.000    0.188    0.529    0.356    0.355
##   bias ~
##     lMOT       (c)    0.720    0.273    2.635    0.008    0.174    1.246    0.720    0.239
##     wmc        (d)    0.020    0.037    0.541    0.588   -0.047    0.098    0.020    0.051
##
## Intercepts:
##                    Estimate  Std.Err  z-value  P(>|z|) ci.lower ci.upper   Std.lv  Std.all
##    .pfc9             -0.005    0.021   -0.223    0.824   -0.046    0.038   -0.005   -0.021
##    .lMOT              0.052    0.020    2.581    0.010    0.011    0.091    0.052    0.230
##    .bias              1.174    0.062   18.825    0.000    1.054    1.296    1.174    1.737
##
## Variances:
##                    Estimate  Std.Err  z-value  P(>|z|) ci.lower ci.upper   Std.lv  Std.all
##    .pfc9              0.050    0.007    6.821    0.000    0.036    0.065    0.050    1.000
##    .lMOT              0.044    0.007    6.236    0.000    0.031    0.059    0.044    0.874
##    .bias              0.430    0.058    7.374    0.000    0.312    0.542    0.430    0.940
##
## R-Square:
##                    Estimate
##     pfc9              0.000
##     lMOT              0.126
##     bias              0.060
##
## Defined Parameters:
##                    Estimate  Std.Err  z-value  P(>|z|) ci.lower ci.upper   Std.lv  Std.all
##     abc               0.000    0.003    0.005    0.996   -0.007    0.007    0.000    0.000
##     total             0.020    0.037    0.538    0.590   -0.047    0.098    0.020    0.051
##     prop              0.001   25.508    0.000    1.000   -0.836    0.810    0.001    0.001
##
##      lhs op         rhs label    est     se      z pvalue ci.lower ci.upper
## 1   pfc9  ~         wmc     a  0.000  0.011  0.005  0.996   -0.021    0.022
## 2   lMOT  ~        pfc9     b  0.356  0.087  4.086  0.000    0.188    0.529
## 3   bias  ~        lMOT     c  0.720  0.273  2.635  0.008    0.174    1.246
```

```
## 4   bias  ~         wmc     d  0.020  0.037  0.541  0.588   -0.047    0.098
## 5   pfc9 ~~        pfc9        0.050  0.007  6.821  0.000    0.036    0.065
## 6   lMOT ~~        lMOT        0.044  0.007  6.236  0.000    0.031    0.059
## 7   bias ~~        bias        0.430  0.058  7.374  0.000    0.312    0.542
## 8    wmc ~~         wmc        3.022  0.000     NA     NA    3.022    3.022
## 9   pfc9 ~1                   -0.005  0.021 -0.223  0.824   -0.046    0.038
## 10  lMOT ~1                    0.052  0.020  2.581  0.010    0.011    0.091
## 11  bias ~1                    1.174  0.062 18.825  0.000    1.054    1.296
## 12   wmc ~1                   -0.065  0.000     NA     NA   -0.065   -0.065
## 13   abc :=     (a*b*c)   abc  0.000  0.003  0.005  0.996   -0.007    0.007
## 14 total :=   d+(a*b*c) total  0.020  0.037  0.538  0.590   -0.047    0.098
## 15  prop := a*b*c/total  prop  0.001 25.508  0.000  1.000   -0.836    0.810
```

##### S4.5 Path Analysis Between Mediation: Proactive with Baseline values partialled out

Next, we tested using Proactive data with Baseline values partialed out for further sensitivity: Note this is Table 4 in the main manuscript.

| lhs | op | rhs | label | est | se | z | pvalue | ci.lower | ci.upper |
|---|---|---|---|---|---|---|---|---|---|
| pfc9 | | wmc | a | -0.003 | 0.008 | -0.355 | 0.723 | -0.018 | 0.013 |
| lMOT | | pfc9 | b | 0.406 | 0.095 | 4.256 | 0.000 | 0.221 | 0.597 |
| bias | | lMOT | c | 0.925 | 0.300 | 3.085 | 0.002 | 0.328 | 1.509 |
| bias | | wmc | d | 0.016 | 0.037 | 0.443 | 0.657 | -0.052 | 0.093 |
| pfc9 | | pfc9 | | 0.030 | 0.004 | 7.710 | 0.000 | 0.022 | 0.038 |
| lMOT | | lMOT | | 0.033 | 0.005 | 7.168 | 0.000 | 0.024 | 0.042 |
| bias | | bias | | 0.408 | 0.054 | 7.594 | 0.000 | 0.301 | 0.511 |
| wmc | | wmc | | 3.022 | 0.000 | NA | NA | 3.022 | 3.022 |
| pfc9 | 1 | | | 0.000 | 0.017 | -0.011 | 0.991 | -0.033 | 0.032 |
| lMOT | 1 | | | 0.000 | 0.017 | 0.000 | 1.000 | -0.033 | 0.034 |
| bias | 1 | | | -0.014 | 0.061 | -0.235 | 0.814 | -0.132 | 0.106 |
| wmc | 1 | | | -0.065 | 0.000 | NA | NA | -0.065 | -0.065 |
| abc | := | (a*b*c) | abc | -0.001 | 0.003 | -0.335 | 0.738 | -0.008 | 0.006 |
| total | := | d+abc | total | 0.015 | 0.037 | 0.419 | 0.675 | -0.052 | 0.092 |

```
## lavaan 0.6-18 ended normally after 12 iterations
##
##   Estimator                                         ML
##   Optimization method                           NLMINB
##   Number of model parameters                        10
##
##                                                   Used       Total
##   Number of observations                           112         118
##
## Model Test User Model:
##
##   Test statistic                                 2.162
##   Degrees of freedom                                 2
##   P-value (Chi-square)                           0.339
##
## Model Test Baseline Model:
##
##   Test statistic                                26.793
##   Degrees of freedom                                 6
##   P-value                                        0.000
##
## User Model versus Baseline Model:
##
```

```
##   Comparative Fit Index (CFI)                    0.992
##   Tucker-Lewis Index (TLI)                       0.977
##
## Loglikelihood and Information Criteria:
##
##   Loglikelihood user model (H0)                -40.031
##   Loglikelihood unrestricted model (H1)        -38.950
##
##   Akaike (AIC)                                 100.062
##   Bayesian (BIC)                               127.247
##   Sample-size adjusted Bayesian (SABIC)         95.643
##
## Root Mean Square Error of Approximation:
##
##   RMSEA                                          0.027
##   90 Percent confidence interval - lower         0.000
##   90 Percent confidence interval - upper         0.191
##   P-value H_0: RMSEA <= 0.050                    0.435
##   P-value H_0: RMSEA >= 0.080                    0.438
##
## Standardized Root Mean Square Residual:
##
##   SRMR                                           0.033
##
## Parameter Estimates:
##
##   Standard errors                            Bootstrap
##   Number of requested bootstrap draws             5000
##   Number of successful bootstrap draws            5000
##
## Regressions:
##                    Estimate  Std.Err  z-value  P(>|z|) ci.lower ci.upper   Std.lv  Std.all
##   pfc9 ~
##     wmc        (a)   -0.003    0.008   -0.355    0.723   -0.018    0.013   -0.003   -0.029
##   lMOT ~
##     pfc9       (b)    0.406    0.095    4.256    0.000    0.221    0.597    0.406    0.363
##   bias ~
##     lMOT       (c)    0.925    0.300    3.085    0.002    0.328    1.509    0.925    0.272
##     wmc        (d)    0.016    0.037    0.443    0.657   -0.052    0.093    0.016    0.043
##
## Intercepts:
##                    Estimate  Std.Err  z-value  P(>|z|) ci.lower ci.upper   Std.lv  Std.all
##    .pfc9             -0.000    0.017   -0.011    0.991   -0.033    0.032   -0.000   -0.001
##    .lMOT              0.000    0.017    0.000    1.000   -0.033    0.034    0.000    0.000
##    .bias             -0.014    0.061   -0.235    0.814   -0.132    0.106   -0.014   -0.022
##
## Variances:
##                    Estimate  Std.Err  z-value  P(>|z|) ci.lower ci.upper   Std.lv  Std.all
##    .pfc9              0.030    0.004    7.710    0.000    0.022    0.038    0.030    0.999
##    .lMOT              0.033    0.005    7.168    0.000    0.024    0.042    0.033    0.868
##    .bias              0.408    0.054    7.594    0.000    0.301    0.511    0.408    0.924
##
## R-Square:
##                    Estimate
##     pfc9              0.001
##     lMOT              0.132
##     bias              0.076
##
## Defined Parameters:
##                    Estimate  Std.Err  z-value  P(>|z|) ci.lower ci.upper   Std.lv  Std.all
##     abc              -0.001    0.003   -0.335    0.738   -0.008    0.006   -0.001   -0.003
```

```
##     total             0.015    0.037    0.419    0.675   -0.052    0.092    0.015    0.040
##
##      lhs op     rhs label    est    se      z pvalue ci.lower ci.upper
## 1   pfc9  ~     wmc     a -0.003 0.008 -0.355  0.723   -0.018    0.013
## 2   lMOT  ~    pfc9     b  0.406 0.095  4.256  0.000    0.221    0.597
## 3   bias  ~    lMOT     c  0.925 0.300  3.085  0.002    0.328    1.509
## 4   bias  ~     wmc     d  0.016 0.037  0.443  0.657   -0.052    0.093
## 5   pfc9 ~~    pfc9        0.030 0.004  7.710  0.000    0.022    0.038
## 6   lMOT ~~    lMOT        0.033 0.005  7.168  0.000    0.024    0.042
## 7   bias ~~    bias        0.408 0.054  7.594  0.000    0.301    0.511
## 8    wmc ~~     wmc        3.022 0.000     NA     NA    3.022    3.022
## 9   pfc9 ~1                0.000 0.017 -0.011  0.991   -0.033    0.032
## 10  lMOT ~1                0.000 0.017  0.000  1.000   -0.033    0.034
## 11  bias ~1               -0.014 0.061 -0.235  0.814   -0.132    0.106
## 12   wmc ~1               -0.065 0.000     NA     NA   -0.065   -0.065
## 13   abc := (a*b*c)   abc -0.001 0.003 -0.335  0.738   -0.008    0.006
## 14 total :=   d+abc total  0.015 0.037  0.419  0.675   -0.052    0.092
```

Visualization of between path analysis using residualized proactive session values with significant beta estimates listed. In this model, the direct path was no longer significant, but the indirect (abc) and total d + (abc) was significant.

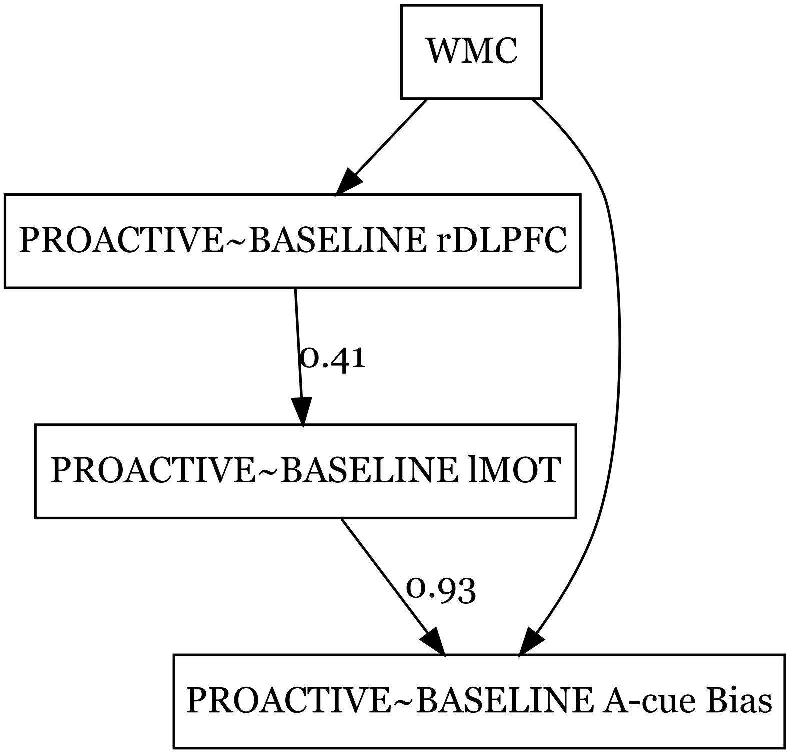


##### S4.6 Control Analysis Between Mediation: Baseline

For additional control, we tested the indirect path using Baseline session values to ensure the path is selective to the proactive session.

```
## lavaan 0.6-18 ended normally after 12 iterations
##
##   Estimator                                         ML
##   Optimization method                           NLMINB
##   Number of model parameters                        10
##
```

| lhs | op | rhs | label | est | se | z | pvalue | ci.lower | ci.upper |
|---|---|---|---|---|---|---|---|---|---|
| pfc9 | | wmc | a | -0.003 | 0.008 | -0.355 | 0.723 | -0.018 | 0.013 |
| lMOT | | pfc9 | b | 0.406 | 0.095 | 4.256 | 0.000 | 0.221 | 0.597 |
| bias | | lMOT | c | 0.925 | 0.300 | 3.085 | 0.002 | 0.328 | 1.509 |
| bias | | wmc | d | 0.016 | 0.037 | 0.443 | 0.657 | -0.052 | 0.093 |
| pfc9 | | pfc9 | | 0.030 | 0.004 | 7.710 | 0.000 | 0.022 | 0.038 |
| lMOT | | lMOT | | 0.033 | 0.005 | 7.168 | 0.000 | 0.024 | 0.042 |
| bias | | bias | | 0.408 | 0.054 | 7.594 | 0.000 | 0.301 | 0.511 |
| wmc | | wmc | | 3.022 | 0.000 | NA | NA | 3.022 | 3.022 |
| pfc9 | 1 | | | 0.000 | 0.017 | -0.011 | 0.991 | -0.033 | 0.032 |
| lMOT | 1 | | | 0.000 | 0.017 | 0.000 | 1.000 | -0.033 | 0.034 |
| bias | 1 | | | -0.014 | 0.061 | -0.235 | 0.814 | -0.132 | 0.106 |
| wmc | 1 | | | -0.065 | 0.000 | NA | NA | -0.065 | -0.065 |
| abc | := | (a*b*c) | abc | -0.001 | 0.003 | -0.335 | 0.738 | -0.008 | 0.006 |
| total | := | d+abc | total | 0.015 | 0.037 | 0.419 | 0.675 | -0.052 | 0.092 |

```
##                                                   Used       Total
##   Number of observations                           112         118
##
## Model Test User Model:
##
##   Test statistic                                 2.162
##   Degrees of freedom                                 2
##   P-value (Chi-square)                           0.339
##
## Model Test Baseline Model:
##
##   Test statistic                                26.793
##   Degrees of freedom                                 6
##   P-value                                        0.000
##
## User Model versus Baseline Model:
##
##   Comparative Fit Index (CFI)                    0.992
##   Tucker-Lewis Index (TLI)                       0.977
##
## Loglikelihood and Information Criteria:
##
##   Loglikelihood user model (H0)                -40.031
##   Loglikelihood unrestricted model (H1)        -38.950
##
##   Akaike (AIC)                                 100.062
##   Bayesian (BIC)                               127.247
##   Sample-size adjusted Bayesian (SABIC)         95.643
##
## Root Mean Square Error of Approximation:
##
##   RMSEA                                          0.027
##   90 Percent confidence interval - lower         0.000
##   90 Percent confidence interval - upper         0.191
##   P-value H_0: RMSEA <= 0.050                    0.435
##   P-value H_0: RMSEA >= 0.080                    0.438
##
## Standardized Root Mean Square Residual:
##
##   SRMR                                           0.033
##
## Parameter Estimates:
##
```

```
##   Standard errors                             Bootstrap
##   Number of requested bootstrap draws              5000
##   Number of successful bootstrap draws             5000
##
## Regressions:
##                    Estimate  Std.Err  z-value  P(>|z|) ci.lower ci.upper   Std.lv  Std.all
##   pfc9 ~
##     wmc        (a)   -0.003    0.008   -0.355    0.723   -0.018    0.013   -0.003   -0.029
##   lMOT ~
##     pfc9       (b)    0.406    0.095    4.256    0.000    0.221    0.597    0.406    0.363
##   bias ~
##     lMOT       (c)    0.925    0.300    3.085    0.002    0.328    1.509    0.925    0.272
##     wmc        (d)    0.016    0.037    0.443    0.657   -0.052    0.093    0.016    0.043
##
## Intercepts:
##                    Estimate  Std.Err  z-value  P(>|z|) ci.lower ci.upper   Std.lv  Std.all
##    .pfc9             -0.000    0.017   -0.011    0.991   -0.033    0.032   -0.000   -0.001
##    .lMOT              0.000    0.017    0.000    1.000   -0.033    0.034    0.000    0.000
##    .bias             -0.014    0.061   -0.235    0.814   -0.132    0.106   -0.014   -0.022
##
## Variances:
##                    Estimate  Std.Err  z-value  P(>|z|) ci.lower ci.upper   Std.lv  Std.all
##    .pfc9              0.030    0.004    7.710    0.000    0.022    0.038    0.030    0.999
##    .lMOT              0.033    0.005    7.168    0.000    0.024    0.042    0.033    0.868
##    .bias              0.408    0.054    7.594    0.000    0.301    0.511    0.408    0.924
##
## R-Square:
##                    Estimate
##     pfc9              0.001
##     lMOT              0.132
##     bias              0.076
##
## Defined Parameters:
##                    Estimate  Std.Err  z-value  P(>|z|) ci.lower ci.upper   Std.lv  Std.all
##     abc              -0.001    0.003   -0.335    0.738   -0.008    0.006   -0.001   -0.003
##     total             0.015    0.037    0.419    0.675   -0.052    0.092    0.015    0.040
##
##      lhs op     rhs label    est    se      z pvalue ci.lower ci.upper
## 1   pfc9  ~     wmc     a -0.003 0.008 -0.355  0.723   -0.018    0.013
## 2   lMOT  ~    pfc9     b  0.406 0.095  4.256  0.000    0.221    0.597
## 3   bias  ~    lMOT     c  0.925 0.300  3.085  0.002    0.328    1.509
## 4   bias  ~     wmc     d  0.016 0.037  0.443  0.657   -0.052    0.093
## 5   pfc9 ~~    pfc9        0.030 0.004  7.710  0.000    0.022    0.038
## 6   lMOT ~~    lMOT        0.033 0.005  7.168  0.000    0.024    0.042
## 7   bias ~~    bias        0.408 0.054  7.594  0.000    0.301    0.511
## 8    wmc ~~     wmc        3.022 0.000     NA     NA    3.022    3.022
## 9   pfc9 ~1                0.000 0.017 -0.011  0.991   -0.033    0.032
## 10  lMOT ~1                0.000 0.017  0.000  1.000   -0.033    0.034
## 11  bias ~1               -0.014 0.061 -0.235  0.814   -0.132    0.106
## 12   wmc ~1               -0.065 0.000     NA     NA   -0.065   -0.065
## 13   abc := (a*b*c)   abc -0.001 0.003 -0.335  0.738   -0.008    0.006
## 14 total :=   d+abc total  0.015 0.037  0.419  0.675   -0.052    0.092
```

##### S4.7 Control Analysis Between Mediation: New Order

For control, we first tested the most sensitive indirect path (partialed) in a different order to ensure the path is selective to the rDLPFC lMOT circuit:

| lhs | op | rhs | label | est | se | z | pvalue | ci.lower | ci.upper |
|---|---|---|---|---|---|---|---|---|---|
| lMOT | | wmc | a | -0.004 | 0.010 | -0.385 | 0.701 | -0.023 | 0.018 |
| pfc9 | | lMOT | b | 0.324 | 0.068 | 4.785 | 0.000 | 0.191 | 0.459 |
| bias | | pfc9 | c | -0.088 | 0.344 | -0.256 | 0.798 | -0.750 | 0.579 |
| bias | | wmc | d | 0.012 | 0.039 | 0.320 | 0.749 | -0.059 | 0.093 |
| lMOT | | lMOT | | 0.038 | 0.005 | 7.419 | 0.000 | 0.028 | 0.048 |
| pfc9 | | pfc9 | | 0.026 | 0.004 | 7.045 | 0.000 | 0.019 | 0.034 |
| bias | | bias | | 0.440 | 0.058 | 7.602 | 0.000 | 0.320 | 0.551 |
| wmc | | wmc | | 3.022 | 0.000 | NA | NA | 3.022 | 3.022 |
| lMOT | 1 | | | 0.000 | 0.019 | -0.014 | 0.989 | -0.038 | 0.036 |
| pfc9 | 1 | | | 0.000 | 0.015 | 0.000 | 1.000 | -0.030 | 0.031 |
| bias | 1 | | | -0.015 | 0.063 | -0.235 | 0.814 | -0.140 | 0.104 |
| wmc | 1 | | | -0.065 | 0.000 | NA | NA | -0.065 | -0.065 |
| abc | := | (a*b*c) | abc | 0.000 | 0.001 | 0.090 | 0.928 | -0.002 | 0.003 |
| total | := | d+(a*b*c) | total | 0.013 | 0.039 | 0.323 | 0.746 | -0.058 | 0.094 |

```
## lavaan 0.6-18 ended normally after 1 iteration
##
##   Estimator                                         ML
##   Optimization method                           NLMINB
##   Number of model parameters                        10
##
##                                                   Used       Total
##   Number of observations                           112         118
##
## Model Test User Model:
##
##   Test statistic                                10.664
##   Degrees of freedom                                 2
##   P-value (Chi-square)                           0.005
##
## Model Test Baseline Model:
##
##   Test statistic                                26.793
##   Degrees of freedom                                 6
##   P-value                                        0.000
##
## User Model versus Baseline Model:
##
##   Comparative Fit Index (CFI)                    0.583
##   Tucker-Lewis Index (TLI)                      -0.250
##
## Loglikelihood and Information Criteria:
##
##   Loglikelihood user model (H0)                -44.282
##   Loglikelihood unrestricted model (H1)        -38.950
##
##   Akaike (AIC)                                 108.563
##   Bayesian (BIC)                               135.748
##   Sample-size adjusted Bayesian (SABIC)        104.145
##
## Root Mean Square Error of Approximation:
##
##   RMSEA                                          0.197
##   90 Percent confidence interval - lower         0.092
##   90 Percent confidence interval - upper         0.320
##   P-value H_0: RMSEA <= 0.050                    0.014
##   P-value H_0: RMSEA >= 0.080                    0.965
##
```

```
## Standardized Root Mean Square Residual:
##
##   SRMR                                           0.075
##
## Parameter Estimates:
##
##   Standard errors                            Bootstrap
##   Number of requested bootstrap draws             5000
##   Number of successful bootstrap draws            5000
##
## Regressions:
##                    Estimate  Std.Err  z-value  P(>|z|) ci.lower ci.upper   Std.lv  Std.all
##   lMOT ~
##     wmc        (a)   -0.004    0.010   -0.385    0.701   -0.023    0.018   -0.004   -0.036
##   pfc9 ~
##     lMOT       (b)    0.324    0.068    4.785    0.000    0.191    0.459    0.324    0.363
##   bias ~
##     pfc9       (c)   -0.088    0.344   -0.256    0.798   -0.750    0.579   -0.088   -0.023
##     wmc        (d)    0.012    0.039    0.320    0.749   -0.059    0.093    0.012    0.033
##
## Intercepts:
##                    Estimate  Std.Err  z-value  P(>|z|) ci.lower ci.upper   Std.lv  Std.all
##    .lMOT             -0.000    0.019   -0.014    0.989   -0.038    0.036   -0.000   -0.001
##    .pfc9              0.000    0.015    0.000    1.000   -0.030    0.031    0.000    0.000
##    .bias             -0.015    0.063   -0.235    0.814   -0.140    0.104   -0.015   -0.022
##
## Variances:
##                    Estimate  Std.Err  z-value  P(>|z|) ci.lower ci.upper   Std.lv  Std.all
##    .lMOT              0.038    0.005    7.419    0.000    0.028    0.048    0.038    0.999
##    .pfc9              0.026    0.004    7.045    0.000    0.019    0.034    0.026    0.868
##    .bias              0.440    0.058    7.602    0.000    0.320    0.551    0.440    0.998
##
## R-Square:
##                    Estimate
##     lMOT              0.001
##     pfc9              0.132
##     bias              0.002
##
## Defined Parameters:
##                    Estimate  Std.Err  z-value  P(>|z|) ci.lower ci.upper   Std.lv  Std.all
##     abc               0.000    0.001    0.090    0.928   -0.002    0.003    0.000    0.000
##     total             0.013    0.039    0.323    0.746   -0.058    0.094    0.013    0.033
##
##      lhs op       rhs label    est    se      z pvalue ci.lower ci.upper
## 1   lMOT  ~       wmc     a -0.004 0.010 -0.385  0.701   -0.023    0.018
## 2   pfc9  ~      lMOT     b  0.324 0.068  4.785  0.000    0.191    0.459
## 3   bias  ~      pfc9     c -0.088 0.344 -0.256  0.798   -0.750    0.579
## 4   bias  ~       wmc     d  0.012 0.039  0.320  0.749   -0.059    0.093
## 5   lMOT ~~      lMOT        0.038 0.005  7.419  0.000    0.028    0.048
## 6   pfc9 ~~      pfc9        0.026 0.004  7.045  0.000    0.019    0.034
## 7   bias ~~      bias        0.440 0.058  7.602  0.000    0.320    0.551
## 8    wmc ~~       wmc        3.022 0.000     NA     NA    3.022    3.022
## 9   lMOT ~1                  0.000 0.019 -0.014  0.989   -0.038    0.036
## 10  pfc9 ~1                  0.000 0.015  0.000  1.000   -0.030    0.031
## 11  bias ~1                 -0.015 0.063 -0.235  0.814   -0.140    0.104
## 12   wmc ~1                 -0.065 0.000     NA     NA   -0.065   -0.065
## 13   abc :=   (a*b*c)   abc  0.000 0.001  0.090  0.928   -0.002    0.003
## 14 total := d+(a*b*c) total  0.013 0.039  0.323  0.746   -0.058    0.094
```

### Supplemental 5: Twin-Related Analyses

#### S5.1 Working Memory Capacity

First we checked for correlations amongst the twins for the two span tasks used to make the composite working memory capacity (WMC) score ($r=0.63$ $p<0.001$). Next, we checked if removing a twin from the full dataset drastically changed the correlation between these two scores by comparing the correlation with and without the randomly removed twin, which it did not ($r=0.547$ vs 0.551).

**OSPAN and SYMSPAN correlations amongst twins only** (n=60)

| | Correlation | p |
|---|---|---|
| OSPAN | 0.634 | 0.000 |
| SYMSPAN | 0.451 | 0.012 |
| WMC COMPOSITE | 0.031 | 0.871 |

**OSPAN and SYMSPAN correlations (one twin from a pair dataset n=88)**

| | M | SD | Skew | Kurtosis | Correlation |
|---|---|---|---|---|---|
| OSPAN | 26.682 | 7.532 | -0.947 | 0.284 | 0.546*** |
| SYMSPAN | 17.886 | 4.672 | -0.025 | -0.403 | - |
| WMC compositeNONTWIN | 0.029 | 1.753 | -0.842 | 0.958 | - |

#### S5.2 Behavioral Replication: no twin

Next, we checked the behavioral replication results without a randomly removed twin. Hypothesis test: Is there a significant interaction between the Proactive Session and WMC <0?

The hypothesis is plotted as 2 probability distributions surrounding the estimated effect: light blue (prior estimate) and dark blue (posterior estimate). This visualizes how consistent and variable the effect is in the replication, compared to the prior analysis.

**Hypothesis Test: Data without twin pairs**
Is there a significant interaction between the Proactive Session and WMC <0? Additionally, the hypothesis is plotted as 2 probability distributions surrounding the estimated effect: light blue (prior estimate) and dark blue (posterior estimate). This visualizes how consistent and variable the effect is in the replication, compared to the prior analysis.

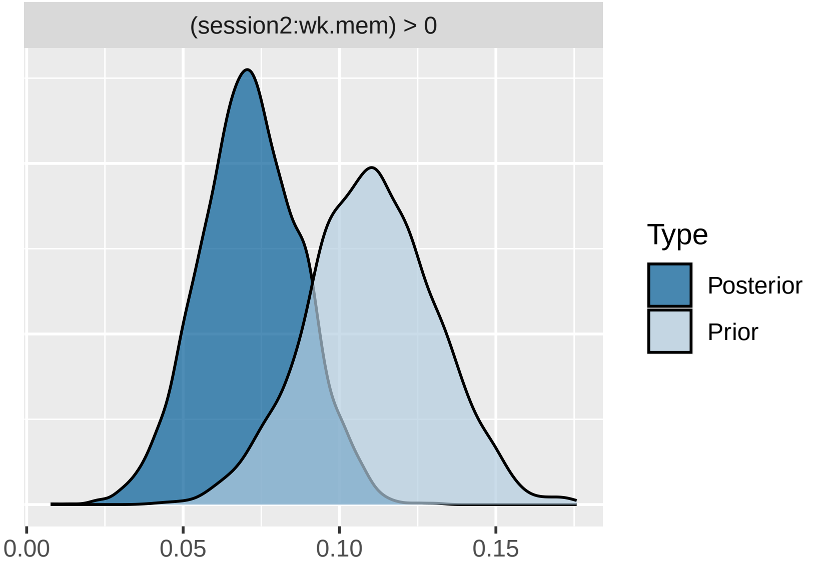


```
## Hypothesis Tests for class b:
##              Hypothesis Estimate Est.Error CI.Lower CI.Upper Evid.Ratio Post.Prob Star
## 1 (session2:wk.mem) > 0     0.07      0.02     0.05      0.1        Inf         1    *
## ---
## 'CI': 90%-CI for one-sided and 95%-CI for two-sided hypotheses.
## '*': For one-sided hypotheses, the posterior probability exceeds 95%;
## for two-sided hypotheses, the value tested against lies outside the 95%-CI.
## Posterior probabilities of point hypotheses assume equal prior probabilities.
```

| Fixed Effects | Estimate (SD) | 95% CI | Odds (SD) |
|---|---|---|---|
| Intercept | .284 (.036) | [.212, .353]* | 1.329 (.048) |
| Bas | -.254 (.024) | [-.3, -.205]* | .056 (.002) |
| Pro | .612 (.025) | [.564, .66]* | .776 (.019) |
| Rea | -.242 (.025) | [-.291, -.193]* | .785 (.02) |
| WMC | .085 (.029) | [.027, .142]* | .946 (.023) |
| Trial | -2.888 (.043) | [-2.971, -2.803]* | .947 (.018) |
| Bas:WMC | -.054 (.019) | [-.092, -.018] | .982 (.019) |
| Pro:WMC | .072 (.016) | [.041, .102]* | 1.845 (.046) |
| Rea:WMC | -.05 (.021) | [-.091, -.009]* | .952 (.02) |
| Bas:Trial | -.056 (.024) | [-.102, -.011]* | 1.491 (.027) |
| Pro:Trial | .399 (.018) | [.362, .435]* | 1.074 (.017) |
| Rea:Trial | -.06 (.025) | [-.108, -.013]* | .942 (.023) |
| WMC:Trial | .016 (.033) | [-.05, .08] | 1.009 (.016) |
| Bas:WMC:Trial | -.018 (.02) | [-.057, .02]* | 1.089 (.031) |
| Pro:WMC:Trial | .009 (.015) | [-.022, .038]* | 1.017 (.033) |
| Rea:WMC:Trial | -.034 (.021) | [-.074, .006]* | .967 (.02) |

#### S5.3 Trial Level A-cue Bias:

We were also curious if our A-cue bias metric was correlated between twins.Suprisingly, A-cue bias was not signifcantly correlated for the proactive session (r=0.13 n.s.) and only marginally correlated in the baseline (0.36, p=0.056).

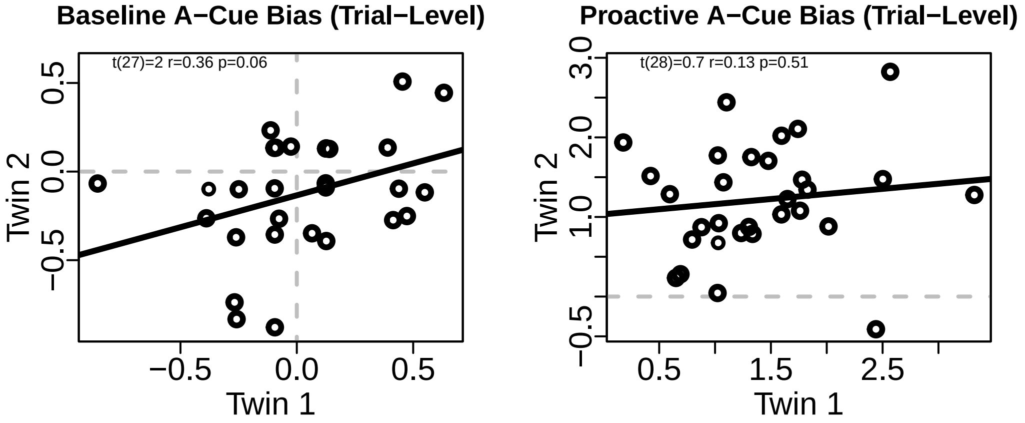


#### S5.3 Neural Activation:

Next we checked brain-behavior correlations between twins for the parcels selected for the main analyses (i.e. lMOT and rDLPFC)

Suprisingly twins were not correlated in their neural activation, but the N of 30 pairs is small for neuroimaging correlations.

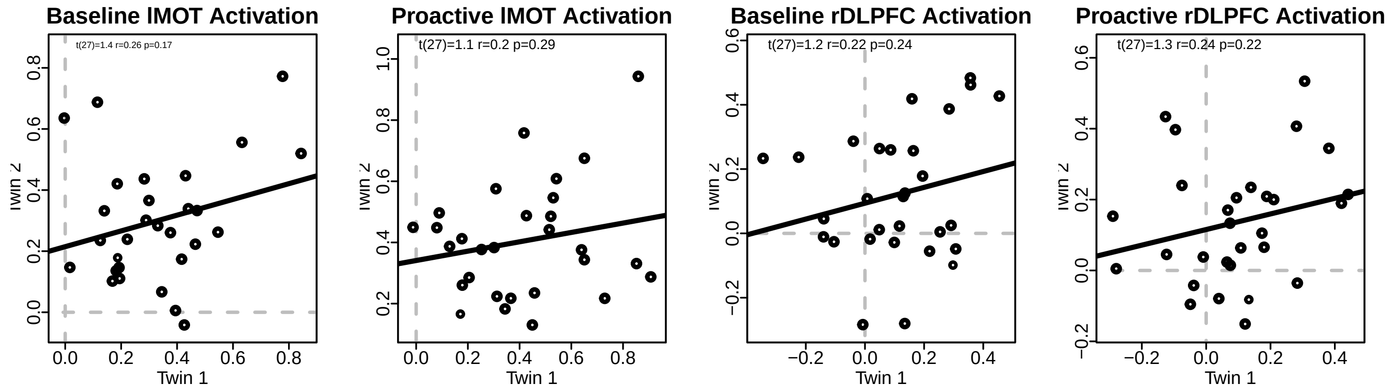


#### S5.4 Brain Behavior:

Next we checked brain-behavior correlations between twins for the parcels selected for the main analyses (i.e. lMOT and rDLPFC)

For lMOT and A-cue Bias, rDLPFC and WMC, and between lMOT and rDLPFC were significant for twins and for the full dataset after removing 1 twin at random.

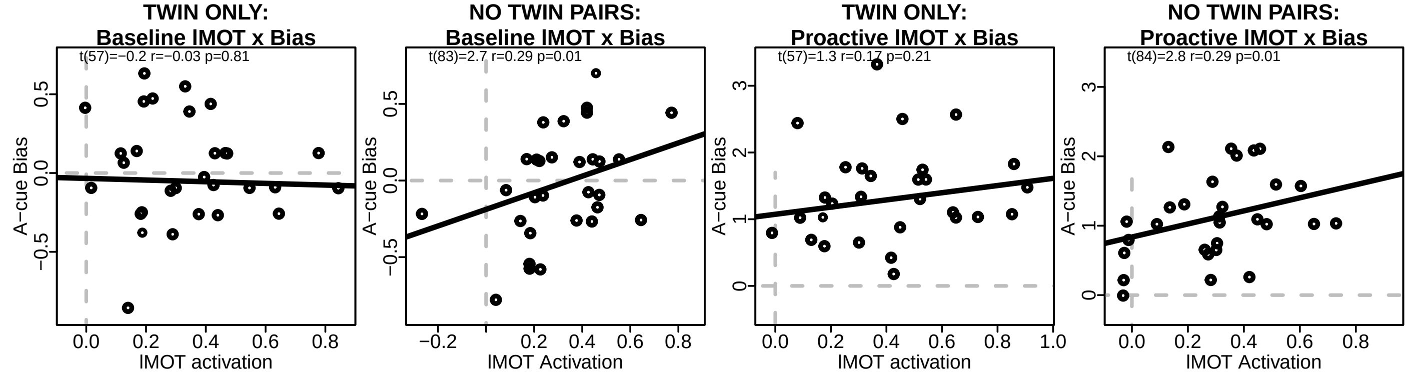

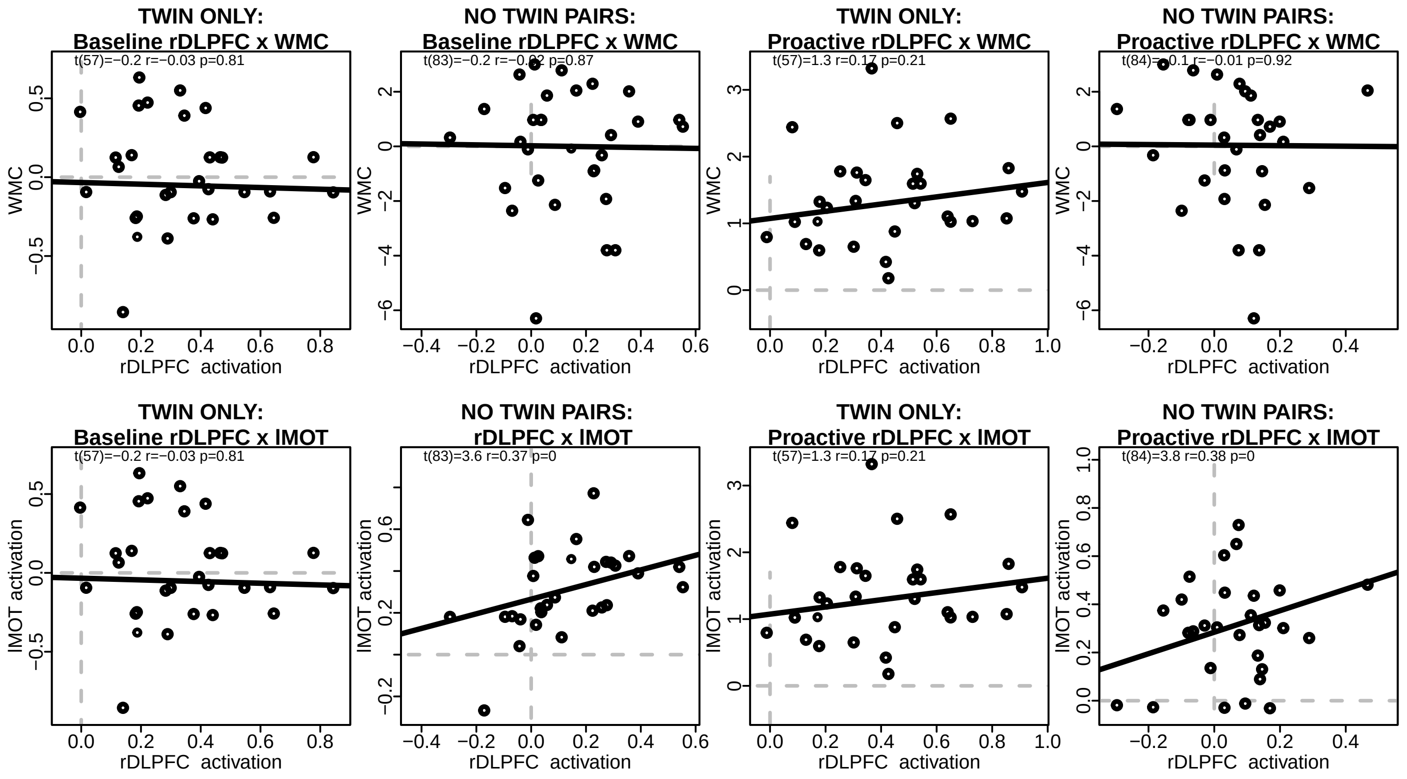


#### S5.5 Within Person Mediation:

Next, we reran our within person mediation testing if the change in A-cue behavioral bias from baseline to proactive was mediated by the increase in A-cue activation in the lMOT.
**after removing 1 twin results:**

| | Estimate | SD | T-value | p-value |
|---|---|---|---|---|
| (Intercept) | 1.164 | 0.052 | 22.302 | ¡.001 |
| H.sum.centered | -.179 | 0.137 | -1.310 | .192 |
| H.diff | .692 | 0.224 | 3.090 | .002 |

```
##
## Call:
## lm(formula = Y.diff ~ H.sum.centered + H.diff)
##
## Residuals:
##      Min       1Q   Median       3Q      Max
## -1.56462 -0.43578  0.00545  0.40678  1.69261
##
## Coefficients:
##                Estimate Std. Error t value Pr(>|t|)
## (Intercept)     1.16398    0.05219   22.30  < 2e-16 ***
## H.sum.centered -0.17922    0.13686   -1.31  0.19220
## H.diff          0.69243    0.22406    3.09  0.00235 **
## ---
## Signif. codes:  0 '***' 0.001 '**' 0.01 '*' 0.05 '.' 0.1 ' ' 1
##
## Residual standard error: 0.6717 on 163 degrees of freedom
##   (10 observations deleted due to missingness)
## Multiple R-squared:  0.06095, Adjusted R-squared:  0.04943
```

```
## F-statistic:  5.29 on 2 and 163 DF,  p-value: 0.005943
```

#### S5.6 Between Person Path Analysis:

We ran our between person path analysis with and without twin pairs.
Indirect results using proactive session values after partialling out shared variance from the baseline session. This model tests for an additive relationship between WMC-rDLPFC-lMOT-Acue bias. Rows with RED text denote a confidence interval that is all positive or negative; LHS: left hand side; RHS: right hand side; OP: operator; regression; (residual)(co)variance; 1 intercept; NOTE WMC is a static measure mediation path testing for (a) individuals with higher WMC have higher rDLPFC activity, (b) higher rDLPFC activity results in increased LMOT activation during A-cue presentation, and (c) which results in an increased likelihood to make a target response, and further, whether this pathway explains significant variance in (d) the direct path of WMC predicting A-cue bias.
**No Twin Pair dataset Zero-Order path analysis:**

| lhs | op | rhs | label | est | se | z | pvalue | ci.lower | ci.upper |
|---|---|---|---|---|---|---|---|---|---|
| pfc9 | | wmc | a | 0.001 | 0.125 | 0.005 | 0.996 | -0.235 | 0.255 |
| lMOT | | pfc9 | b | 0.325 | 0.091 | 3.563 | 0.000 | 0.141 | 0.498 |
| bias | | lMOT | c | 0.217 | 0.103 | 2.099 | 0.036 | 0.006 | 0.412 |
| bias | | wmc | d | 0.117 | 0.122 | 0.955 | 0.339 | -0.123 | 0.356 |
| pfc9 | | pfc9 | | 630.666 | 60.176 | 10.480 | 0.000 | 503.987 | 739.120 |
| lMOT | | lMOT | | 564.188 | 62.499 | 9.027 | 0.000 | 432.334 | 678.917 |
| bias | | bias | | 594.440 | 61.720 | 9.631 | 0.000 | 455.675 | 695.829 |
| wmc | | wmc | | 450.738 | 0.000 | NA | NA | 450.738 | 450.738 |
| pfc9 | 1 | | | 43.978 | 5.668 | 7.759 | 0.000 | 32.959 | 55.364 |
| lMOT | 1 | | | 29.715 | 4.790 | 6.203 | 0.000 | 20.548 | 39.383 |
| bias | 1 | | | 30.172 | 6.797 | 4.439 | 0.000 | 16.967 | 43.611 |
| wmc | 1 | | | 36.632 | 0.000 | NA | NA | 36.632 | 36.632 |
| abc | := | (a*b*c) | abc | 0.000 | 0.010 | 0.004 | 0.997 | -0.019 | 0.022 |
| total | := | d+(a*b*c) | total | 0.117 | 0.120 | 0.974 | 0.330 | -0.120 | 0.353 |

```
## lavaan 0.6-18 ended normally after 8 iterations
##
##   Estimator                                         ML
##   Optimization method                           NLMINB
##   Number of model parameters                        10
##
##                                                   Used       Total
##   Number of observations                            87          88
##
## Model Test User Model:
##
##   Test statistic                                 4.778
##   Degrees of freedom                                 2
##   P-value (Chi-square)                           0.092
##
## Model Test Baseline Model:
##
##   Test statistic                                19.616
##   Degrees of freedom                                 6
##   P-value                                        0.003
##
## User Model versus Baseline Model:
##
##   Comparative Fit Index (CFI)                    0.796
##   Tucker-Lewis Index (TLI)                       0.388
##
## Loglikelihood and Information Criteria:
```

```
##
##   Loglikelihood user model (H0)              -1204.229
##   Loglikelihood unrestricted model (H1)             NA
##
##   Akaike (AIC)                                2428.457
##   Bayesian (BIC)                              2453.116
##   Sample-size adjusted Bayesian (SABIC)       2421.563
##
## Root Mean Square Error of Approximation:
##
##   RMSEA                                          0.126
##   90 Percent confidence interval - lower         0.000
##   90 Percent confidence interval - upper         0.277
##   P-value H_0: RMSEA <= 0.050                    0.140
##   P-value H_0: RMSEA >= 0.080                    0.782
##
## Standardized Root Mean Square Residual:
##
##   SRMR                                           0.057
##
## Parameter Estimates:
##
##   Standard errors                            Bootstrap
##   Number of requested bootstrap draws             5000
##   Number of successful bootstrap draws            5000
##
## Regressions:
##                    Estimate  Std.Err  z-value  P(>|z|) ci.lower ci.upper   Std.lv  Std.all
##   pfc9 ~
##     wmc        (a)    0.001    0.125    0.005    0.996   -0.235    0.255    0.001    0.001
##   lMOT ~
##     pfc9       (b)    0.325    0.091    3.563    0.000    0.141    0.498    0.325    0.325
##   bias ~
##     lMOT       (c)    0.217    0.103    2.099    0.036    0.006    0.412    0.217    0.217
##     wmc        (d)    0.117    0.122    0.955    0.339   -0.123    0.356    0.117    0.099
##
## Intercepts:
##                    Estimate  Std.Err  z-value  P(>|z|) ci.lower ci.upper   Std.lv  Std.all
##    .pfc9             43.978    5.668    7.759    0.000   32.959   55.364   43.978    1.751
##    .lMOT             29.715    4.790    6.203    0.000   20.548   39.383   29.715    1.183
##    .bias             30.172    6.797    4.439    0.000   16.967   43.611   30.172    1.202
##
## Variances:
##                    Estimate  Std.Err  z-value  P(>|z|) ci.lower ci.upper   Std.lv  Std.all
##    .pfc9            630.666   60.176   10.480    0.000  503.987  739.120  630.666    1.000
##    .lMOT            564.188   62.499    9.027    0.000  432.334  678.917  564.188    0.895
##    .bias            594.440   61.720    9.631    0.000  455.675  695.829  594.440    0.943
##
## R-Square:
##                    Estimate
##     pfc9              0.000
##     lMOT              0.105
##     bias              0.057
##
## Defined Parameters:
##                    Estimate  Std.Err  z-value  P(>|z|) ci.lower ci.upper   Std.lv  Std.all
##     abc               0.000    0.010    0.004    0.997   -0.019    0.022    0.000    0.000
##     total             0.117    0.120    0.974    0.330   -0.120    0.353    0.117    0.099
```

**No Twin Pair dataset Proactive   Baseline path analysis:**

| lhs | op | rhs | label | est | se | z | pvalue | ci.lower | ci.upper |
|---|---|---|---|---|---|---|---|---|---|
| pfc9 | | wmc | a | -0.016 | 0.009 | -1.799 | 0.072 | -0.033 | 0.003 |
| lMOT | | pfc9 | b | 0.335 | 0.098 | 3.419 | 0.001 | 0.154 | 0.537 |
| bias | | lMOT | c | 1.004 | 0.493 | 2.037 | 0.042 | 0.054 | 1.992 |
| bias | | wmc | d | 0.038 | 0.044 | 0.854 | 0.393 | -0.048 | 0.126 |
| pfc9 | | pfc9 | | 0.025 | 0.004 | 5.901 | 0.000 | 0.017 | 0.034 |
| lMOT | | lMOT | | 0.023 | 0.004 | 6.090 | 0.000 | 0.015 | 0.030 |
| bias | | bias | | 0.461 | 0.069 | 6.712 | 0.000 | 0.319 | 0.588 |
| wmc | | wmc | | 2.796 | 0.000 | NA | NA | 2.796 | 2.796 |
| pfc9 | 1 | | | -0.011 | 0.017 | -0.623 | 0.533 | -0.045 | 0.024 |
| lMOT | 1 | | | 0.004 | 0.017 | 0.227 | 0.820 | -0.029 | 0.036 |
| bias | 1 | | | 0.004 | 0.076 | 0.053 | 0.958 | -0.142 | 0.158 |
| wmc | 1 | | | -0.013 | 0.000 | NA | NA | -0.013 | -0.013 |
| abc | := | (a*b*c) | abc | -0.005 | 0.005 | -1.140 | 0.254 | -0.017 | 0.001 |
| total | := | d+(a*b*c) | total | 0.033 | 0.044 | 0.743 | 0.458 | -0.052 | 0.120 |

```
## lavaan 0.6-18 ended normally after 1 iteration
##
##   Estimator                                         ML
##   Optimization method                           NLMINB
##   Number of model parameters                        10
##
##                                                   Used       Total
##   Number of observations                            84          88
##
## Model Test User Model:
##
##   Test statistic                                 1.901
##   Degrees of freedom                                 2
##   P-value (Chi-square)                           0.387
##
## Model Test Baseline Model:
##
##   Test statistic                                19.482
##   Degrees of freedom                                 6
##   P-value                                        0.003
##
## User Model versus Baseline Model:
##
##   Comparative Fit Index (CFI)                    1.000
##   Tucker-Lewis Index (TLI)                       1.022
##
## Loglikelihood and Information Criteria:
##
##   Loglikelihood user model (H0)                -11.537
##   Loglikelihood unrestricted model (H1)        -10.587
##
##   Akaike (AIC)                                  43.075
##   Bayesian (BIC)                                67.383
##   Sample-size adjusted Bayesian (SABIC)         35.838
##
## Root Mean Square Error of Approximation:
##
##   RMSEA                                          0.000
##   90 Percent confidence interval - lower         0.000
##   90 Percent confidence interval - upper         0.213
##   P-value H_0: RMSEA <= 0.050                    0.460
##   P-value H_0: RMSEA >= 0.080                    0.442
##
```

```
## Standardized Root Mean Square Residual:
##
##   SRMR                                           0.036
##
## Parameter Estimates:
##
##   Standard errors                            Bootstrap
##   Number of requested bootstrap draws             5000
##   Number of successful bootstrap draws            5000
##
## Regressions:
##                    Estimate  Std.Err  z-value  P(>|z|) ci.lower ci.upper   Std.lv  Std.all
##   pfc9 ~
##     wmc        (a)   -0.016    0.009   -1.799    0.072   -0.033    0.003   -0.016   -0.162
##   lMOT ~
##     pfc9       (b)    0.335    0.098    3.419    0.001    0.154    0.537    0.335    0.339
##   bias ~
##     lMOT       (c)    1.004    0.493    2.037    0.042    0.054    1.992    1.004    0.229
##     wmc        (d)    0.038    0.044    0.854    0.393   -0.048    0.126    0.038    0.091
##
## Intercepts:
##                    Estimate  Std.Err  z-value  P(>|z|) ci.lower ci.upper   Std.lv  Std.all
##    .pfc9             -0.011    0.017   -0.623    0.533   -0.045    0.024   -0.011   -0.067
##    .lMOT              0.004    0.017    0.227    0.820   -0.029    0.036    0.004    0.024
##    .bias              0.004    0.076    0.053    0.958   -0.142    0.158    0.004    0.006
##
## Variances:
##                    Estimate  Std.Err  z-value  P(>|z|) ci.lower ci.upper   Std.lv  Std.all
##    .pfc9              0.025    0.004    5.901    0.000    0.017    0.034    0.025    0.974
##    .lMOT              0.023    0.004    6.090    0.000    0.015    0.030    0.023    0.885
##    .bias              0.461    0.069    6.712    0.000    0.319    0.588    0.461    0.942
##
## R-Square:
##                    Estimate
##     pfc9              0.026
##     lMOT              0.115
##     bias              0.058
##
## Defined Parameters:
##                    Estimate  Std.Err  z-value  P(>|z|) ci.lower ci.upper   Std.lv  Std.all
##     abc              -0.005    0.005   -1.140    0.254   -0.017    0.001   -0.005   -0.013
##     total             0.033    0.044    0.743    0.458   -0.052    0.120    0.033    0.078
```
